# Towards prevention of re-entrant arrhythmias: Injectable hydrogel electrodes enable direct capture of previously inaccessible cardiac tissue

**DOI:** 10.1101/2021.11.03.467102

**Authors:** Gabriel J. Rodriguez-Rivera, Allison Post, Mathews John, Skylar Buchan, Megan Wancura, Malgorzata Chwatko, Christina Waldron, Abbey Nkansah, Drew Bernard, Nikhith Kalkunte, Janet Zoldan, Mathieu Arseneault, Mehdi Razavi, Elizabeth Cosgriff-Hernandez

**Author notes:** Department of Chemical and Materials Engineering; University of Kentucky, Lexington, KY 40506, USA. OmegaChem Inc.; Quebec G6W 7V6, Canada (Mathieu Arseneault). Corresponding authors: Elizabeth Cosgriff-Hernandez; Mehdi Razavi.

## Abstract

Re-entrant arrhythmias—the leading cause of sudden cardiac death—are caused by diseased myocardial tissue and consequent delayed myocardial conduction. Access to the coronary veins that cross the “culprit” scar regions where re-entry originates can provide improved pacing to these delayed regions, offering a novel opportunity to prevent ventricular arrhythmias and circumvent the need for painful defibrillation, risky cardiac ablation, or toxic and often ineffective antiarrhythmic medications. However, there are no pacing electrodes which are small or focal enough to navigate these tributaries. To address this need, we have developed an injectable conductive hydrogel that can fill the epicardial coronary veins and their mid-myocardial tributaries. When connected to a standard pacing lead, these injected hydrogels can be converted into flexible electrodes that directly pace the previously inaccessible mid-myocardial tissue. In our two-component system, hydrogel precursor solutions can be injected through a dual lumen catheter in a minimally invasive deployment strategy to provide direct access to the diseased regions with precision and ease. Mixing of the two solutions upon injection into the vein activates redox-initiated crosslinking of the gel for rapid *in situ* cure without an external stimulus. An *ex vivo* porcine model was used to identify the requisite viscosity and cure rate for gel retention and homogeneity. Ionic species added to the hydrogel precursor solutions conferred conductivity above target myocardium values that was retained after implantation. Successful *in vivo* deployment demonstrated that the hydrogel electrode filled the anterior interventricular vein with extension into the septal (mid-myocardial) venous tributaries to depths far more distal and refined than any current technologies allow. In addition to successful capture and pacing of the heart, analysis of surface ECG tracings revealed a novel pacing observation highly specific for and suggestive of capture of extensive swaths of septal myocardial tissue. This is the first report of an injectable electrode used to successfully pace the mid-myocardium and mimic physiologic conduction. Furthermore, *in vivo* cardiac electroanatomical mapping studies in an ablation scar model showed uniform capture along the hydrogel in the vessels as well as increased capture area compared to point pacing. Collectively, these findings demonstrate that this injectable hydrogel electrode can be deployed to scarred regions of the heart to provide a reliable pacing modality that most closely resembles native conduction with the potential to eliminate delayed myocardial conduction and associated re-entry.

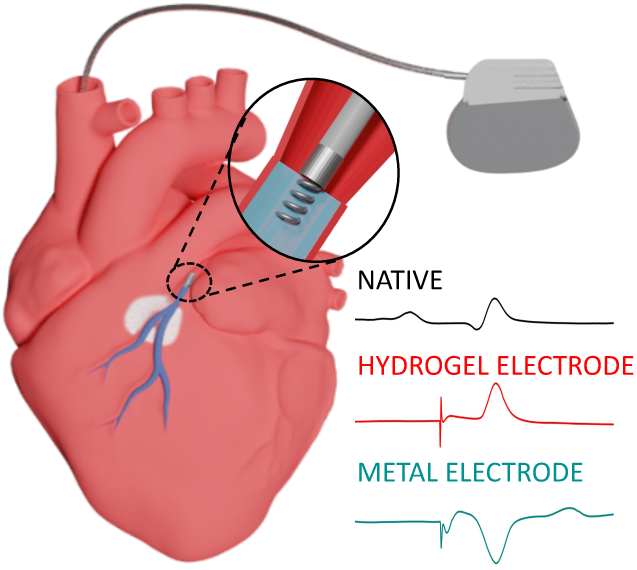

**One Sentence Summary:** Injectable hydrogel electrodes achieve pacing that mimics physiologic conduction by capturing midmyocardial tissue

## INTRODUCTION

Ventricular arrhythmias (VA) are by far the leading cause of sudden cardiac death in the United States. Current pacing modalities are insufficient to prevent or treat these (*1*). Scarred or otherwise diseased myocardium leads to slowed electrical conduction of cardiac tissue. The attendant spatial and temporal heterogeneities cause wave break, re-entry, and chaotic cardiac activity and fibrillation (*2–5*). Much like chemotherapy, current approaches to treat are inherently primitive: Current management relies on a combination of antiarrhythmic drugs – all with high toxicity profiles – and/or ablation and destruction of tissues adjacent or in the vicinity of the diseased regions. Antiarrhythmic drugs work by further slowing conduction velocity in an effort towards complete elimination of such conduction (*6*). This renders them pro-arrhythmic for exactly the same reason: decreased conduction’s requisite role in the initiation and sustenance of re-entry (*7*). Although widely adopted, ablative strategies have a high failure rate with recurrent arrhythmias in 18 - 40% of cases (*8*). Of paramount importance, none of these technologies correct re-entry’s major underpinning: delayed conduction. For many patients, the only option is an implantable cardiac defibrillator that uses high energy shocks to extinguish re-entry. These far exceed the pain threshold and have a well-documented negative impact on quality of life, including post-traumatic stress disorder and depression (*9*). A preventative treatment regimen addressing its underlying pathophysiology remains lacking.

In this work, we developed an injectable hydrogel electrode that fills the epicardial coronary veins and tributaries, converting them into flexible electrodes that can finally reach heretofore inaccessible mid-myocardium (Figure 1). Transforming these tributaries into flexible electrodes enables simultaneous pacing from multiple sites along the length of the electrode rather than froma single point stimulus. We hypothesize that the resultant planar wave propagation from the length of the electrode will stimulate wide areas of ventricular tissue that would have been otherwise been subject to delayed activation. This normalizes and eliminates the regions of delayed activation that are the underpinning of re-entry. To this end, we have developed a unique hydrogel system that has the requisite conductivity, biostability, and rapid *in situ* cure to be delivered via a transvenous catheter. First, a new polyethylene glycol-based macromer with improved hydrolytic stability and durability was designed and synthesized. Redox chemistry was used to provide rapid *in situ* cure without an external stimulus and an *ex vivo* porcine model was used to identify the requisite viscosity and cure rate for gel retention and homogeneity. Ionic species added to the hydrogel precursor solutions conferred conductivity above target myocardium values and the stability of the gel conductivity was assessed *in vitro* and *in vivo*. A porcine model was then used to confirm intravital delivery and successful pacing via the hydrogel electrode. We then showed that these electrodes maintain robust conduction properties in the setting of myocardial scar using a porcine cardiac ablation model. Finally, we defined the subacute safety profile of this technique in a porcine model using a combination of cardiac function, cardiac enzymes, and histology. Collectively, this work demonstrates for the first time the ability to confer direct electrical stimulation of the native and scarred mid-myocardium using an injectable hydrogel electrode as a new pacing modality.

**Figure 1.**
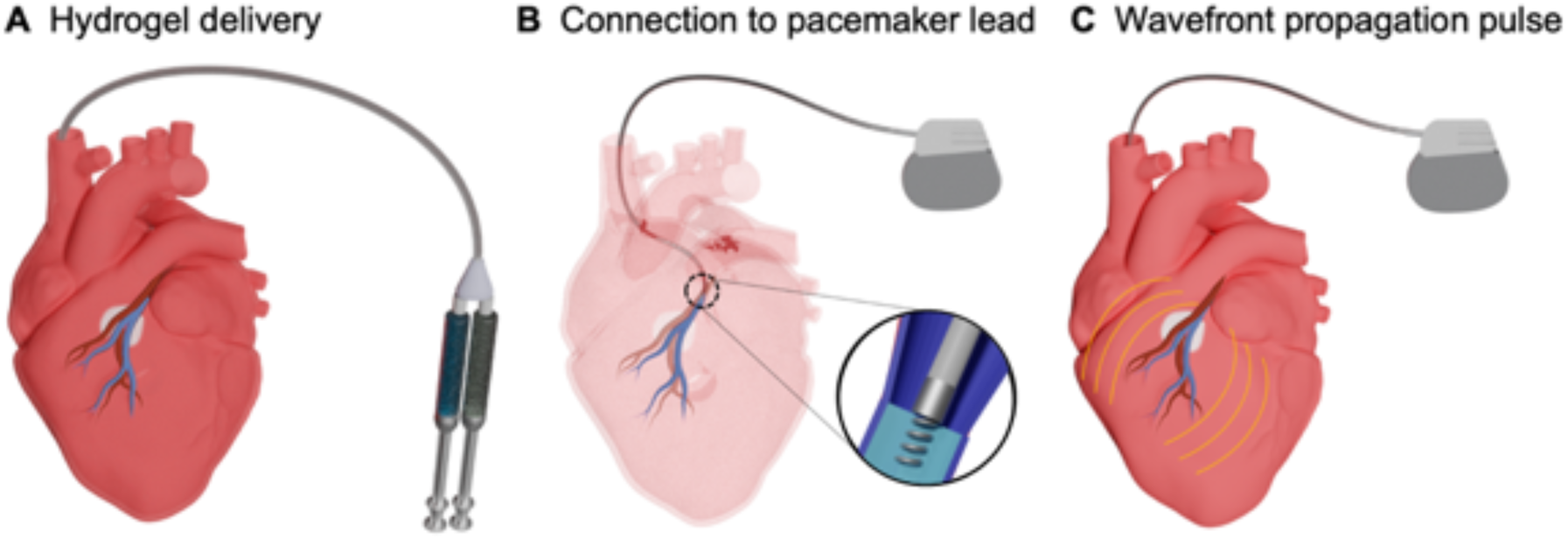
Schematic of injectable hydrogel system to transform coronary veins into flexible electrodes that capture inaccessible cardiac tissue. **(A)** Hydrogel precursors solutions are delivered using a dual lumen catheter to fill coronary veins and tributaries that span the myocardium near the scarred tissue. Upon mixing of the two hydrogel solutions in the coronary veins, redox-initiated crosslinking results in rapid cure of the ionic hydrogel. **(B)** Subsequent connection to a pacemaker lead to the ionic hydrogel electrode provides increased tissue contact across the myocardium. **(C)** Wavefront activation of the myocardium along the hydrogel electrode reduces the energy required for defibrillation.

## RESULTS

### Injectable hydrogels with rapid *in situ* cure

There are no current hydrogels that can provide the multifaceted needs of a cardiac electrode with endovascular delivery to the midmyocardium: injectability, conductivity, and biostability. Poly(ethylene glycol) (PEG) hydrogels are a popular choice based on their established biocompatibility and highly tunable, soft tissue-like properties; however, traditional acrylate- derivatized PEG hydrogels are susceptible to slow degradation *in vivo* due to hydrolysis. To generate a PEG-based hydrogel suitable for long-term implantable applications, we synthesized a new hydrogel chemistry that combines biostability with flexibility and durability. Poly(ether urethane diacrylamide) (PEUDAm) contains urethane and amide groups that are resistant to hydrolysis at physiological conditions. The detailed synthetic route is described in **Figure S1** and successful conversion (>95%) was confirmed with H^1^ NMR (**Figure S2-S4**). The resulting PEUDAm macromer displayed a number average molecular weight of 19,960 ± 40 g/mol and a narrow polydispersity index of 1.08 ± 0.01. A small degree of polymer coupling was indicated by a secondary peak at Mn = 49,340 ± 220 g/mol (**Figure S5**). In addition to the biostable PEG macromer, we synthesized N-acryloyl glycinamide (NAGA), a small molecule crosslinker with bidentate hydrogen bonding (*10*). Successful acrylation and purity of NAGA were confirmed by H^1^ NMR analysis (**Figure S6**). The resulting hydrogel (20kDa, 20wt% PEUDAm+ 1% NAGA) properties are summarized in **Table 1**. The hydrogel stiffness, 59.1 ± 7.5 kPa, was within the range of myocardial tissue, reported as 20 to 200 kPa by Ramadan et al. for ventricles (*11*) or 20 to 500 kPa reported from mathematical models. The hydrogel ultimate elongation of 111 ± 28% was above the average myocardium strains reported from imaging analysis ranging from 10-55% (*12*) and above the axial strain of cardiac veins, which is up to 30% strain for the larger veins (*13*). In addition to tissue-matching mechanical properties, accelerated hydrolytic degradation studies indicated that the new hydrogel was biostable, **Figure S7**.

The proposed design also requires rapid *in situ* cure of the hydrogel without external stimuli (e.g UV). Our lab has previously shown tunable gelation using the redox pair APS and IG (*14*). In this study, we demonstrated faster gelation kinetics than previously described by increasing APS concentration at constant APS: IG of 1:2 molar ratio. A double-barrel syringe with mixing head was used to deliver the precursor solution as an analog for future delivery from a dual lumen catheter, **Figure 2A**. The effect of compositional variables on network formation was investigated by monitoring changes in storage and loss modulus with rheology. The crossover point was achieved too quickly for data quantification for most samples, **Figure S8**. Complete network formation was determined as the third point, after which there was a less than 1.0% change in modulus. Iterative testing was used to identify a target cure rate of approximately 2 minutes which corresponded to an APS concentrations ≥0.75 mM (**Figure 2B**). The inclusion of 1 w/w% NAGA increased the cure rate and gel fraction (**Figure S9**, **Table S1**). We hypothesized that this was due to the increase in acrylamide groups and mobility of the smaller molecules. In the presence of blood, full network formation was delayed from 1.7 to 5 minutes (**Figure S9**).

**Table 1.** Physical properties of the hydrogel electrode. : PEUDAm 20kDa, 20% + 1% NAGA, 0.75 mM APS/1.5 mM IG. Average of 3 independent hydrogels (n=3).

| Physical Property | Average value |
| --- | --- |
| Gel fraction | $95.2 \pm 1.4 \%$ |
| Mass swelling ratio | $17.4 \pm 0.1 \text{ g/g}$ |
| Young's modulus | $59.1 \pm 7.5 \text{ kPa}$ |
| Ultimate tensile strength | $50.6 \pm 13.3 \text{ kPa}$ |
| Ultimate elongation | $111 \pm 28 \%$ |

To determine the suitability of this hydrogel composition for *in vivo* assessment, the cytocompatibility of the *in situ* cure mechanism was examined prior to *in vivo* deployment. In the target application, the hydrogels will be cured in direct contact with the endothelial cells on the cardiac veins with potential diffusion to the myocardium. To assess *in vitro* cytocompatibility, the potential cytotoxic effects of hydrogel leachables were evaluated on human umbilical vein endothelial cells (HUVECs) and human-induced pluripotent stem cells – cardiomyocytes (hiPSC-MCs). **Figure 2C** shows that cell viability 24 hours after indirect exposure to a freshly prepared hydrogels (20% 20kDa PEUDAm, 1 w/w% NAGA, 1mM APS/2mM IG) was tested on two different cell lines. Cell viability after hydrogel exposure was above 90% and comparable to positive controls (**Figure S10**). No adverse effect from indirect contact to the hydrogels immediately after gelation was observed indicating overall cytocompatibility of the redox-initiated hydrogel.

An initial biocompatibility assessment of the hydrogel was performed using an *in vivo* subcutaneous rat model. Pre-cured hydrogel and silicone discs were implanted to compare the host response to the hydrogel to a clinical control. The hydrogel and silicone disc implants elicited comparable, normal wound healing responses after 1, 2 and 4 weeks (**Figure S11**). At 1-week post-implant, the interface between the discs and surrounding tissue was composed mostly of macrophages, fibroblasts and capillaries, indicated by the red staining within a background of incipient extracellular matrix deposition. By 4 weeks, the hydrogel was surrounded with a 68.1 ± 21.8 μm thick fibrous capsule rich in collagen (as indicated by the blue staining, **Figure 2D**), which was comparable to the 83.9 ± 28.5 μm thick layer observed on the silicone control (**Table S2**, **Figure S11**). A hydrogel was also injected subcutaneously to further assess the impact of the curing process in vivo. The injected, uncured hydrogel elicited similar wound healing response in the subcutaneous tissue as that of implanted pre-cured hydrogel discs (**Figure 2D**, **Figure S11**, **Table S3**). This data indicated the requisite biocompatibility of the *in situ*-curing hydrogel formulation to proceed with preclinical testing in a large animal model to assess the safety and efficacy of this approach.

### Precursor solution viscosity and cure rate determine hydrogel retention in an *ex vivo* model

An *ex vivo* porcine heart model was used to test hydrogel retention and homogeneity after injection. First, an *in vitro* method developed to visualize polymer mixing during delivery and gelation (**Movies S1-S2**) indicated that poor mixing strongly correlated to low gel fractions. This was attributed to insufficient initiation due to a lack of radical formation that is dependent on the reducing agent and initiator reaction. A mixing head at the end of the double-barrel syringe ensured adequate mixing upon delivery and high gel fractions. Once suitable mixing was confirmed, the effect of precursor solution viscosity and cure rate on hydrogel retention in the cardiac vein was studied. Hydrogel compositions with lower viscosity precursor solutions (∼42 mPa-s) and slower cure rates (∼5.8 min) did not form contiguous gels. Increasing the viscosity (∼83 mPa-s) by increasing the precursor macromer concentration and increasing the cure rate (∼2.2 min) with higher initiator concentrations resulted in contiguous gel formation, **Figure 3A**. The homogeneity across the hydrogel length after cure was studied *in vitro* in tubing and after dissection from the vein in the *ex vivo* studies. We also evaluated the potential impact of mixing with blood *in vitro* by pre-filling the tubing with porcine blood. Hydrogel swelling ratios were consistently higher in the *ex vivo* system or when combined with blood (**Figure 3B**). Differences in gel fractions between the *in vitro* gel cured in the empty tube and ex vivo studies were not significant. Swelling ratios were not significantly different from the proximal to the distal end indicating that the injectable gel was homogenous across its length.

**Figure 2.**
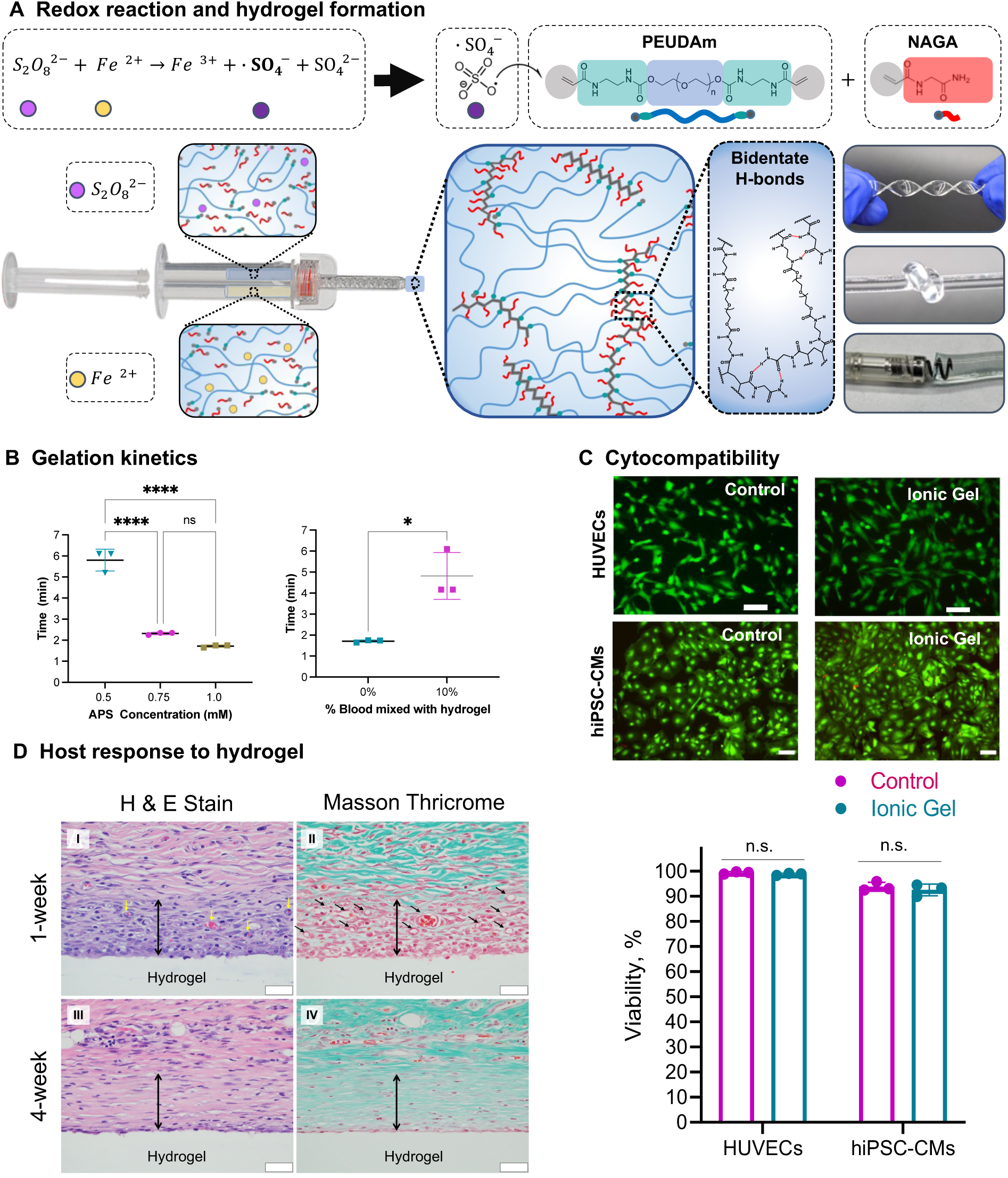
In situ cure kinetics and cytocompatibility of redox hydrogel. (**A)** Redox initiation reaction of new PEUDAm macromer + NAGA delivered using double barrel syringe with a mixing head. Resulting hydrogels display bidentate hydrogen bonding at netpoints for improved durability. **(B)** Effect of redox concentration (APS:IG ratio 1:2) on hydrogel cure rate (time to complete network formation, n=3). (**C**) Representative images of HUVECs and hiPSC-cardiomyocytes exposed to ionic hydrogels shortly after cure (n=3). Scale bar is 100 μm. (**D**) Representative histology images (H&E, Masson trichrome staining) showing the host response and development of a thin fibrous cap around the implanted hydrogel after 1 and 4 weeks. Note the diminishing cellularity (red staining) and increased collagen deposition (blue staining) around the implants over time. Scale bar is 40 μm. Individual data points, means, and standard deviation are presented (n = 3). Significant differences between groups are marked as follows: * (p < 0.05), **** (p < 0.0001).

**Figure 3.**
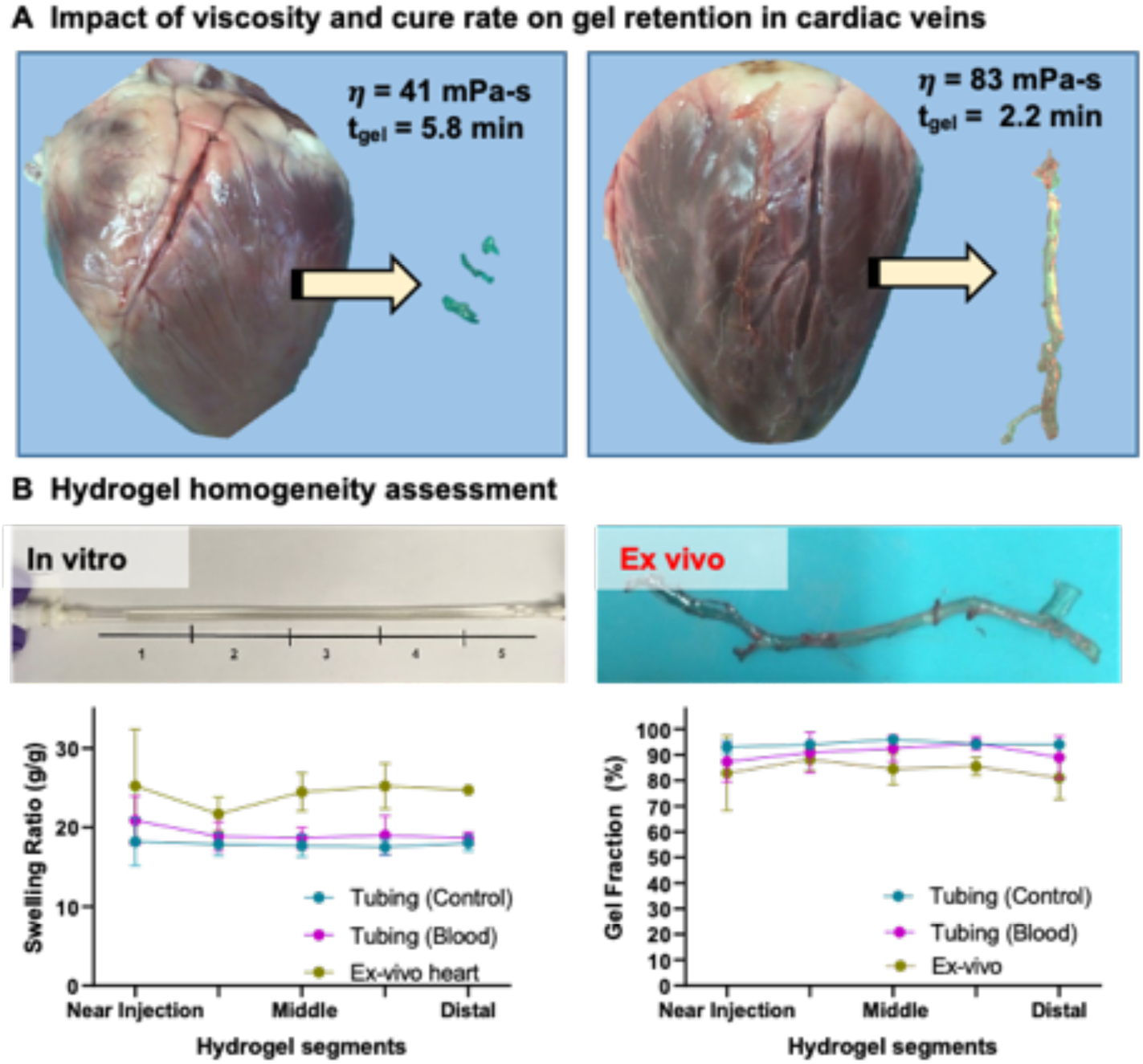
Hydrogel retention and homogeneity assessment in an *ex vivo* model. **(A)** Delivery of the hydrogel to main cardiac vein using an ex vivo porcine model. Poor gel retention was observed with lower viscosity precursor solution and slower cure rates. **(B)** Hydrogel segmental homogeneity assessment (mass swelling fraction, gel fraction) after cure in tubing, tubing filled with blood, and ex vivo porcine model (n = 3).

Flow simulations and rheological testing were used to further investigate the mechanistic role of these variables on hydrogel retention. When using the mixing head, there was minimal effect of precursor solution viscosity on the mixing of the two solutions upon delivery, **Figure 4**. Given that mixing of the two components is directly related to the rate of initiator decomposition and crosslinking, this flow simulation indicated that the mixing head negated possible effects on the cure rate. Rheology indicated that the complex viscosity of the solution after delivery was initially dependent on the polymer concentration and then increased over time as radical crosslinking was initiated until the gel point. Increasing the initiator concentration increased the rate of crosslinking with a corollary effect on the rate of viscosity build up. We hypothesized that poor hydrogel retention was due to venous draining of the precursor solution through the tributaries. To achieve sufficient hydrogel retention to form a contiguous hydrogel electrode in the vein, the system should be designed with sufficient precursor solution viscosity and a rapid cure to slow the dispersal prior to gelation, **Figure 4**. Gelation kinetics that were too rapid were found to clog the dispensing catheter. The hydrogel precursor formulation that balanced these parameters was 20 w% 20kDa PEUDAm with 1 wt% NAGA in saline solution at 0.75 mM APS and 1.5 mM IG. This composition resulted in sufficient hydrogel retention and homogeneity in the *ex vivo* model to support its evaluation in subsequent *in vivo* studies.

**Figure 4.**
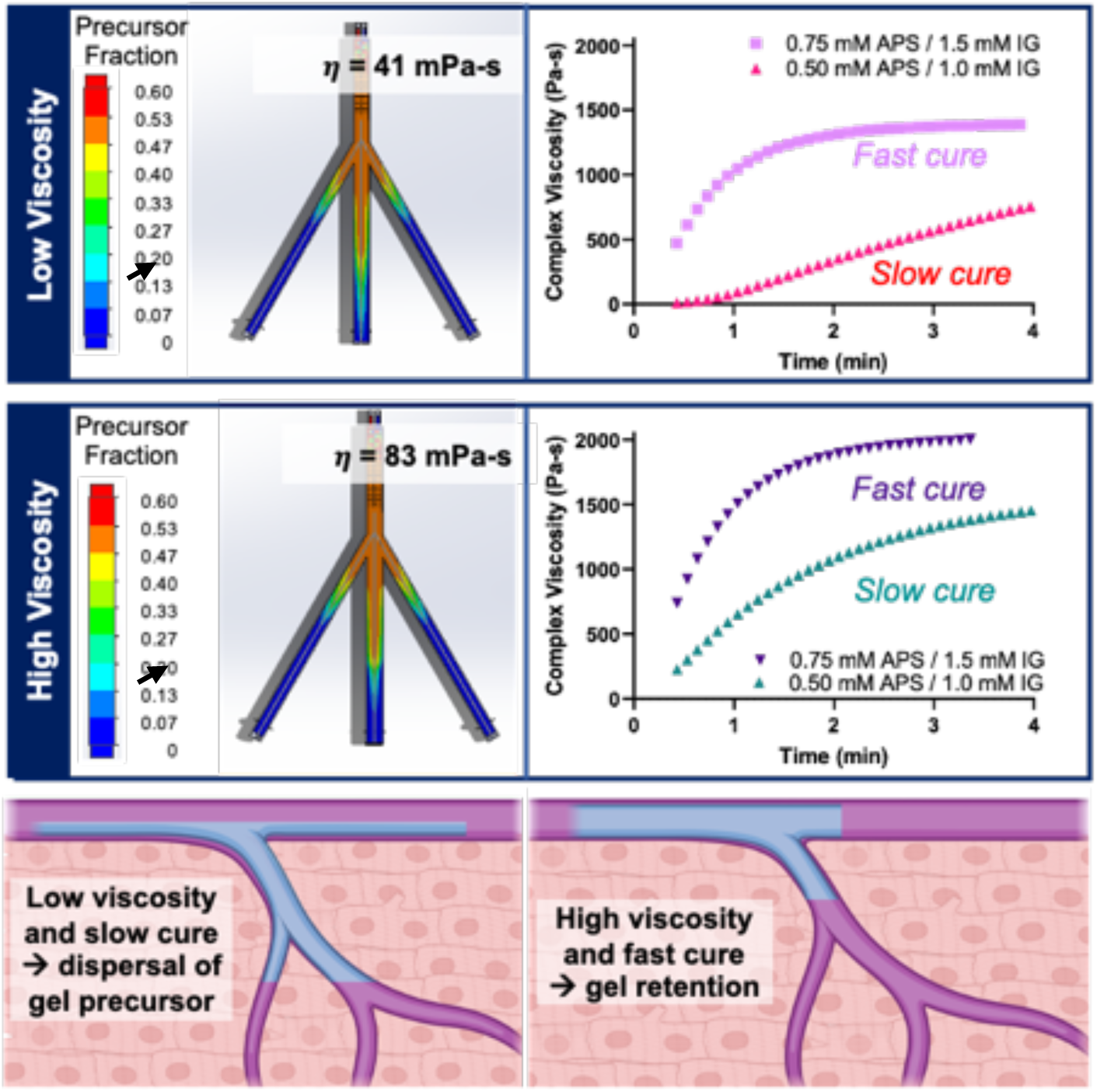
Role of mixing and *in situ* cure on hydrogel retention. Fluid simulations show adequate mixing and negligible differences in the volume fraction of the precursor solutions at both high and low viscosities. Time sweep indicates that high-viscosity precursor solutions with fast cure (higher initiator concentration) display a rapid increase in complex viscosity that was correlated with increased gel retention.

### Ionic hydrogels display stable conductivity higher than native myocardium

The ionic PEUDAm hydrogels rely on salts dissolved in the precursor solution to serve as charge carriers to generate a current when connected to a power source. As a basic proof-of-principle, **Figure 5** demonstrates that a current could be generated by connecting the ionic hydrogel to a battery and a light-emitting diode. Initially phosphate buffered saline was used to provide the ionic species (**Figure S12**); however, it was found that the phosphate led to iron precipitation which negatively impacted the redox-mediated initiation mechanism. To address this limitation, saline was used as an alternative source of ions in the hydrogel precursor solutions, **Figure S13**. In order to activate the myocardium beyond a single point of contact *in vivo*, the conductivity of the hydrogel must be above the myocardial conductivity that ranges from 0.1 to 6.0 mS/cm.(*15–17*) Initial conductivity performed with a four-point probe indicated that the ionic hydrogel more than doubled these target values (12.8 ± 1.5 mS/cm, **Figure S12**).

**Figure 5.**
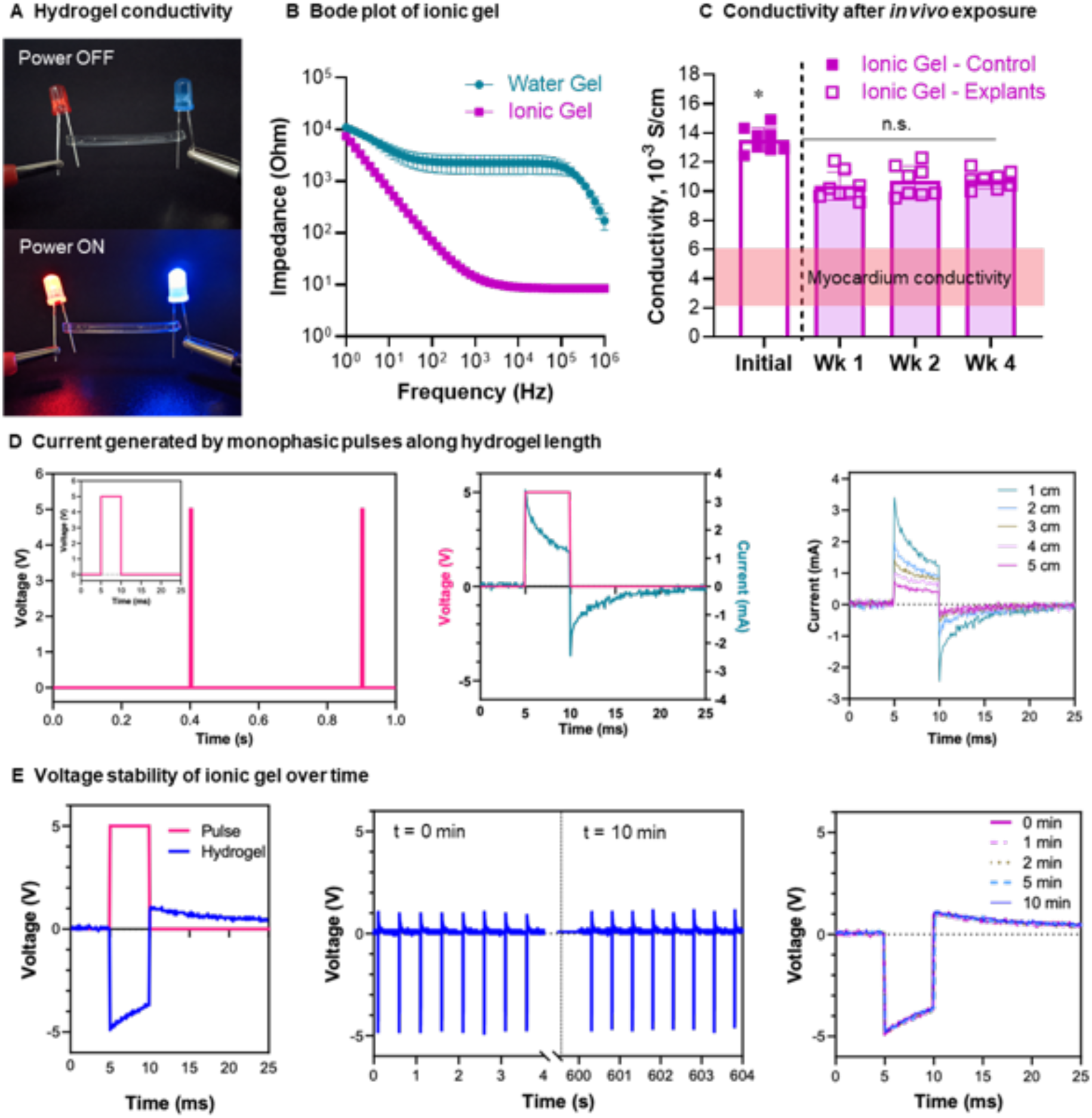
Conductivity of ionic hydrogel. **(A)** Demonstration of hydrogel conductivity after application of direct current. (**B)** Bode plot from EIS of hydrogel after gelation and equilibration in DI water (water gel) and PBS (ionic gel). (**C)** Effect of subcutaneous implantation on conductivity as calculated from EIS data, area highlighted in red represent the ranges of myocardium conductivity. Individual data points, mean, and standard deviation are presented (n = 8). Significant differences between groups are marked with **** (p < 0.0001). Significant differences between groups are marked with * (p < 0.05). **(D)** Current generated at different hydrogel lengths after 5V pulses. Pulses were 5 ms width at a frequency of 2 Hz (equivalent to 120 beat per minute). As hydrogel length increased, the increase in impedance caused a decrease in the current generated. (**E)** Hydrogel voltage response to stimuli remains stable after multiple pulses.

Electrochemical impedance spectroscopy was used to assess the ionic gel resistance and capacitance response to an applied voltage, **Figure 5B**. The slope on the Bode plots show the effects of ions transport and double-layer capacitance resulting in an increase in impedance at lower frequencies. Significantly, the ion concentration of the ionic gel resulted in reduced overall impedance of this gel relative to the water gel. The conductivity as measured by the impedance in the plateau regions was 13.5 ± 0.8 mS/cm for the ionic gel as compared to <0.1 mS/cm for the water gel control. Given that the conductivity was dependent on ion concentration in these gels, we hypothesized that there would be rapid ion diffusion in the hydrogels after implantation with corollary effects on gel conductivity. To determine the effect of ion flux on gel conductivity, and long-term ionic equilibrium, ionic gels were implanted subcutaneously into rats and the effect on conductivity measured after 1, 2 and 4 weeks. Gels reached and maintained a conductivity value of approximately 11 mS/cm (**Figure 5C**), which supports that the ion equilibration is higher than myocardial conductivity. Moreover, there were no significant changes in conductivity and gel properties after the first week, supporting stable hydrogel conduction. The swelling ratio and modulus of the gels were not significantly different before and after implantation (**Figure S14**), demonstrating network properties did not change significantly during one month, further supporting initial hydrogel biostability.

In the injectable hydrogel system, the pacing lead is connected to the proximal end of the hydrogel extending into the vein reaching the midmyocardium away from the power source. To assess how the current varies with hydrogel length, hydrogels were subjected to monophasic pulses of 5 milliseconds at a frequency of 2Hz, correlating to a pacing of 120 beats per minute (**Figure 5D**). A 5V power supply was used to function as a circuit that was comparable to standard pacemakers (*18*) (**Figure S15**). As expected, due to increased hydrogel impedance with length, the current decreased from 3.4 to 0.7 mA as the length increases from 1 to 5 cm (**Figure 5D**). These studies indicate that the hydrogel electrode was able to provide stable electrical stimuli over many cycles and across a substantial length of the cardiac vein (**Figure 5E**).

### Successful deployment and safety of injectable hydrogel electrode in a porcine model

Although the *ex vivo* model indicated strong promise of the injectable hydrogel electrode, it does not fully replicate clinical deployment in a beating, perfused heart. To this end, precursor solutions were injected into the anterior intraventricular vein using a porcine model and the *in vivo* gel formation and retention in the vein and tributaries characterized (**Figure 6A**). Explanted hydrogels displayed excellent segmental uniformity with minimal differences in equilibrium swelling ratio and gel fraction across the hydrogel length (proximal to distal). It should be noted that the calculated gel fraction does not consider any potential leaching from the gel between the time of injection until explant. The equilibrium swelling ratio provides a measure of the hydrogel network structure and indicated that the gel formation was uniform across the length of the vein (**Figure 6B**).

**Figure 6.**
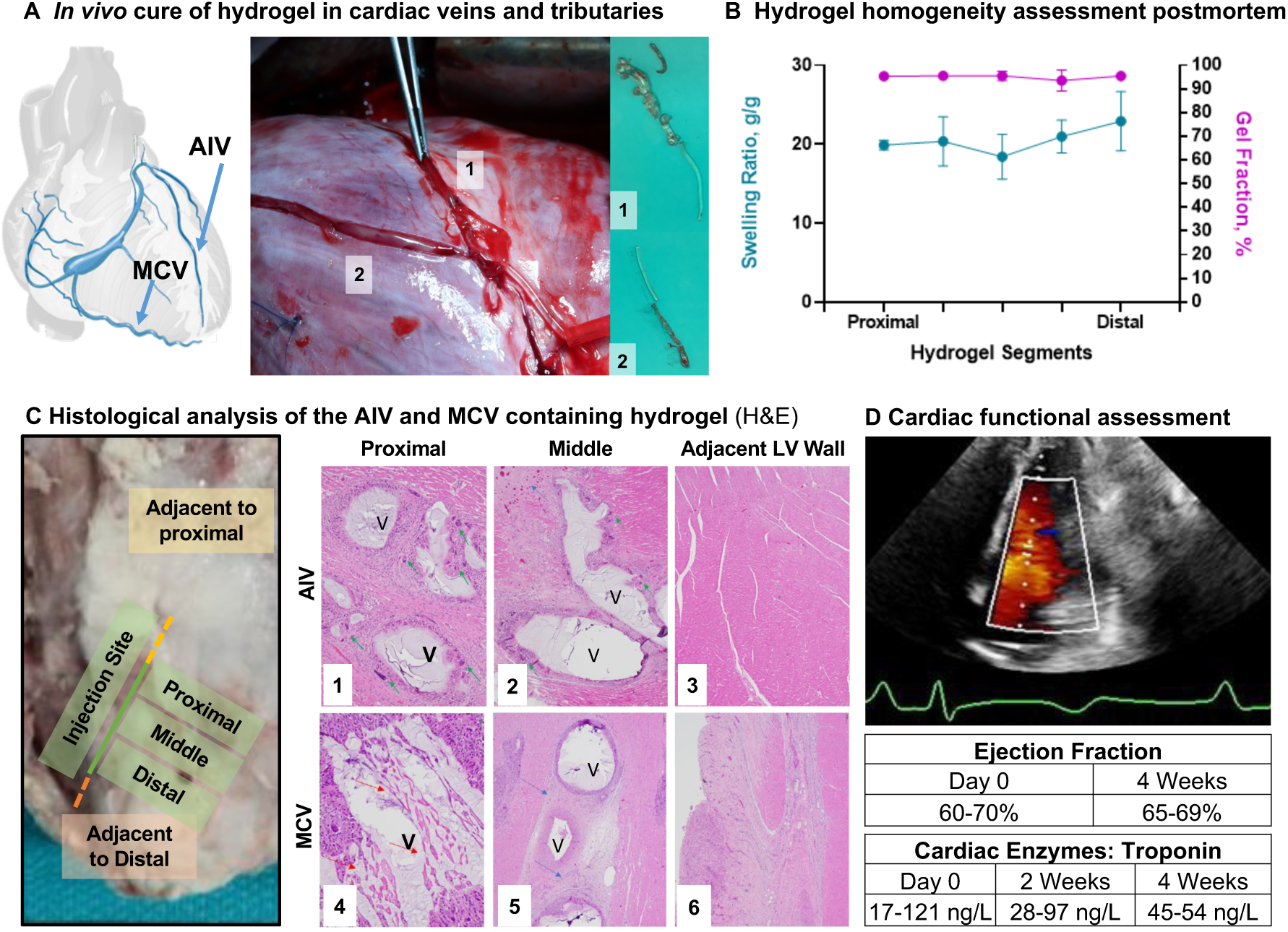
*In vivo* assessment of injectable electrode in coronary vein of a porcine model. **(A)** Postmortem confirmation of *in situ* cure of hydrogel in AIV (H1) and tributaries (H2). **(B)** Hydrogel homogeneity assessment of gel fraction and swelling ratio. Means and standard deviation are presented (n = 3). **(C)** Host response after 4 weeks (P-2165, Animal AIV 2) implantation using slices of the location proximal to the injection site, middle of the vein, and the control from an alternate section of the anterior wall. The response includes damage induced at the hydrogel injection site, with fibrosing epicarditis (red arrows) with foreign body giant cell reaction 1) fibrosing epicarditis with foreign body giant cell reaction (green arrows). Mid branch of the hydrogel injection show 2) focal replacement fibrosis, and fibrosing epicarditis. The control images at an alternate section of the anterior wall indicated 3) preserved myocardium. Histology from the MCV is also shown after 2 weeks of implantation, with 4) fibrosing epicarditis (red arrows) with foreign body giant cell reaction. Mid branch of the hydrogel injection show 5) mild focal replacement fibrosis, and fibrosing epicarditis. The control images at the distal branch indicated 6) preserved myocardium with only fibrosing epicarditis with focal extension into myocardium. Coronary vein image from IMAIOS.com. (**D**) High sensitivity troponin measurements from blood draw and measured ejection fraction from cardiac echocardiography. The echo image shown is from day of termination for Animal AIV 2. For full videos and measurements, see **Movie S3-S4**.

Once the hydrogel is injected into the vein and connected to a pacing device, it will remain in that location during the patient’s lifetime. We conducted a preliminary safety assessment to confirm that the occlusion of the vein does not result in acute adverse effects on the animal and to evaluate the host response to the gel. Unlike arterial interventions, targeting the venous vasculature offers several advantages: 1) occlusion is well tolerated clinically with no ischemic events(*19–21*); 2) lack of vascular remodelling for stable placement; and 3) low venous pressures (< 10mmHg) that are unlikely to dislodge the hydrogel as confirmed in this study.(*22*) as confirmed in this study. In clinical settings the hydrogel will be delivered to the veins via a catheter; however, for this initial feasibility study, an epicardial incision was made in order to access the vein and deliver the hydrogel. In this specific case, the hydrogel was injected into the anterior intraventricular vein (AIV) and retained for four weeks (n=3) and into the middle cardiac vein (MCV) and retained for 2 weeks (n=1). Tissue slices of location proximal, adjacent, and distal to the injection site, were excised and analyzed to assess the extent of cardiac injury and inflammation after two (**Figure S16**) and four weeks, respectively (**Figure 6C**, Figures S17-18). There was no evidence of myocardial necrosis on histopathology and no evidence of damage to the left ventricular myocardium. Mild perivascular and interstitial fibrosis was observed across all locations, with chronic epicarditis closer to the incision/injection sites (**Figure 6C**). Chronic inflammation with foreign body giant cell reaction was observed in the hydrogel-containing vessels. This reaction extended into the myocardium immediately surrounding the vessel. Replacement fibrosis was observed in some of these areas immediately surrounding the vessel. There was no diminution in myocardial contractility and no evidence of regional wall motion abnormalities at four weeks compared to baseline imaging (serial echocardiograms). The overall QRS morphology was also preserved 4 weeks after injection into the AIV (**Figures S19 – S21**) and similar two weeks after injection into the MCV (**Figures S22**). In addition to cardiac function, cardiac enzyme levels were monitored on the day of injection and the weeks after injection. Troponin levels before the procedure ranged from 17-121 ng/L with no significant change at the end of the studies (MCV, 2 weeks: 28-97 ng/L; AIV, 4 weeks: 45-54 ng/L), indicating that the hydrogel did not induce ischemia or myocardial necrosis (**Figure 6D**). No evidence of platelet accumulation and clot formation was observed in the vessels after hydrogel formation. No thrombotic activity was expected given that there was no blood flow after the epicardial vessel was occluded with the injected hydrogel. Upon necropsy after 4 weeks, the lungs were clear of clots or other signs of adverse events which supports this observation, as any venous embolization of materials would manifest in the lung.

### Hydrogel electrode mimics physiologic conduction by capturing midmyocardial tissue

The fundamental assumption in development of the injectable hydrogel electrode is that it should be able to pace the heart from the cardiac veins. At a pulse width of 5 ms, unipolar pacing thresholds were measured to be 2.2 ± 1.2 mA for the metal electrode, 2.3 ± 0.6 mA, 1.5 ± 1.28 mA for an epicardial hydrogel point source (0.8 cm disk), 1.5 ± 1.3 mA for the epicardial hydrogel line source and 2.7 ± 1.8 mA with the hydrogel in the AIV. There was no significant difference in the capture thresholds of the different electrode configurations at 5 ms and 10 ms pulse widths. Pacing thresholds measured at other pulse widths are detailed in (**Figure S23**). Two-way ANOVA revealed significant differences only for capture thresholds with pulse widths below or equal to 1 ms. As seen in **Figure 7**, epicardial pacing using a metal electrode, hydrogel point, or hydrogel line, created an inverted QRS morphology indicative of a deviation from normal conduction. Remarkably, the QRS morphology by pacing through the hydrogel in the AIV was comparable to the QRS at sinus rhythm (**Figure 7B**). No inversion of the QRS morphology and the short isoelectric window before initial activation indicate the hydrogel electrode captures the deep septal bundle branches and Purkinje fibers.(*23, 24*) In particular, the upright deflections in leads I, II, and aVL; biphasic deflections in leads III and aVF; and inverted deflections in aVR are remarkably similar between sinus rhythm and hydrogel-mediated pacing. Point pacing with metal electrodes did not yield such morphological similarities. It should be noted that the QT intervals were not identical to baseline in all cases of AIV hydrogel pacing. The primary motivation in analyzing the surface morphologies was to assess mid-myocardial capture, using native conduction system capture as a surrogate. The extensive literature in the field has focused almost exclusively of QRS morphology analysis.(*25–28*) T-wave analysis is not used clinically for this purpose. Surface 12- lead traces and corresponding analysis for each condition is included in the Supplemental Information (**Figures S24 – S38**). It can be appreciated that QRS morphologies and T-wave morphologies are far more similar with AIV hydrogel pacing as compared to all other conditions. An illustrative example is shown in **Figure 7**.

**Figure 7.**
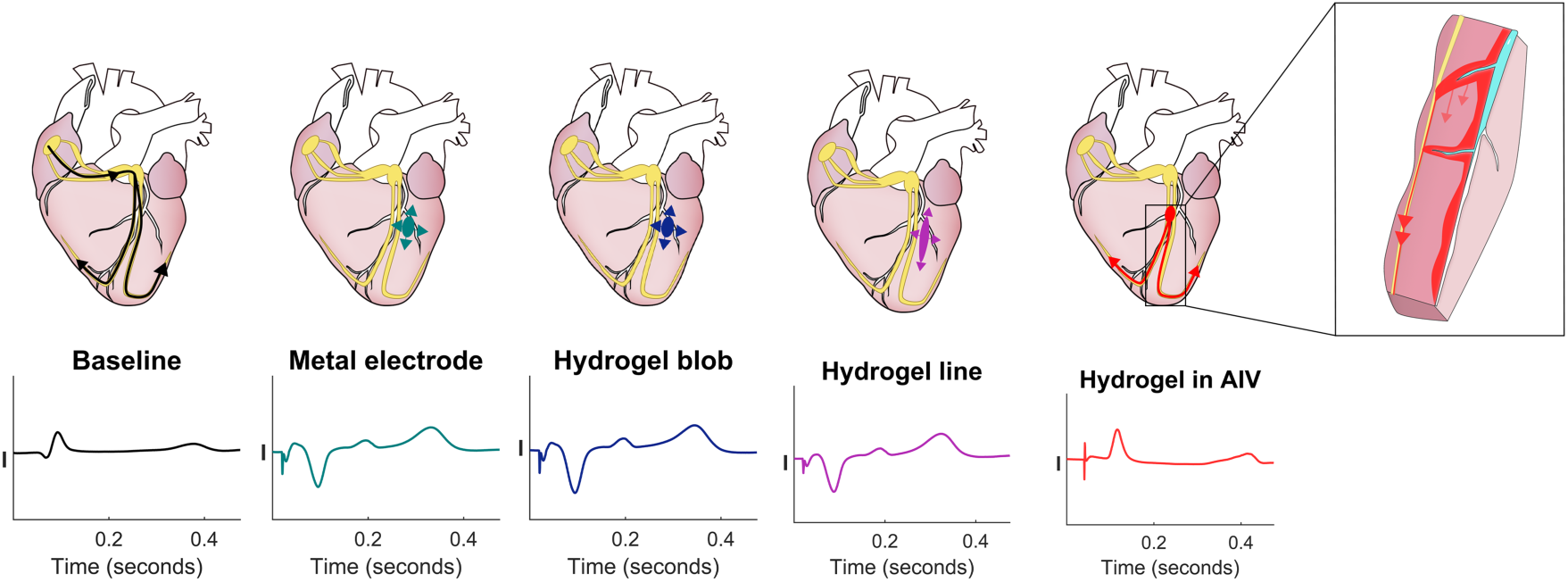
Schematic of myocardial activation with different stimulation sources. Lead I of surface ECG showing electrical activation at baseline (sinus rhythm) and when paced. Peak of the baseline QRS morphology was matched with the peak of QRS morphology obtained from pacing with hydrogel in AIV (Top, left to right) Baseline: Activation during sinus rhythm starts from the sinoatrial node and travels through the atrioventricular node into the Purkinje fibers in the ventricles. Pacing the myocardium directly causes the QRS morphology to invert indicative of deviation from normal conduction. Point pacing with metal electrode and hydrogel blob creates a dog-bone shaped activation that radiates outward from the source of the electrical stimulus. Hydrogel line pacing also created an inverted QRS morphology which indicates direct myocardial activation. Pacing from hydrogel in the AIV, clear capture is seen with a QRS morphology existing after every pacing spike. No observed inversion of the QRS morphology and a short isoelectric window before initial activation indicates possible capture of the deep septal bundle branches and Purkinje fibers (inset).

### Pacing via hydrogel electrode normalizes tissue activation across heterogeneous myocardium

Electroanatomical mapping of impulse propagation in an ablation scar model was used to study the effect of pacing from a point source and the hydrogel electrodes on conduction across heterogenous tissue in a large animal model. Briefly, ablation was performed on the epicardium of a pig heart near the AIV to disrupt the native conduction and mimic scarred myocardium conditions. Post-mortem dissection was used to demonstrate the depth of the scar formation following ablation, **Figure 8A**. Point pacing demonstrated a delayed and heterogeneous (focal) activation wavefront after ablation which was attributed to the scar formation, **Figure 8B**. The hydrogel was injected and cured in the AIV as described above and subsequent voltage mapping confirmed that the hydrogel electrode increased tissue activation area, **Figure 8C**. Moreover, the electroanatomical mapping demonstrates for the first time that the activation wavefront from the hydrogel reaches the midmyocardium and endocardium much earlier in the scar model than point pacing. The second critical observation is the broad swatch of early activation observed across the mid-myocardial and endocardial aspects with AIV hydrogel pacing. Supplementary **Figures S39- S41** confirm these findings in two independent replicates.

**Figure 8.**
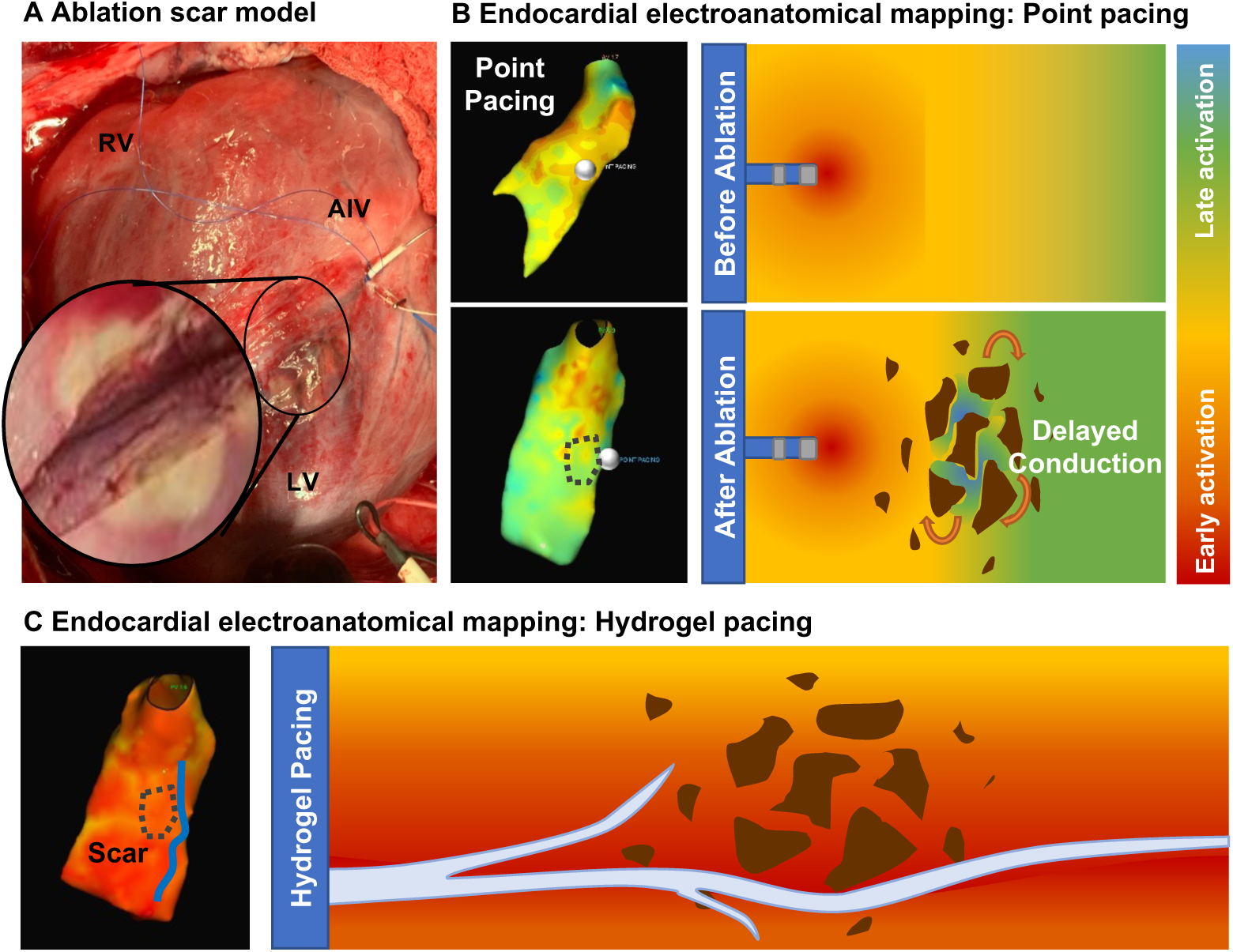
Cardiac electroanatomical mapping studies in an ablation scar model. **A)** Image of cardiac ablation near the AIV with substantial depth of the scar formation. **(B)** Endocardial electroanatomical mapping of point pacing before and after ablation with red indicating early capture and blue indicating late capture. Point pacing location is indicated by a small white circle and region of scar by dotted black line. Illustration demonstrating how point pacing allows for development of re-entrant circuits (indicated by the orange arrows) due to heterogenous capture and activation. (**C)** Endocardial electroanatomical mapping of hydrogel pacing after ablation with red indicating early capture and blue indicating late capture. Hydrogel electrode location is indicated by a blue line and region of scar by dotted black line. Illustration demonstrating how the hydrogel generates a planar wavefront through long, linear areas of capture that prevent re-entry by normalizing conduction across heterogenous tissue. Videos comparing point and hydrogel pacing can be found in the supplementary information, **Movies S5-S6**.

## DISCUSSION

Re-entry is the leading cause of lethal arrhythmias (*1*). Its most common mechanism is delayed cardiac conduction in scarred tissue, predominantly observed after coronary artery occlusion during a myocardial infarction (*29*). The ability to directly pace and capture these regions would enable targeted correction of the local activation delay; however, development and testing of such an approach has been limited by the inability to directly access and pace these regions. Given that the venous structures drain the same region where the arterial occlusion caused ischemic damage, these vessels and their tributaries provide a direct approach to the scarred tissue. However, there are no pacing leads small enough to navigate these smaller tributaries and epicardial approaches have limited efficacy due to the greater mass and thickness of ventricular tissue. In this work, we developed an injectable hydrogel electrode that can fill coronary veins to convert them into flexible electrodes that directly pace the inaccessible mid-myocardium. Hydrogels are being explored as potential flexible electrodes due to their tunable properties and ease of incorporating conductive elements (*30*). An important criterion for translation was identifying a system that was biostable, biocompatible, and can be used with current endovascular approaches. We designed a new hydrogel chemistry that could meet the multifaceted needs of this application. The PEUDAm hydrogels displayed stiffness similar to the myocardium, necessary to avoid adverse tissue responses(*31*), and increased biostability as compared to traditional PEG diacrylate. Uniaxial tensile testing indicated that the PEUDAm gels could withstand the strains imposed during a cardiac cycle; however, additional studies are needed to evaluate the long-term fatigue resistance. These hydrogel electrodes also needed adequate conductivity to ensure stimulation of the myocardium along the length of the hydrogel. Many of the conductive polymers are not suitable as an injectable electrode due to insolubility, toxicity, or processing limitations. Recent efforts to create injectable conductive hydrogels have utilized side-chain functionalization, grafting approaches, or dispersal strategies to increase solubility or maintain mild cure conditions; however, many of the resulting gels only attained conductivities at the lower end of reported myocardium values (*32*). Metallic nanoparticles are an alternative option but require high concentrations to achieve target conductivity with corollary effects on solution viscosity that can impede mixing and delivery (*33*). Our ionic hydrogel utilizes salts added to the precursor solutions to confer conductivity without detriment to other hydrogel properties, solution viscosity, or cure rate. The resulting ionic hydrogel electrode achieved conductivity more than double target myocardium values. Given that ionic conductivity is dependent on ion mobility, the conductivity of hydrogels is likely substantially different than the myocardium or blood, even at similar ions concentration. When considering living tissues such as blood, the presence of cells reduces the plasma conductivity (*34*). One advantage of the hydrogel is that the mesh size is orders of magnitude larger than ions providing a highway for ions transport in response to external voltage and small enough to exclude cell and tissue infiltration. The higher conductivity of the hydrogel with respect to tissue will allow ions to move across the gel much faster than the tissue, creating a more homogenous tissue activation along with the hydrogel. We refer to this as a planar activation. We hypothesized that the planar propagation would circumvent the heterogeneity across scar tissue, allowing us to terminate re-entrant currents around the scar at low energies. This is not possible using a single-point activation with a metal electrode. The ionic conductivity of the hydrogel was stable in vitro over numerous cycles and retained after implantation *in vivo* for up to 1 month.

For this application, an injectable system was needed that could be delivered via catheter in a minimally invasive deployment strategy and cure *in situ* without external stimuli. Multiple strategies have been used to achieve *in situ* cure, ranging from photopolymerization, ionic interactions, and temperature or pH-induced transitions (*32, 35, 36*). Our two-component system relies on delivering hydrogel precursor solutions through a dual lumen catheter to provide direct access to the diseased regions with relative precision and ease. Mixing of the two solutions upon injection into the vein activates redox-initiated crosslinking of the gel for rapid *in situ* cure. In contrast to other systems that rely on reversible bonds or physical crosslinking, our approach relies on inducing chemical crosslinking between the polymer chains via *in situ* radical polymerization to ensure long-term stability of the hydrogel. Ammonium persulfate (APS) and TEMED are one of the more common pairs used for in-situ polymerization of hydrogels (*37–42*), including cardiac applications (*37, 43*). Our lab has shown that iron gluconate (IG), a common supplement, can be used as an alternative reducing agent with a lower toxicity profile (*14*). This system is highly tunable and components can be adjusted to achieve a broad range of hydrogel properties and cure rates without changing the chemistry of the macromers. An *ex vivo* porcine model was used to identify the requisite properties for gel retention and homogeneity. A critical finding from these studies was that the initial precursor solution viscosity in combination with the hydrogel cure rate strongly influenced gel retention in the vein. A mixing head was necessary to ensure adequate mixing of the two components for hydrogel homogeneity but modeling of the mixing behavior displayed minimal effects of precursor solution viscosity. To apply these findings to endovascular delivery, we are using computationally directed design of a mixing head at the distal end of the dual lumen catheter to ensure adequate mixing for uniform gel formation and facile release of the hydrogel at the distal end upon catheter retrieval.

Regarding clinical safety, by delivering the hydrogel to the veins, we avoid arterial occlusive ischemia. Still, there may be some concern regarding cardiac engorgement consequent to the occlusion of the epicardial veins with the hydrogel electrode. However, extensive clinical experience strongly argues in favor of the safety of this practice—hundreds of thousands of occlusive leads are placed in these same vessels each time a CRT device is implanted—with no reports of complications. Longstanding literature also supports the safety of total occlusion of the coronary sinus (into which all venous tributaries drain) - a vessel much larger than the branch vessels we will be targeting (*19–21*). There is a theoretical risk of edema due to reduced venous return after occlusion; however, we expect minimal effect due to established venous remodeling under similar conditions. Pilot large animal data shows that occluding the epicardial venous tributaries was associated with minimal, if any, myocardial necrosis or loss of cardiac function. Histopathologic studies demonstrated myocardial inflammation that was increased at the regions directly instrumented in this open chest model. Clinically, such a traumatic delivery system is unlikely to be utilized. Our delivery system is currently being modified for transcatheter delivery using a dual lumen catheter. Gaining access to these venous branches using a closed chest transcatheter approach is a standard skill set utilized by clinical electrophysiologists when placing a cardiac resynchronization device (a pacemaker with a lead in the ventricular branches of the epicardial venous system). Such an approach will reduce the observed inflammatory response at the injection site. The extent of inflammation in the myocardium and the replacement fibrosis observed immediately surrounding the vessels containing the hydrogel in the chronic porcine studies warrants further investigation to determine its origin and temporal stability. Given that the cytocompatibility and subcutaneous implant study support the biocompatibility of the hydrogel, additional studies will explore vessel dilation and mechanical disruption of the vessel with hydrogel fill ratios and the corollary effect on remodeling. Most notably for early demonstration of the safety of this approach, cardiac function was preserved and cardiac enzymes at the end of the study were within normal ranges, indicating a lack of myocardial necrosis or serious disease state. Furthermore, our preliminary data indicated no gross emboli in the lungs which indicates that full occlusion with the hydrogel does not carry the risk of platelet accumulation or embolism.

With this new injectable hydrogel electrode, we are able to explore the possibility of *prevention* of re-entry by directly capturing zones of delayed conduction, thus eliminating one of the necessary conditions for re-entry to arise. Unlike past efforts that used conductive hydrogels or other materials as scaffolds to regenerate myocardial tissue and restore conductivity, (*44–46*) these conductive hydrogels will interface with current cardiac resynchronization devices and act as biostable, flexible electrodes to provide an immediate intervention to prevent VA. By injecting into the corresponding venous branches with the conductive hydrogels, electrophysiologists can target specific locations corresponding to each patient’s scar location(s), enabling patient-specific cardiac resynchronization paradigms involving increased capture areas compared to currently available lead-based systems by virtue of the mid-myocardial nature of the hydrogel electrode. The advantage of direct midmyocardial tissue capture is not limited to prevention of re-entry. Recent progress in atrial defibrillation technology has demonstrated that using multiple electrodes to deliver multi-stage electrotherapy can reduce the energy required defibrillation (*47*). Delivering multiple low-energy pulses over a larger area has shown benefits for terminating fibrillation with energies below pain thresholds (*48–50*). Moreno et al. performed an *ex vivo* study with epicardial line electrodes that generated a planar activation wavefront that depolarized a large ventricular area at energy levels below the human pain threshold (*51*). These reports indicate that planar wavefront propagation from the hydrogel electrode as it spans the mid-myocardium also provides the opportunity for painless ventricular defibrillation. Other multi-site pacing approaches to prevent and treat VA rely on epicardial placement of flexible electrodes. The lack of a minimally invasive deployment strategy of these hydrogels as flexible electrodes represents a large translational challenge for this technology. Furthermore, epicardial stimuli are less effective for defibrillation due to the greater mass and thickness of ventricular tissue. In contrast, this endovascular approach provides unprecedented simultaneous multi-site pacing in the mid-myocardium while also integrating with current clinical practices. In our preliminary *in vivo* studies, once the hydrogel was injected into the AIV, we observed uniform capture along the vessel both pre and post scar formation as well as increased capture compared to point pacing. This data supports the idea that the planar wavefront will persist without causing re-entrant activity.

Cardiac pacing studies using these injectable hydrogel electrodes in a large animal model provided proof-of-concept for this new approach. Although we did not assess the effect of such pacing in diseased myocardium, our work demonstrates for the first time the ability to directly pace these erstwhile unapproachable regions of the heart. The evidence to support this is the nearly identical nature of surface electrocardiographic morphologies of the paced and native heartbeats. A similar effect normalization of QRS vectors has been observed in clinical reports of deep septal pacing (*52–54*). Furthermore, there is a marked latency between the pacing stimulus and initiation of the surface ECG. This is because the pacing output at least in part captures and conducts via the native conduction system before exiting into myocardial tissue. Thus, there is a delay between the pacing stimulus and onset of the ECG signal. These features were reproducibly observed and are strongly suggestive of direct mid-myocardial tissue capture, something that has been technologically vexing and, to the best of our knowledge, unreported. This first evidence that these electrodes can mimic physiologic conduction could alter the landscape of cardiac rhythm management. Collectively, these injectable hydrogel electrodes show strong promise as a new cardiac resynchronization therapy to provide direct stimulation of wide areas of the heart.

## MATERIALS AND METHODS

### Materials

All materials were purchased from Sigma Aldrich unless otherwise indicated.

### Synthesis of polyether urethane diacrylamide (PEUDAm)

A new PEUDAm macromer was synthesized in a three-step process with a PEG-diamine intermediate. *Synthesis of PEG-CDI:* Carbodiimidizole (CDI, Alfa Aeser, 15 equiv.) was weighed in a dry, 500 mL round bottom flask and dissolved in anhydrous DCM (20 mL). PEG (Mn = 17600, 1 equiv.) was dissolved in anhydrous DCM (10 wt%) and placed under nitrogen atmosphere. The PEG solution was added dropwise to CDI over 2 hours at 30-35°C, then left to react at room temperature overnight, stirring vigorously. The reaction was quenched with the addition of deionized water and the solution was then washed with deionized water three times in a separatory flask, dried with sodium sulfate, and collected via vacuum filtration. Solvent was removed under vacuum with additional drying of the product at 65°C under vacuum for 1 hour to remove residual solvent and water prior to use. *Synthesis of PEG-Diamine:* Ethylenediamine (15 equiv.) was added to a dry 500 mL round bottom flask and a PEG-CDI solution in anhydrous DCM (12 wt%, 1 equiv.) was added dropwise at RT over 3 hours (1 drop per 1-3 seconds) under nitrogen. The reaction was allowed to proceed overnight under vigorous mixing. The solution was then washed with deionized water three times in a separatory flask, dried with sodium sulfate, and collected via vacuum filtration. The product was precipitated in 10x volume of ice-cold ether and collected via suction filtration. The product was then dried under vacuum at 80°C for 3 hrs to remove residual solvent and water. *Synthesis of PEUDAm*: PEG-diamine (1 equiv.) was weighed in a dried 500 mL 3-neck round bottom flask and dissolved in anhydrous DCM at 15 wt% under nitrogen atmosphere. Triethylamine (2 equiv.) was added dropwise. Acryloyl chloride was diluted in DCM (∼4 mL) and added dropwise (1 drop per 4-5 s). The reaction was allowed to proceed for 48 hours at room temperature, then quenched with potassium carbonate (2 M, 8 equiv.). The solution was washed twice with water, dried with sodium sulfate, and collected via suction filtration. The product was precipitated in 10x volume ice-cold diethyl ether and collected via suction filtration. The final product was dried at atmospheric pressure overnight, then briefly under vacuum.

### Macromer characterization

Functionalization of PEG at each step was determined by analyzing proton nuclear magnetic resonance (H^1^ NMR) spectra collected on a 300 MHz Bruker NMR. H^1^ NMR (CDCl3): *PEG-CDI*: δ = 8.12 (s, 2H, -N-C*H*-N-), 7.40 (s, 2H, -N-C*H*=CH), 7.02 (s, 2H, - CH=C*H*-N), 4.51 (m, 4H, -O-C*H2*-CH2-), 3.70 (m, 1720H, -O-C*H2*-CH2-). *PEG-EDA*: δ = 4.18 (t, 4H, -O-C*H2*-CH2-), 3.60 (m, 1720H, -O-C*H2*-CH2-), 3.19 (m, 4H, -CH2-C*H2*-NH), 2.78 (m, 4H, - NH2-C*H2*-CH2-), 3.70 (m, 1720H, -O-C*H2*-CH2-). *PEUDAm*: δ = 6.94 (broad s, 2H, -C-N*H*-CH2-), 6.28 (dd, 2H, *H*2C=CH-C-), 6.17 (m, 2H, H2C=C*H*-C-), 5.85 (broad s, 2H, -C-N*H*-CH2-), 5.61(dd, 2H, *H*2C=CH-C-), 4.20(m, 4H, -H2C-C*H2*-O-), 3.65 (m, 1720H, -O-C*H2*-CH2-), 3.34 (m, 4H, -CH2-C*H2*-NH-). Macromer molecular weight was determined by gel permeation chromatography (GPC) using an Agilent Technologies 1260 Infinity II GPC system. Macromer specimens were dissolved in 0.1 M lithium bromide in dimethylformamide at a concentration of 5 mg/mL at room temperature and syringe filtered. Injections (100 μL) were passed through 2 Phenogel columns with molecular weight ranges of 1–75 kDa at a flow rate of 1.0 mL/min and temperature of 25 °C. The separated components were passed through an Agilent Multi Detector system (RI, RALS, LALS, UV) at 30 °C, and the molecular weight distribution was determined using the Agilent GPC/SEC software. Molecular weight was reported relative to polyethylene glycol standards. Viscosity was measured in triplicate using a Brookfield Viscometer equipped with a CP-40Z flat plate.

### Synthesis of *N*-acryloyl glycinamide

*N*-acryloyl glycinamide was synthesized according to established protocols with minor variations (*10*). Glycinamide hydrochloride (Alfa Aeser, 6.3 g, 1 equiv.) was added to a 250 mL 3-neck RBF, dissolved in 18 mL diethyl ether and 40 mL of 2 M potassium carbonate (1.18 equiv.), and cooled over ice. Acryloyl chloride (0.98 equiv) was diluted in 24 mL cold diethyl ether and added dropwise to the reaction mixture over 30 minutes with vigorous stirring. The reaction was allowed to proceed for 4 hours at room temperature while protected from light. The solution was reduced to pH = 2 with 6 M HCl and then washed with diethyl ether (50 mL) three times. The aqueous solution was then placed under vacuum to remove residual ether, adjusted to neutral pH with 2 M NaOH, then frozen and lyophilized. Warm acetone (200 mL, 40°C) was added to the powdered product and allowed to stir for 8 min to extract the product. This process was repeated four times and approximately half of the solvent was removed via rotary evaporation. The dissolved product was crystallized on an ice salt bath. The crystals were collected and washed then dissolved in minimal warm methanol. Undissolved particles were removed, and the product was recrystallized on ice salt. The product was then dissolved in deionized water, frozen, and lyophilized before analysis with NMR. H^1^ NMR (300 MHz, D2O): δ = 6.31 (dd, 1H, -HC=C*H2*), 6.23(dd, 1H, -HC=C*H2*), 5.80 (dd, 1H, -*H*C=CH2), 3.97 (s, 2H, HN- C*H2*-CONH2)

### Hydrogel fabrication and gelation kinetics

Hydrogels were fabricated with the PEUDAm macromer (20 wt%), NAGA monomer (1 wt%) and a combination of redox initiators ammonium persulfate (APS, 0.75 mM) and iron gluconate (IG, 1.5 mM) in saline solution (1.35% NaCl). Hydrogel solutions were loaded into disposable double-barrel syringes with half the solution containing APS and the other half containing IG. The cure profiles of the injectable hydrogel was characterized by determining gelation onset and complete network formation using a Discovery Hybrid HR-2 rheometer. Hydrogel precursor solutions were prepared at an initiator concentration of 0.5, 0.75, or 1.0 mM with an APS:IG molar ratio of 1.0:2.0, loaded into double-barrel syringes, and injected through a mixing head onto a parallel-plate (20 mm) configuration heated to 37 °C. Storage and loss moduli were measured every 6 seconds with a 1.3 mm gap and 0.5% strain. To determine the effect of blood on cure rate, 50 μL of porcine blood was added at the center of the plate, and the hydrogel was injected directly into the blood. The onset of gelation was measured as the crossover point of loss and storage modulus. Complete network formation was measured as the third point after which there was a less than 1% change in storage modulus. Values were reported as the average of three specimens. To determine tensile properties, dog bone specimens (n = 4) with a gauge width of 5 mm width were cut from swollen hydrogel slabs (thickness∼1.5mm) using a modified 3D printed punch developed by Nelson *et al.* (*55*). Specimens were strained to failure using a dynamic mechanical analyzer (RSAIII, TA Instruments) at a rate of 0.1 mm/s. All mechanical tests were performed at room temperature.

### Characterization of network formation

Macromer solutions were prepared as described above, injected through a mixing head into glass spacers (0.75mm), and allowed to cure for at least 30 minutes at 37°C prior to characterization. Network formation of the injectable hydrogel was determined by measuring gel fraction and swelling ratio. To measure the gel fraction, four specimens (10mm diameter, 0.75 mm height) were dried under vacuum for 24 hours and weighed. The salt content of the precursors was considered in the calculation of initial polymer mass (W0). Specimens were swelled for 24 hours in 2 mL of DI water and weighed. The specimens were dried again for 24 hours to assess the remaining dry polyme1r mass (Wf). The ratio of the final to initial mass (Wf /W0) was reported as the gel fraction. To measure equilibrium swelling ratio, four specimens (10mm diameter, 0.75 mm height) were swelled in DI water for 24 hours and weighed (Ws). The specimens were then dried under vacuum for 24 hours and weighed (WD). The equilibrium mass swelling ratio was calculated as Q=Ws/WD. To determine the homogeneity of gels after injection, hydrogel solutions were prepared and injected into a 3/16” ID PVC Tygon tubing (McMaster-Carr) using the double-barrel syringe with mixing head. The gels were allowed to cure for at least 10 minutes at 37°C. The gel was cut into five segments for characterization of gel fraction and equilibrium swelling as described above.

### Accelerated degradation testing

To confirm the hydrolytic biostability of the new hydrogel composition, accelerated degradation testing of the PEUDAm hydrogels was performed in comparison to a polyethylene glycol diacrylate (PEGDA, 20 kDa, 20wt%) control. Hydrogel specimens (10 mm, n = 4) were placed in PBS (3 mL) in glass vials and incubated at elevated temperatures (70°C) or physiological temperature (37°C) with weekly solution changes. Increases in equilibrium swelling and reductions in compressive storage modulus were used as a measure of hydrolytic degradation. The hydrogels were weighed weekly for 8 weeks and the change in equilibrium swelling was calculated as: equilibrium swelling mass at each time point divided by initial equilibrium swelling mass. For modulus measurements, specimens were tested monthly for 3 months using a dynamic mechanical analyzer (RSAIII, TA Instruments) equipped with a parallel-plate compression clamp. Testing was performed under unconstrained compression at room temperature. Compressive modulus was determined from a frequency sweep at 1 % strain.

The compressive storage modulus was taken at 0.25 Hz in the linear region. Four specimens were tested per gel.

### Cytocompatibility

An indirect assay was used to determine the cytocompatibility of hydrogel leachables immediately following cure. Human umbilical vein endothelial cells (HUVECs) were expanded in Endothelial Cell Growth Medium-2 (Lonza), harvested for use between passage P4- P6, and seeded at 30,000 cells per well in 24 well plates. Human induced pluripotent stem cells (hiPSCs, WTC 11), a gift from Dr. Bruce Conklin, were cultured via feeder-free conditions on vitronectin coated well plates in complete Essential 8 (E8) medium (ThermoFisher). Cardiomyocyte (CM) differentiation was conducted via a modified protocol based on the established Wnt GiWi method (*56, 57*). Briefly, cells were detached with Accutase ® and reseeded at 25,000 cells/cm^2^ on Matrigel® (Corning) coated plates. Following incubation in E8 media supplemented with 10µM ROCK inhibitor for 24 hours, cells were expanded for additional 48 hours. CM differentiation was initiated using 12µM of CHIR99021 (LC Laboratories), in RPMI 1640 supplemented with B27™ without insulin (RB-, Life Technologies). After 24 hours incubation at 37°C, CHIR99021 was removed and replaced with RB- medium for 48 hours. Fresh RB- medium, supplemented with 5µM IWP-2 (Tocris), was then exchanged, and cells were incubated for 48 hours. Next, medium was exchanged for fresh RB- for 48 hours. Finally, medium was replaced with RPMI 1640 supplemented with B27™ and insulin (RB+, Life Technologies). On Day 11, spontaneously beating CMs were reseeded onto Matrigel coated 24 well plates at 50,000 cells per well. After 24-hours, HUVECs and CMs were indirectly exposed to freshly-made hydrogels using a Transwell® insert. The hydrogels were prepared by adding all reagents into a 96 well plate (100μL hydrogels), mixing via pipette, and curing for 30 minutes at 37°C. Upon curing, the hydrogels were transferred to the Transwell® insert and the viability of the cells was determined after 24 hours of contact with leachables from the hydrogel. Cells were stained green-fluorescent calcein-AM and red-fluorescent ethidium homodimer-1 (Live/Dead Viability/Cytotoxicity Kit, Thermofisher). Imaging was conducted with a fluorescence microscope (Nikon Eclipse TE2000-S) and cells were manually counted (n = 3 images per well, 3 replicates).

### *Ex-vivo* gel formation and retention

Hydrogel retention and network formation were tested in an *ex vivo* pig heart model to identify requisite properties of the injectable hydrogel prior to *in vivo* studies. The effect of viscosity was investigated by varying the concentration of the PEUDAm macromer solution from 10 wt% (42 mPa-s) to 20 wt% (83 mPa-s). The effect of cure rate was investigated by varying the initiator concentration from 0.5:1.0 mmol APS:IG to 0.75:1.5 mmol APS:IG. Pig hearts were purchased from Animal Technologies. Increasing the APS concentration up to 1 mM resulted in premature gelation in the catheter and was not tested in the *ex vivo* model. Porcine hearts were submerged in a water bath at 37°C until thermal equilibrium was achieved. An incision was made to access the cardiac vein and porcine blood was used to fill the veins prior to injecting the hydrogel solution. Hydrogel precursor solutions were prepared as described above and loaded into the double-barrel syringe. A catheter was connected to the mixing head to deliver the hydrogel into the veins and the catheter and mixing head were purged with saline to avoid air bubbles. After injection, the hydrogel was allowed to cure for at least 30 minutes and then removed using a scalpel to access the vein. The gel was divided into five segments for characterization of gel fraction and equilibrium swelling as described above. To investigate the key determinants of gel retention, the effect of viscosity on mixing was modeled and the increase in complex viscosity as a function of macromer viscosity and cure rate was measured. The complex viscosity was determined from the gelation kinetics experiments described above. The precursors were dispensed through a mixing head onto a parallel-plate (20 mm) configuration heated to 37 °C. Complex viscosity was measured every 6 seconds.

### Flow simulations

Simulations were performed using SolidWorks Flow Simulation on a branched vessel constructed in SolidWorks 2021 (Dassault Systèmes, Vélizy-Villacoublay, France) with dimensions in the range of typical cardiac vessel sizes (main vessel ID=3mm). A standard spiral mixing head was modeled and inserted into the main branch of the vessel model. Ambient pressure conditions were present at each outlet, and a zero flow condition was applied to the inlet. The vessel model was filled with liquid matching blood viscosity to begin the simulation study. A transient analysis was performed in which the precursor solutions were injected through the mixing head at a rate of 3.5 mL/s for 0.5s, after which the study continued until t=15 seconds. The viscosity of the precursors was adjusted to assess the impact on mixing and initial flow into the vessels.

### Electrical characterization

The resistance of the hydrogel specimens (n = 4) was determined using a four-point probe and electrochemical impedance spectroscopy. The gels were placed over a glass slide and the 4-point probe (Signatone 4Pro) was placed in contact with the center of the specimen. The Voltage-Current characteristics of the hydrogels were recorded by varying the current from 0.001mA to 0.2mA. The resistance (R) of the sample was obtained from the slope of the V-I plot. For impedance analysis, the hydrogel constructs were placed between two electrodes with an alternate current sweep from 1 to 10^6^ Hz. The resistance (R) of the sample was obtained when the impedance plateau in the bode plot. Conductivity was calculated as the inverse of the resistance multiplied by the gel thickness (L) and divided by the area (A). Additionally, the conductivity of ionic species present was determined using an Oakton CON 450 portable conductivity meter. Specimens were submerged in 5mL of DI water and allowed to reach equilibrium. The recorded conductivity value was compared to a standard curve to determine the hydrogel’s relative ion concentration and ionic conductivity. Monophasic pulse signals (5 msec) were generated by a function generator (Rigol DG4000) at a frequency of 2 Hz. An oscilloscope (Tektronix MDO3000) was used to measure the voltage drop to a resistor connected in series with the hydrogel in order to calculate the current as a function of hydrogel length. The circuit was connected to a 5 volts power supply. The schematics of the circuit setup are shown in **Figure S15**.

### In-vivo assessment of gel conductivity retention

All procedures were approved by the IACUC at the Texas Heart Institute. Hydrogels prepared in saline (0.9% NaCl) were implanted subcutaneously in a rat model to evaluate the effect of implantation on the ionic conductivity. Hydrogels were prepared and maintained under sterile conditions until implantation. All precursor and redox solutions were filtered using a 0.22-micron size filter. Specimens (5 mm by 10 mm) were cut and enclosed in sterile cylindrical stainless steel mesh cages (16 mm length, 8 mm diameter) prepared at the UT Austin laboratory as describe above. The wire cages and glass plates were cleaned with 1% alkaline detergent and sterilized with UV light. Gels were submerged into phosphate buffered saline (PBS 1X). Solution was changed three times and gels were allowed to reach equilibrium swelling for at least 24 hours. Gels were kept submerged until implantation. Fischer rats were acquired and quarantined for 7 days prior to the procedure. The rats were anesthetized using 1.5% Isoflurane through a face mask that was secured to the operating table, hair was removed using clippers and depilatory cream. Incisions were made on the side of the rats, and four subcutaneous pockets were prepared above the fascia of the external oblique muscle to implant one cage with a hydrogel in each pocket. The incisions were then closed with 4-0 Vicryl sutures. After 7, 14 or 28 days, rats were euthanized under deep anesthesia via intraperitoneal injection of Pentobarbital solution (Euthasol). The stainless steel cages were explanted, and hydrogel specimens were carefully removed from the cages. The ionic conductivity was assessed using a hand-held conductivity probe. The sol fraction and swelling ratio were measured as described above.

### In vivo biocompatibility assessment in a subcutaneous rat model

Animal procedures were approved by the IACUC at the Texas Heart Institute. Fisher rats were prepared for survival study. All rats will be anesthetized before and during the procedure. The animals were individually placed in an Isoflurane (5%) Chamber for 3-5 minutes to induce anesthesia. Next, the animals were moved to mask anesthesia (Isoflurane 3%) for 3-5 minutes. During this time, pedal responses were continuously monitored for adequate anesthesia. Once the rat was fully anesthetized, the rat was positioned (Ventral recumbency) for surgery. The rats were then shaved. We also utilized Nair to remove additional hair. Nair was applied to the shaved area for 30 seconds. Then it was removed with a clean towel and sterile water. Next, we scrubbed the exposed skin with surgical-grade Betadine. Two incisions were made in the skin about 2 cm above the tail and along the midline. Blunt dissection will be used to prepare 2 implant pockets (1 per side). Specimens were aseptically introduced through the incision and positioned within the pocket and away from the incision site, and a preformed hydrogel disc (12 mm diameter, 1.6 mm thickness after equilibrium swelling) and a silicone disc (10 mm diameter, 0.5 mm thickness) were placed. The incisions were closed with 4-0 Vicryl simple continuous sutures. The rodents were also injected with saline and hydrogel in two separate locations in the upper back. The rats were then recovered and kept in a vivarium for 1, 2, and 4 weeks. At the end of each respective time period, the rats for that time point were euthanized, and the entire dorsal tissue was removed for histology.

After sacrifice, the four separate areas of interest containing the designated implants were carefully dissected and fixed in 10% neutral buffered formalin (NBF) for at least 48 hours. After fixation, silicone and hydrogel disc samples were sliced approximately across the middle of the disc (point of largest diameter). Subcutaneous hydrogel clumps were sliced perpendicular to their longest axis. The samples were then processed for paraffin embedding, sectioned at 5 µm thickness, and stained with hematoxylin and eosin (H&E) and Mason’s trichrome. Histologic sections were evaluated by light microscopy and assessed for the presence of inflammatory cells (polymorphonuclear leukocytes, lymphocytes, macrophages, and foreign body giant cells), fibroblasts, and vascularization, each of which were scored from - to +++, absent to markedly present, and -/+, occasional. Peri-implant fibrous capsule thickness measurements were taken with Image-Pro Premier software, version 9.3.3 (Media Cybernetics, Inc.).

### *In vivo* deployment and pacing in a porcine model

Animal procedures were approved by the IACUC at the Texas Heart Institute. Yorkshire pigs (n=3) between 40kg and 50kgs were used for these studies. The animal was sedated and prepared for a non-survival study. The animal’s vitals were constantly monitored using a 12-lead ECG for the entirety of the procedure. A median sternotomy was performed on the pig to expose the anterior surface of the heart. The anterior intraventricular vein (AIV) branch that would eventually be utilized for injection was identified prior to pacing, and four decapolar catheters were sutured around the target site in a square for sensing the local epicardial electrograms. The pacing target was selected close to the expected injection site on the AIV. Unipolar cathodal pacing using a bare metal electrode, hydrogel point, and hydrogel line were performed with the indifferent electrode placed on the animal’s chest.

Three different modes of pacing were attempted – point pacing, line pacing, and pacing with the hydrogel in the AIV. Hydrogel precursors were prepared as described above and loaded into a double-barrel syringe with a mixing head. To prepare the hydrogel for point pacing, approximately 500μL of hydrogel was dispensed into a 2 mL tube containing an exposed stainless-steel wire such that that the gel cured (∼ 8 mm diameter) around the end of the wire. To prepare the hydrogel line source, the hydrogel precursors were injected into a 14G angiocatheter and allowed to cure for 2 minutes to generate a cylindrical hydrogel line source for epicardial pacing (1.6 mm in diameter, 8 cm in length). The minimum current required to achieve reliable pacing capture (called capture threshold) at a constant pulse width (0.5, 1, 5, and 10ms) was recorded for each pacing mode, yielding different strength-duration curves. After completion of epicardial pacing, the hydrogel was injected into the AIV, using a 14G angiocatheter inserted directly into the AIV with a suture at the proximal end to minimize bleeding and backflow of the hydrogel precursor solutions upon injection. A double barrel syringe with a mixing head was loaded with 2 mL of hydrogel precursor solutions and attached to the angiocatheter via the luer lock system described above. After allowing the gel to cure for ∼2 minutes, the catheter was and a pacing lead was attached to the hydrogel at the proximal end using an alligator clip. Capture threshold and strength-duration curves were again assessed with the intravascular hydrogel. At the end of the study, all animals were humanely euthanized per standard IACUC approved practice. After euthanasia, the AIV was carefully opened using a scalpel and the cured hydrogel was carefully excised from the AIV for inspection. The hydrogel was then sectioned and characterized for sol fraction and swelling ratio as described above.

### Safety assessment in a porcine model

Animal procedures were approved by the IACUC at the Texas Heart Institute. Yorkshire pigs between 40kg and 50kgs were utilized for the chronic assessment of the safety of the injected hydrogel vein occlusion of the AIV (n = 3) and the (MCV, n = 1). The pigs were sedated and prepared for a chronic study, with vitals and ECG recorded and monitored throughout the study. For the AIV, a median sternotomy was performed on the pig to expose the anterior surface of the heart. The sterile hydrogel precursor solutions, prepared as described above, were injected via the double-barrel syringe into the vein. The gel was allowed to cure for 5 minutes before catheter removal. For the MCV, a thoracotomy was performed to expose the left side of the heart. The largest vein in that window, the MCV, was identified by the surgeon and accessed using a 16G angiocatheter. The sterile hydrogel precursor solutions, prepared as described above, were injected via the double-barrel syringe into the vein. The gel was allowed to cure for 5 minutes before catheter removal. A single prolene suture was placed at the injection site to minimize any potential bleeding and as a marker for histological analysis after 2 (MCV, n=1) and 4 (AIV, n=3) weeks. Echo cardiograms and blood enzymes were drawn at the beginning and end of the study to assess myocardial changes. The animal was allowed to recover and was observed for 2 and 4 weeks prior to euthanization. The animals were prepared for termination, and ECGs, echocardiogram, and enzyme blood draws were repeated before termination. After euthanized was complete, the heart was removed and placed in fixative for histological analysis. A small section of hydrogel was also removed for sol fraction and swelling ratio analysis prior to fixation. The heart was sectioned along the length of the hydrogel injection to include the proximal vein for site of injection, the mid-vein, and the distal vein, all approximately 2 cm apart. Paraffin-embedded sections were stained with either hematoxylin and eosin, Masson’s Trichrome, or Masson’s Pentachrome to assess cardiac injury and inflammatory responses of the tissue to the chronic hydrogel implant.

### Voltage mapping of pacing in a porcine ablation model

Yorkshire pigs were prepared (n=3) as per general protocol for an acute procedure approved by the Texas Heart Institute IACUC committee. A midline sternotomy was performed to expose the heart. A baseline sinus rhythm activation map was measured on the epicardium, endocardial left side, and endocardial right side utilizing the EnSite Precision Cardiac Mapping System (Abbot St. Jude, IL). After sinus mapping, the heart was paced at the expected site of hydrogel injection using a temporary pacing lead and mapped again at the same locations described above. Pacing was performed using the BARD Electrophysiology system (Boston Scientific, MA). The hydrogel electrode was then injected and paced as described previously. Activation was mapped during hydrogel pacing in the same locations as above. After mapping these three conditions, ablation was performed distal to the injection site near the AIV to mimic cardiac scarring. Approximately 5 ablations were performed in a lateral line at 50 W for 30 s with saline irrigation using the 8Fr THERMOCOOL Catheter (Biosense Webster, CA) with CoolFlow irrigation pump (Biosense Webster, CA) and Stockert 70 RF Generator System (Biosense Webster, CA). After ablation, the three pacing conditions (sinus rhythm; temporary pacing lead; hydrogel) were repeated along with their associated mapping procedures (n=3). The animals were terminated upon completion of the procedure.

### Statistical Analysis

Quantitative characterization of hydrogel properties is expressed as mean ± standard deviation. Statistical comparisons were made using the Student’s t test for paired data and analysis of variance (ANOVA) for multiple comparisons with Tukey post hoc analysis for parametric data. Computations were performed using GraphPad Prism version 9.0.2 at the significance levels of p < 0.05.

## Supporting information

Movie S1. Mixing of precursors solutions using a dual lumen catheter for dispensing.

Movie S2. Mixing of precursors solutions when dispensed using a double barrel syringe with a mixing head.

Movie S4. Echocardiogram prior injection on the AIV in a pig model.

Movie S4. Echocardiogram 4 weeks after injection on the AIV in a pig model.

Movie S5. Myocardium activation comparison after pacing with the hydrogel in the vein for Animal AIV 1

Movie S6. Myocardium activation comparison after pacing with a single point (metal electrode)

Movie S3. Echocardiogram 2 weeks after injection on the MCV in a pig model.

Movie S6. Myocardium activation comparison after pacing with the hydrogel in the vein for Animal AIV 2

Movie S5. Myocardium activation comparison after pacing with a single point (metal electrode)

## Supplementary Materials

Figure S1. Synthetic route for PEUDAm.

Figure S2. H1 NMR Spectra of PEG-CDI.

Figure S3. H1 NMR Spectra of PEG-EDA.

Figure S4. H1 NMR Spectra of PEUDAm.

Figure S5. Gel Permeation Chromatogram of PEUDAm macromer.

Figure S6. H1 NMR spectra of NAGA.

Figure S7. PEUDAm stability under hydrolytic conditions. PEGDA 20kDa, 20% + 1% NAGA was used as a positive control to assess improved biostability of PEUDAm 20kDa, 20% + 1% NAGA gels. Hydrogel swelling of (A) PEGDA and (B) PEUDAm in PBS was recorded as a function of time at 37°C and under accelerated conditions (70°C). Means and standard deviations are presented (n = 4). In most cases, the standard deviation is smaller than the bullets’ size and was not displayed by the software. Storage modulus of PEUDAm hydrogels (C) under 1% strain at 0.5 Hz (compression) (n = 4).

Figure S8. Rheological data: Time sweep of PEUDAm hydrogels. Time sweep of PEUDAm 20kDa, 15% cured at a concentration at 0.5, 0.75, and 1.0mM APS. The concentration of IG was twice the APS concentration: 1.0, 1.5, and 2.0, respectively.

Figure S9. Effect on time to complete network formation. A) Effect of PEUDAm molecular weight (at 15 w/w%). B) Effect of adding 1% NAGA (to a PEUDAm 20k, 15% solution). C) Effect of contact with blood.

Figure S10. Cytocompatibility by indirect contact with the hydrogels. Percentage of HUVECs and hiPSC-cardiomyocytes viability 1 day after continuous exposure to hydrogel in comparison with TCPS. All data represents average ± standard deviation for n = 3 wells (x 3 location per well). There was no statistical differences when comparing to the TCPS control

Figure S11. Biocompatibility of PEUDAm hydrogel. Representative higher magnification images showing the host response and evolution of a thin fibrous cap around the implanted hydrogel discs. (A, B) At 1 week post-implant, the interface between the hydrogel disc and surrounding tissue is composed mostly of macrophages, fibroblasts and capillaries (neovascularization, arrows) within a background of incipient extracellular matrix deposition. (C, D) By 4 weeks post-implant, a hypocellular fibrous capsule with increased collagen content has developed. Note the interface has remained of similar thickness (double headed arrows). Stains: H&E (left column) Masson’s trichrome (right column). Scale bar = 20 μm.

Figure S12. Conductivity of hydrogel determined by 4-point probe and conductivity probe. A) Comparison of hydrogels conductivity using the four-point probe and the hand-held conductivity probe of hydrogels in equilibrium in deionized water (water gel) or PBS (ionic gel). B) Standard curve of conductivity vs. phosphate buffered saline solution concentration.

Figure S13. Hydrogel conductivity at different NaCl concentrations. Initial hydrogel conductivity at different NaCl concentrations and conductivity after reaching swelling equilibrium in PBS.

Figure S14. Network properties of PEUDAm prior to and after subcutaneous implantation in rats for 1, 2 or 4 weeks (n=8).

Figure S15. Circuit used to test the impact of hydrogel length on current. An oscilloscope measured the voltage across the sense resistor to determine the current.

Figure S16. In vivo assessment of injectable electrode in coronary vein (middle cardiac vein) of a porcine model. Host response after 2 weeks implantation in MCV using slices of the location proximal to the injection site, middle of the vein, and distal to the injection site (green arrow). The white arrow indicates the approximate location of the hydrogel in the vein. The response includes damage induced at the hydrogel injection site, with 1) moderate perivascular and interstitial fibrosis with focal replacement fibrosis (blue arrows), and 2) fibrosing epicarditis with foreign body giant cell reaction (red arrows). Mid branch of the hydrogel injection show 3) mild perivascular and interstitial fibrosis, 4) mild focal replacement fibrosis, and fibrosing epicarditis. Distal branch indicated 5) preserved myocardium with 6) only fibrosing epicarditis with focal extension into myocardium.

Figure S17. In vivo assessment of injectable electrode in coronary vein (anterior interventricular vein) of a porcine model (Animal 1). Host response after 4 weeks implantation in AIV using slices of the location proximal to the injection site, middle of the vein (indicated by V), and the control that is distal to the hydrogel. The response includes damage induced at the hydrogel injection site, with 1) moderate perivascular and interstitial fibrosis with focal replacement fibrosis (blue arrows), and 2) fibrosing epicarditis (red arrows) with foreign body giant cell reaction (blue arrows). Mid branch of the hydrogel injection show 3) mild perivascular and interstitial fibrosis, 4) mild focal replacement fibrosis, and fibrosing epicarditis. The control images at the distal branch indicated 5) preserved myocardium with 6) only fibrosing epicarditis with focal extension into myocardium.

Figure S18. In vivo assessment of injectable electrode in coronary vein (anterior interventricular vein) of a porcine model (Animal 2). Host response after 4 weeks implantation in AIV using slices of the location proximal to the injection site, middle of the vein (indicated by V), and the control from an alternate section of the anterior wall. The response includes damage induced at the hydrogel injection site, with 1) moderate perivascular and interstitial fibrosis with focal replacement fibrosis (blue arrows), and 2) fibrosing epicarditis with foreign body giant cell reaction (green arrows). Mid branch of the hydrogel injection show 3) moderate perivascular and interstitial (black arrows) fibrosis, and replacement fibrosis (blue arrows), 4) focal replacement fibrosis, and fibrosing epicarditis. The control images at an alternate section of the anterior wall indicated 5) 6) preserved myocardium

Figure S19. In vivo assessment of injectable electrode in coronary vein (anterior intraventricular vein) of a porcine model. Animal AIV 1. ECG tracings from lead 1 of the cardiac activity before injection of the hydrogel (black tracing) and 4 weeks after injection (blue tracing). The overall QRS morphology is preserved. Heart rate differs between the two recordings, but this is not indicative of a disease state.

Figure S20. In vivo assessment of injectable electrode in coronary vein (anterior intraventricular vein) of a porcine model. Animal AIV 2. ECG tracings from lead 1 of the cardiac activity before injection of the hydrogel (black tracing) and 4 weeks after injection (blue tracing). The overall QRS morphology is preserved.

Figure S21. In vivo assessment of injectable electrode in coronary vein (anterior intraventricular vein) of a porcine model. Animal AIV 3. ECG tracings from lead 1 of the cardiac activity before injection of the hydrogel (black tracing) and 4 weeks after injection (blue tracing). The overall QRS morphology is similar, with nearly identical vectors. Heart rate differs between the two recordings, but this is not indicative of a disease state.

Figure S22. In vivo assessment of injectable electrode in coronary vein (middle cardiac vein) of a porcine model. Animal MCV 1. ECG tracings from lead 1 of the cardiac activity before injection of the hydrogel (black tracing) and 2 weeks after injection (blue tracing). The overall QRS morphology is similar, with nearly identical vectors. Heart rate differs between the two recordings, but this is not indicative of a disease state. The pre-injection ECG tracing contains low frequency baseline drift which is not reflective of the model’s electrophysiology.

Figure S23. Capture thresholds for each acute porcine study (n=3). Figure S24. 12-lead ECG of the heart at sinus rhythm.

Figure S25. 12-lead ECG of unipolar pacing with a metal electrode placed directly on the left ventricular myocardium. Pacing artifacts can be seen as spikes in each of the channels. Pacing was performed at 120 bpm. Figure S26. 12-lead ECG of unipolar pacing with a small blob of hydrogel electrode placed directly on the left ventricular myocardium. Pacing artifacts can be seen as spikes in each of the channels. Pacing was performed at 120 bpm.

Figure S27. 12-lead ECG of unipolar pacing with a linear hydrogel electrode (∼4 cm) placed directly on the left ventricular myocardium close to the interventricular septum. Pacing artifacts can be seen as spikes in each of the channels. Pacing was performed at 120 bpm.

Figure S28. 12-lead ECG of unipolar pacing from hydrogel cured in situ in the AIV. Pacing artifacts can be seen as spikes in each of the channels. Pacing was performed at 120 bpm.

Figure S29. Animal02: 12-lead ECG of the heart at sinus rhythm.

Figure S30. Animal02: 12-lead ECG of unipolar pacing with a metal electrode placed directly on the left ventricular myocardium. Pacing artifacts can be seen as spikes in each of the channels. Pacing was performed at 120 bpm.

Figure S31. Animal02: 12-lead ECG of unipolar pacing with a small blob of hydrogel electrode placed directly on the left ventricular myocardium. Pacing artifacts can be seen as spikes in each of the channels. Pacing was performed at 120 bpm.

Figure S32. Animal02: 12-lead ECG of unipolar pacing with a linear hydrogel electrode (∼4 cm) placed directly on the left ventricular myocardium close to the interventricular septum. Pacing artifacts can be seen as spikes in each of the channels. Pacing was performed at 120 bpm.

Figure S33. Animal02: 12-lead ECG of unipolar pacing from hydrogel cured in situ in the AIV. Pacing artifacts can be seen as spikes in each of the channels. Pacing was performed at 120 bpm.

Figure S34. Animal03: 12-lead ECG of the heart at sinus rhythm.

Figure S35. Animal03: 12-lead ECG of unipolar pacing with metal electrode placed directly on the left ventricular myocardium. Pacing artifacts can be seen as spikes in each of the channels. Pacing was performed at 120 bpm.

Figure S36. Animal03: 12-lead ECG of unipolar pacing with a small blob of hydrogel electrode placed directly on the left ventricular myocardium. Pacing artifacts can be seen as spikes in each of the channels. Pacing was performed at 120 bpm.

Figure S37. Animal03: 12-lead ECG of unipolar pacing with a linear hydrogel electrode (∼4 cm) placed directly on the left ventricular myocardium close to the interventricular septum. Pacing artifacts can be seen as spikes in each of the channels. Pacing was performed at 120 bpm.

Figure S38. Animal03: 12-lead ECG of unipolar pacing from hydrogel cured in situ in the AIV. Pacing artifacts can be seen as spikes in each of the channels. Pacing was performed at 120 bpm.

Figure S39. Epicardial and endocardial electroanatomical mapping for animal AIV 1. The AIV hydrogel pacing is indicated by the blue circles.

Figure S40. Epicardial and endocardial electroanatomical mapping for animal AIV 2. Point pacing location is indicated by a small white circle, and the AIV hydrogel pacing is indicated by the blue circles. Points of ablation are marked by the red circles.

Figure S41. Epicardial and endocardial electroanatomical mapping for animal AIV 3. Point pacing location is indicated by a small white circle, and the AIV is indicated by the blue circles. Points of ablation are marked by the red circles

Table S1. Effect of incorporating NAGA on hydrogel gel fraction.

Table S2. Fibrous capsule thickness (units in 1m)

Table S3. Inflammation severity scoring

Table S4. High sensitivity Troponin I measurements.

Table S5. Capture thresholds (in mA) for each material and acute animal tested (n=3). NC = no capture.

Movie S1. Mixing of precursors solutions using a dual lumen catheter for dispensing. Press play to see the video. Movie S2. Mixing of precursors solutions when dispensed using a double barrel syringe with a mixing head. Press play to see the video.

Movie S3. Echocardiogram 2 weeks after injection on the MCV in a pig model. Echocardiograms show no evidence of regional wall motion abnormalities after two weeks. Press play to see the videos.

Movie S4. Echocardiogram prior and 4 weeks after injection on the AIV in a pig model. Echocardiograms show no evidence of regional wall motion abnormalities at four weeks compared to baseline imaging. Press play to see the videos.

Movie S5. Myocardium activation comparison after pacing with a single point (metal electrode) or with the hydrogel in the vein for Animal AIV 1. The AIV is indicated by the blue circles. Press play to see the animations. Movie S6. Myocardium activation comparison after pacing with a single point (metal electrode) or with the hydrogel in the vein for Animal AIV 2. A small green circle indicates point pacing location on the first image. Press play to see the animations.

## Acknowledgments

The authors are grateful for the contributions of the THI Animal Lab staff, particularly Dr. Abdelmotagaly Elgalad and Dr. Angel Moctezuma-Ramirez, as well as the THI Pathology Lab, particularly Dr. Pamela Potts and Dr. Ana Segura. We also thank Dr. Lynd (The University of Texas at Austin) for the use of the EIS and the Center for Electrochemistry and Texas Materials Institute for the use of the four-point probe and rheometer, respectively.

## Funding

• Ford Pre-Doctoral Fellowship, administered by the National Academy of Science, Engineering and Medicine (GJRR)

• Ford Dissertation Fellowship, administered by the National Academy of Science, Engineering and Medicine (GJRR)

• Office of Vice President for Research, The University of Texas at Austin, (ECH)

• The Roderick D. MacDonald Research Fund Award 19RDM004 (MR)

• The Sultan Qaboos Chair in Cardiology at the St. Luke’s Foundation (MR)

## Author contributions

Each author’s contribution(s) to the paper should be listed [we encourage you to follow the CRediT model]. Each CRediT role should have its own line, and there should not be any punctuation in the initials.

• Conceptualization: ECH, MR
• Methodology: ECH, MR, GJRR, MJ, AP, SB, MW, MA
• Investigation: GJRR, MW, AP, MJ, ECH
• Visualization: GJRR, CW, MJ
• Supervision: ECH, MR
• Writing—original draft: GJRR, MW, AP, MJ, ECH
• Writing—review & editing: ECH, MR

## Data and materials availability

All data are available in the main text or the supplementary materials. All the raw data will be uploaded into The University of Texas at Austin UTBox cloud storage. Access to the data will be provided upon request.

## References and Notes

1. R. M. John et al., Ventricular arrhythmias and sudden cardiac death. The Lancet 380, 1520–1529 (2012).

2. R. Derksen et al., Tissue discontinuities affect conduction velocity restitution: a mechanism by which structural barriers may promote wave break. Circulation 108, 882–888 (2003).

3. E. Patterson, B. J. Scherlag, E. J. Berbari, R. Lazzara, Slow conduction through an arc of block: A basis for arrhythmia formation postmyocardial infarction. J Cardiovasc Electrophysiol 28, 1203–1212 (2017).

4. Z. Qu, J. N. Weiss, Mechanisms of ventricular arrhythmias: from molecular fluctuations to electrical turbulence. Annu Rev Physiol 77, 29–55 (2015).

5. W. G. Stevenson, J. N. Weiss, I. Wiener, K. Nademanee, Slow conduction in the infarct scar: relevance to the occurrence, detection, and ablation of ventricular reentry circuits resulting from myocardial infarction. Am Heart J 117, 452–467 (1989).

6. H. J. Wellens, P. Brugada, J. Farre, Ventricular arrhythmias: mechanisms and actions of antiarrhythmic drugs. Am Heart J 107, 1053–1057 (1984).

7. E. S. Williams, M. N. Viswanathan, Current and emerging antiarrhythmic drug therapy for ventricular tachycardia. Cardiol Ther 2, 27–46 (2013).

8. R. Cardoso, F. R. Assis, A. D’Avila, Endo-epicardial vs endocardial-only catheter ablation of ventricular tachycardia: A meta-analysis. J Cardiovasc Electrophysiol, (2019).

9. H. Versteeg, D. A. Theuns, R. A. Erdman, L. Jordaens, S. S. Pedersen, Posttraumatic stress in implantable cardioverter defibrillator patients: the role of pre-implantation distress and shocks. International journal of cardiology 146, 438–439 (2011).

10. J. Seuring, F. M. Bayer, K. Huber, S. Agarwal, Upper Critical Solution Temperature of Poly(N-acryloyl glycinamide) in Water: A Concealed Property. Macromolecules 45, 374–384 (2012).

11. S. Ramadan, N. Paul, H. E. Naguib, Standardized static and dynamic evaluation of myocardial tissue properties. Biomedical Materials 12, 025013 (2017).

12. M. S. Amzulescu et al., Myocardial strain imaging: review of general principles, validation, and sources of discrepancies. European Heart Journal - Cardiovascular Imaging 20, 605–619 (2019).

13. X. Guo, Y. Liu, G. S. Kassab, Diameter-dependent axial prestretch of porcine coronary arteries and veins. Journal of applied physiology 112, 982–989 (2012).

14. M. Whitely et al., Improved in situ seeding of 3D printed scaffolds using cell-releasing hydrogels. Biomaterials 185, 194–204 (2018).

15. M. Solazzo, F. J. O’Brien, V. Nicolosi, M. G. Monaghan, The rationale and emergence of electroconductive biomaterial scaffolds in cardiac tissue engineering. APL bioengineering 3, 041501 (2019).

16. D. M. Pedrotty et al., Three-dimensional printed biopatches with conductive ink facilitate cardiac conduction when applied to disrupted myocardium. Circulation: Arrhythmia and Electrophysiology 12, e006920 (2019).

17. M. Potse, B. Dubé, A. Vinet, Cardiac anisotropy in boundary-element models for the electrocardiogram. Medical & biological engineering & computing 47, 719–729 (2009).

18. J. G. Webster, I. E. i. Medicine, B. Society, Design of Cardiac Pacemakers. (IEEE, 1995).

19. A. Ido, N. Hasebe, H. Matsuhashi, K. Kikuchi, Coronary sinus occlusion enhances coronary collateral flow and reduces subendocardial ischemia. American Journal of Physiology-Heart and Circulatory Physiology 280, H1361–H1367 (2001).

20. B. Merkely et al., Chronic implantation of intravascular cardioverter defibrillator in a canine model: device stability, vascular patency, and anchor histology. Pacing and Clinical Electrophysiology 36, 1251–1258 (2013).

21. J. J. Thornton, D. E. Gregg, Effect of chronic cardiac venous occlusion on coronary arterial and cardiac venous hemodynamics. American Journal of Physiology-Legacy Content 128, 179–184 (1939).

22. M. Meesmann et al., Selective perfusion of ischemic myocardium during coronary venous retroinjection: a study of the causative role of venoarterial and venoventricular pressure gradients. Journal of the American College of Cardiology 10, 887–897 (1987).

23. S. S. Ponnusamy et al., Left bundle branch pacing: a comprehensive review. Journal of cardiovascular electrophysiology 31, 2462–2473 (2020).

24. N. Ali et al., His bundle pacing: a new frontier in the treatment of heart failure. Arrhythmia & electrophysiology review 7, 103 (2018).

25. S. K. Mulpuru, M. Madhavan, C. J. McLeod, Y.-M. Cha, P. A. Friedman, Cardiac Pacemakers: Function, Troubleshooting, and Management. Journal of the American College of Cardiology 69, 189–210 (2017).

26. M. Jastrzębski, ECG and Pacing Criteria for Differentiating Conduction System Pacing from Myocardial Pacing. Arrhythmia & Electrophysiology Review *2021*;10(*3*):172*–*80., (2021).

27. M. Jastrzębski et al., Physiology-based electrocardiographic criteria for left bundle branch capture. Heart Rhythm 18, 935–943 (2021).

28. M.-Y. Gao et al., Electrocardiographic morphology during left bundle branch area pacing: Characteristics, underlying mechanisms, and clinical implications. Pacing and Clinical Electrophysiology 43, 297–307 (2020).

29. M. J. Janse, A. L. Wit, Electrophysiological mechanisms of ventricular arrhythmias resulting from myocardial ischemia and infarction. Physiological Reviews 69, 1049–1169 (1989).

30. G. Camci-Unal, N. Annabi, M. R. Dokmeci, R. Liao, A. Khademhosseini, Hydrogels for cardiac tissue engineering. NPG Asia Materials 6, e99 (2014).

31. J. Prager et al., Stiffness-matched biomaterial implants for cell delivery: clinical, intraoperative ultrasound elastography provides a ‘target’ stiffness for hydrogel synthesis in spinal cord injury. Journal of Tissue Engineering 11, 204173142093480 (2020).

32. C. Yu, F. Yao, J. Li, Rational design of injectable conducting polymer-based hydrogels for tissue engineering. Acta Biomaterialia, (2021).

33. M. Shin, K. H. Song, J. C. Burrell, D. K. Cullen, J. A. Burdick, Injectable and Conductive Granular Hydrogels for 3D Printing and Electroactive Tissue Support. Advanced Science 6, 1901229 (2019).

34. A. Zhbanov, S. Yang, Effects of Aggregation on Blood Sedimentation and Conductivity. PLOS ONE 10, e0129337 (2015).

35. M. Zyl et al., Injectable conductive hydrogel restores conduction through ablated myocardium. Journal of Cardiovascular Electrophysiology 31, 3293–3301 (2020).

36. J. Huang, X. Jiang, Injectable and Degradable pH-Responsive Hydrogels via Spontaneous Amino–Yne Click Reaction. ACS Applied Materials & Interfaces 10, 361–370 (2018).

37. J. L. Ifkovits et al., Injectable hydrogel properties influence infarct expansion and extent of postinfarction left ventricular remodeling in an ovine model. Proceedings of the National Academy of Sciences 107, 11507–11512 (2010).

38. J. Shi et al., Cell-compatible hydrogels based on a multifunctional crosslinker with tunable stiffness for tissue engineering. Journal of Materials Chemistry 22, 23952–23962 (2012).

39. N. J. Steinmetz, E. A. Aisenbrey, K. K. Westbrook, H. J. Qi, S. J. Bryant, Mechanical loading regulates human MSC differentiation in a multi-layer hydrogel for osteochondral tissue engineering. Acta biomaterialia 21, 142–153 (2015).

40. J. S. Temenoff et al., Thermally cross-linked oligo (poly (ethylene glycol) fumarate) hydrogels support osteogenic differentiation of encapsulated marrow stromal cells in vitro. Biomacromolecules 5, 5–10 (2004).

41. T. S. Wilems et al., Effects of free radical initiators on polyethylene glycol dimethacrylate hydrogel properties and biocompatibility. Journal of Biomedical Materials Research Part A 105, 3059–3068 (2017).

42. S. A. Young et al., Mechanically resilient injectable scaffolds for intramuscular stem cell delivery and cytokine release. Biomaterials 159, 146–160 (2018).

43. S. M. Dorsey et al., MRI evaluation of injectable hyaluronic acid-based hydrogel therapy to limit ventricular remodeling after myocardial infarction. Biomaterials 69, 65–75 (2015).

44. J. H. Traverse et al., First-in-Man Study of a Cardiac Extracellular Matrix Hydrogel in Early and Late Myocardial Infarction Patients. JACC: Basic to Translational Science 4, 659–669 (2019).

45. S. Liang et al., Paintable and rapidly bondable conductive hydrogels as therapeutic cardiac patches. Advanced Materials 30, 1704235 (2018).

46. A. Navaei et al., Gold nanorod-incorporated gelatin-based conductive hydrogels for engineering cardiac tissue constructs. Acta biomaterialia 41, 133–146 (2016).

47. S. Buchan, R. Kar, M. John, A. Post, M. Razavi, Electrical Stimulation for Low-Energy Termination of Cardiac Arrhythmias: a Review. Cardiovascular Drugs and Therapy, (2021).

48. L. Jin, J. Wang, B. Song, X. Wu, Z. Fang, in 2015 37th Annual International Conference of the IEEE Engineering in Medicine and Biology Society (EMBC). (2015), pp. 5688–5691.

49. X. Zheng et al., Reduction of Atrial Defibrillation Threshold With an Interatrial Septal Electrode. Circulation 102, 2659–2664 (2000).

50. X. Zheng, M. E. Benser, G. P. Walcott, R. E. Ideker, Right Atrial Septal Electrode for Reducing the Atrial Defibrillation Threshold. Circulation 104, 1066–1070 (2001).

51. A. Moreno et al., Wide-area low-energy surface stimulation of large mammalian ventricular tissue. Scientific reports 9, 1–11 (2019).

52. S. K. Padala, K. A. Ellenbogen, Left bundle branch pacing is the best approach to physiological pacing. Heart Rhythm *O2* 1, 59–67 (2020).

53. L. I. B. Heckman et al., Electrical characteristics of deep septal vs. left bundle branch (area) pacing. European Heart Journal 41, (2020).

54. A. Di Marco et al., Deep septal pacing to upgrade patients with pacing-induced cardiomyopathy. HeartRhythm Case Reports.

55. S. J. Nelson, J. J. Creechley, M. E. Wale, T. J. Lujan, Print-A-Punch: A 3D printed device to cut dumbbell-shaped specimens from soft tissue for tensile testing. Journal of Biomechanics 112, 110011 (2020).

56. N. Huebsch et al., Automated Video-Based Analysis of Contractility and Calcium Flux in Human-Induced Pluripotent Stem Cell-Derived Cardiomyocytes Cultured over Different Spatial Scales. Tissue Engineering Part C: Methods 21, 467–479 (2015).

57. X. Lian et al., Directed cardiomyocyte differentiation from human pluripotent stem cells by modulating Wnt/β-catenin signaling under fully defined conditions. Nature Protocols 8, 162–175 (2013).

