## Supplementary Information for "Towards prevention of re-entrant arrhythmias: Injectable hydrogel electrodes enable direct capture of previously inaccessible cardiac tissue"

Movie S6. Myocardium activation comparison after pacing with a single point (metal electrode) or with the hydrogel in the vein for Animal AIV 2. A small green circle indicates point pacing location on the first image. Press play to see the animations.

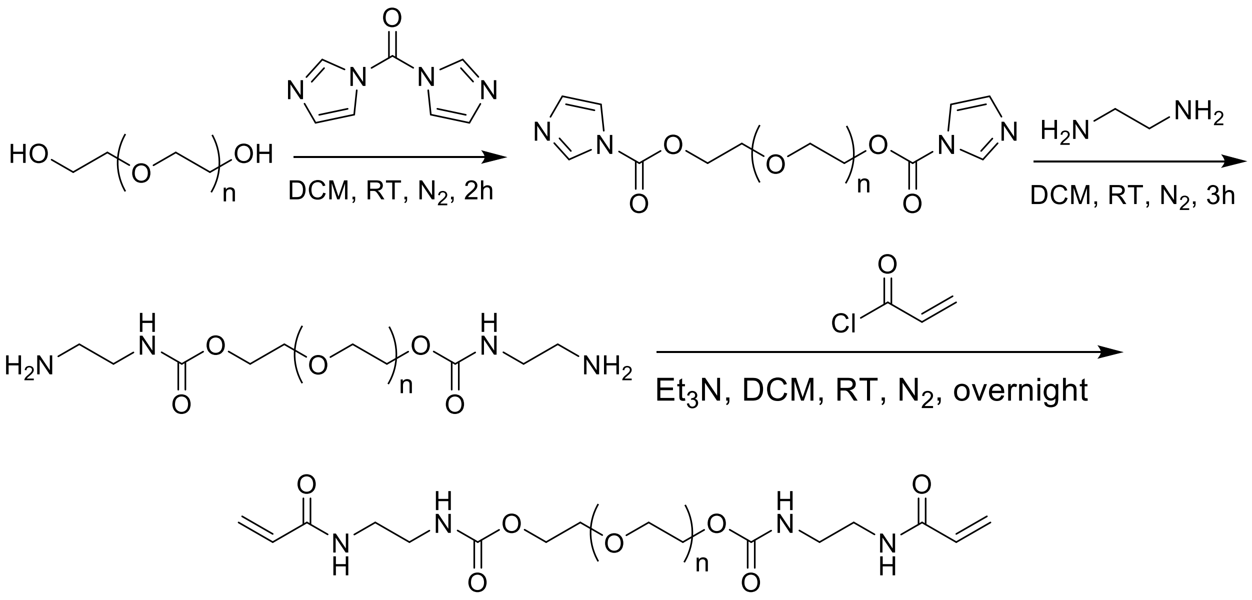


**Figure S1. Synthetic route for PEUDAm.**

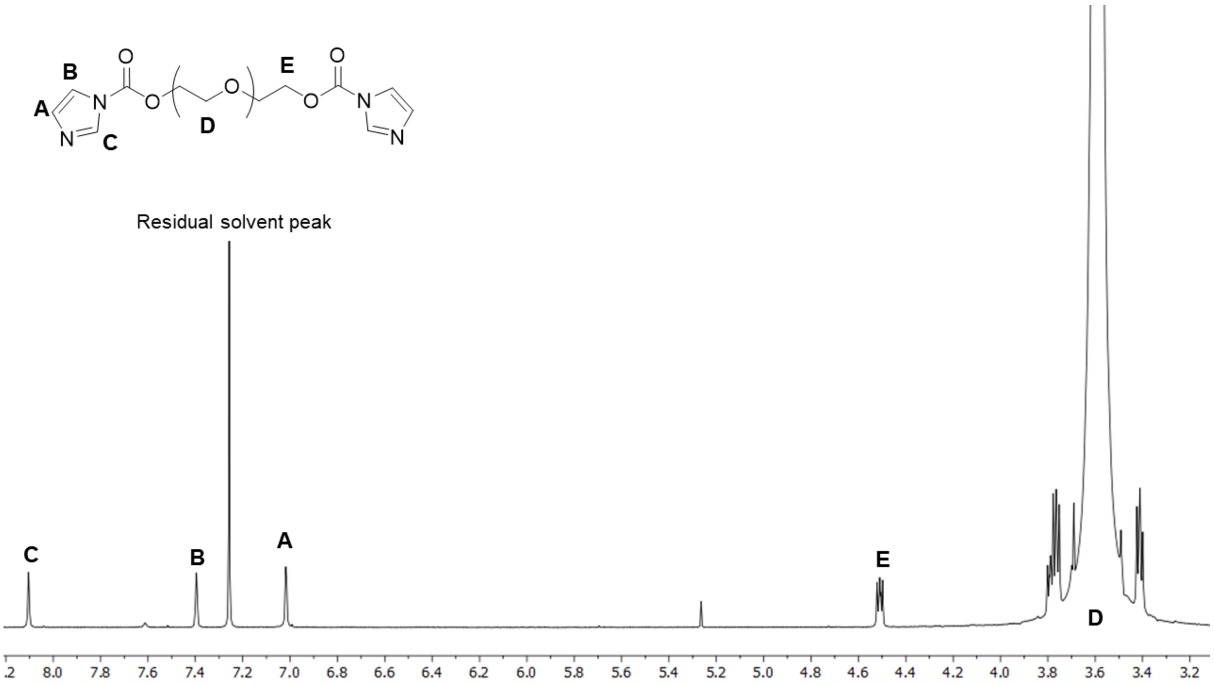

**Figure S2. H[1] NMR Spectra of PEG-CDI.**

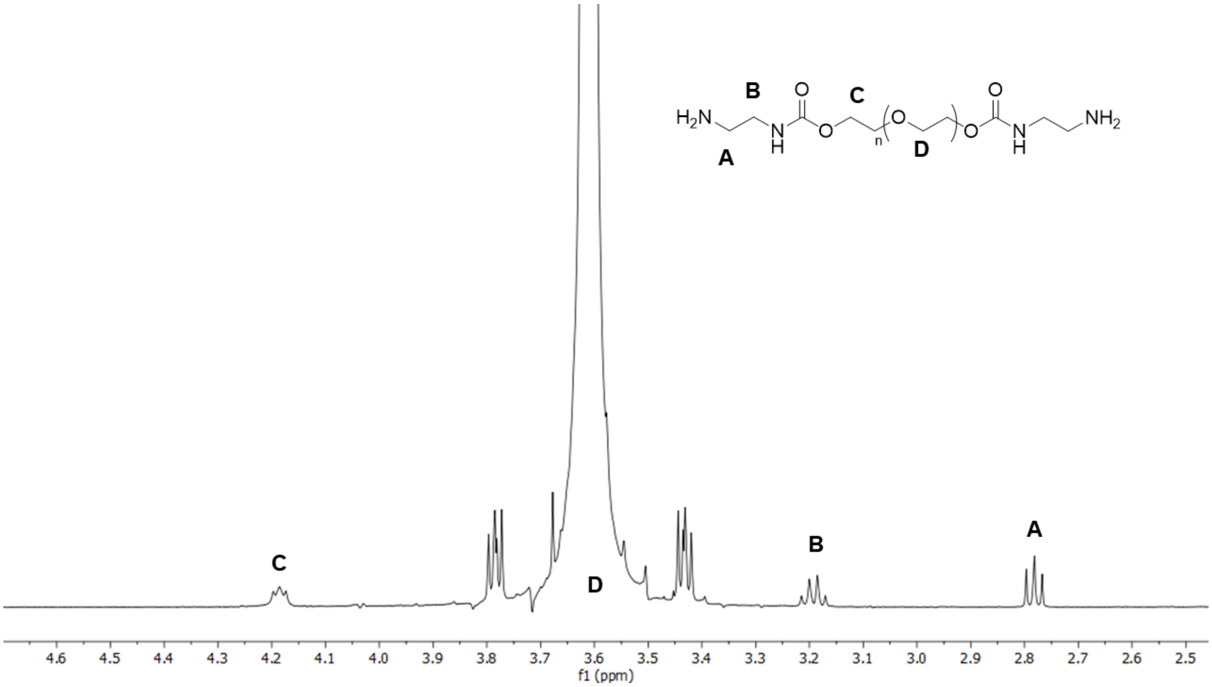


**Figure S3. H[1] NMR Spectra of PEG-EDA.**

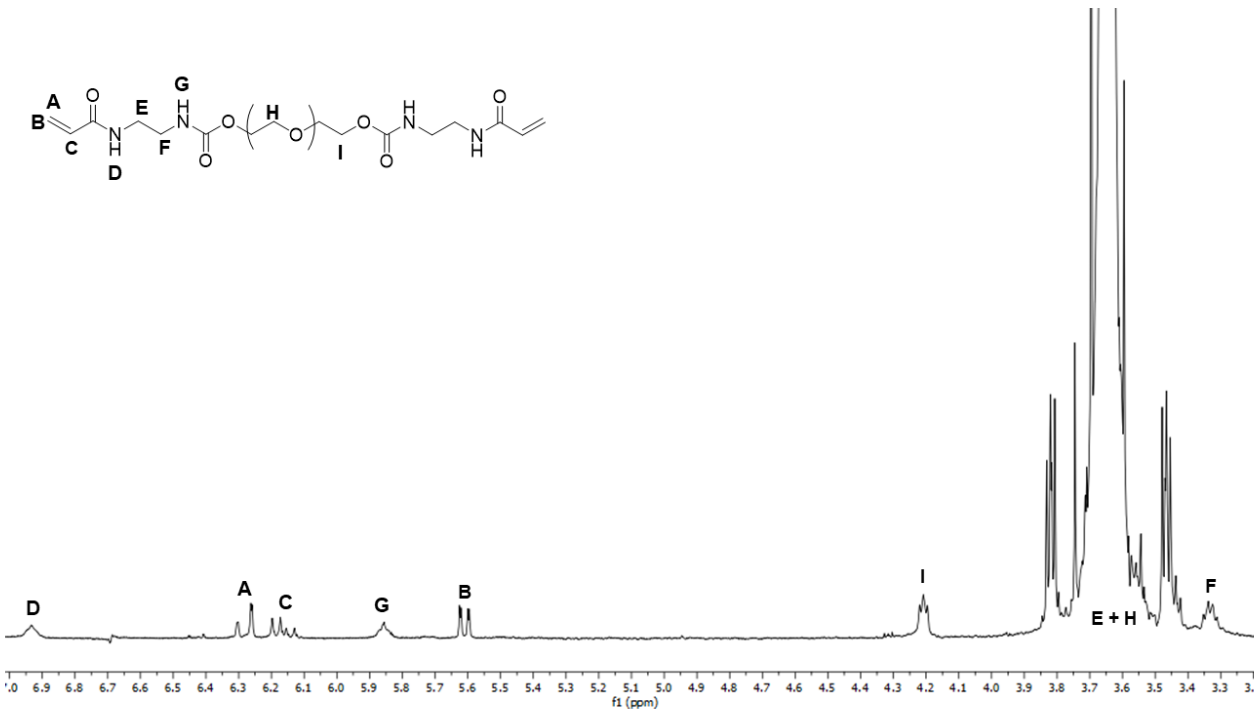


**Figure S4. H[1] NMR Spectra of PEUDAm.**

| | Mn (g/mol) | Mw (g/mol) | PDI |
|---|---|---|---|
| Peak 1 | 19960 ± 43 | 21530 ± 24 | 1.08 ± 0.00 |
| Peak 2 | 49340 ± 220 | 50600 ± 430 | 1.03 ± 0.00 |

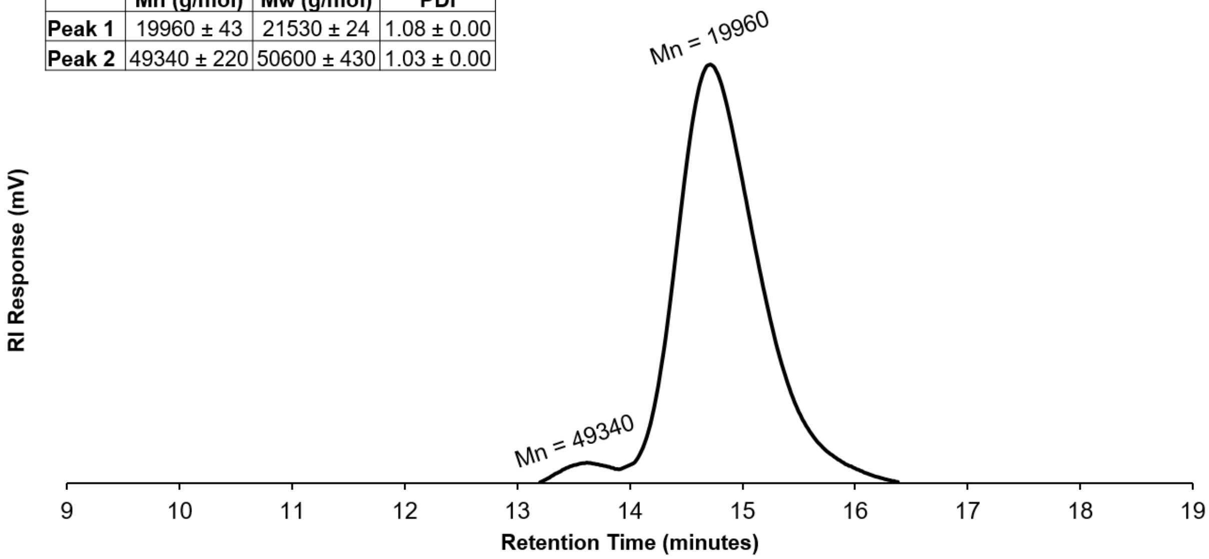


**Figure S5. Gel Permeation Chromatogram of PEUDAm macromer.**

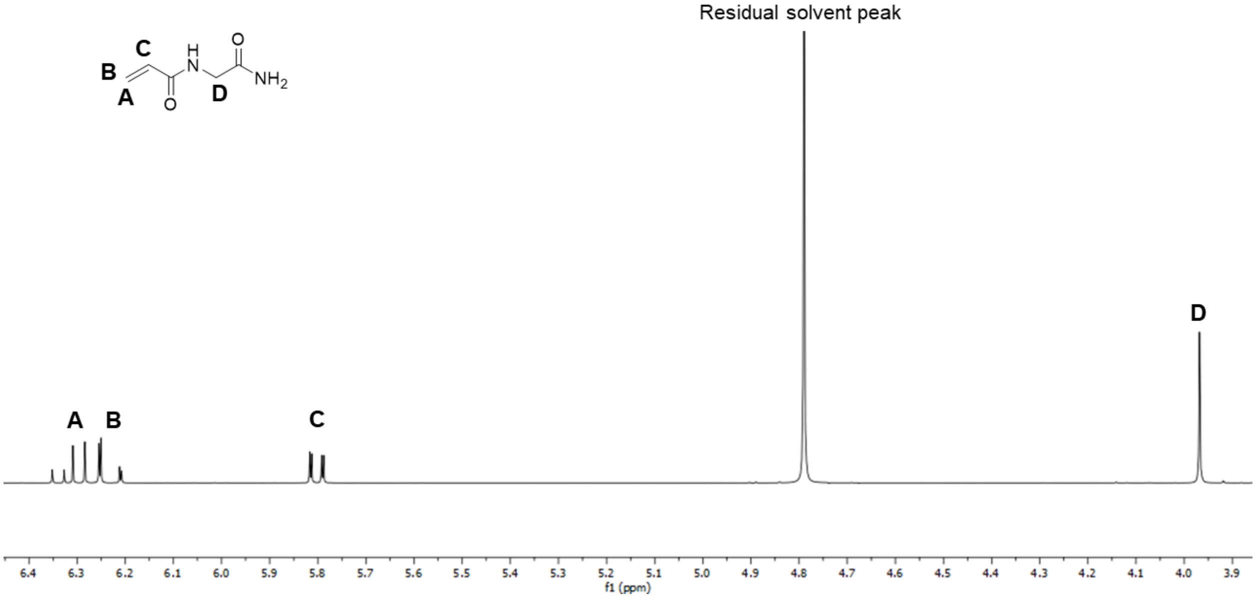


**Figure S6. H[1] NMR spectra of NAGA.**

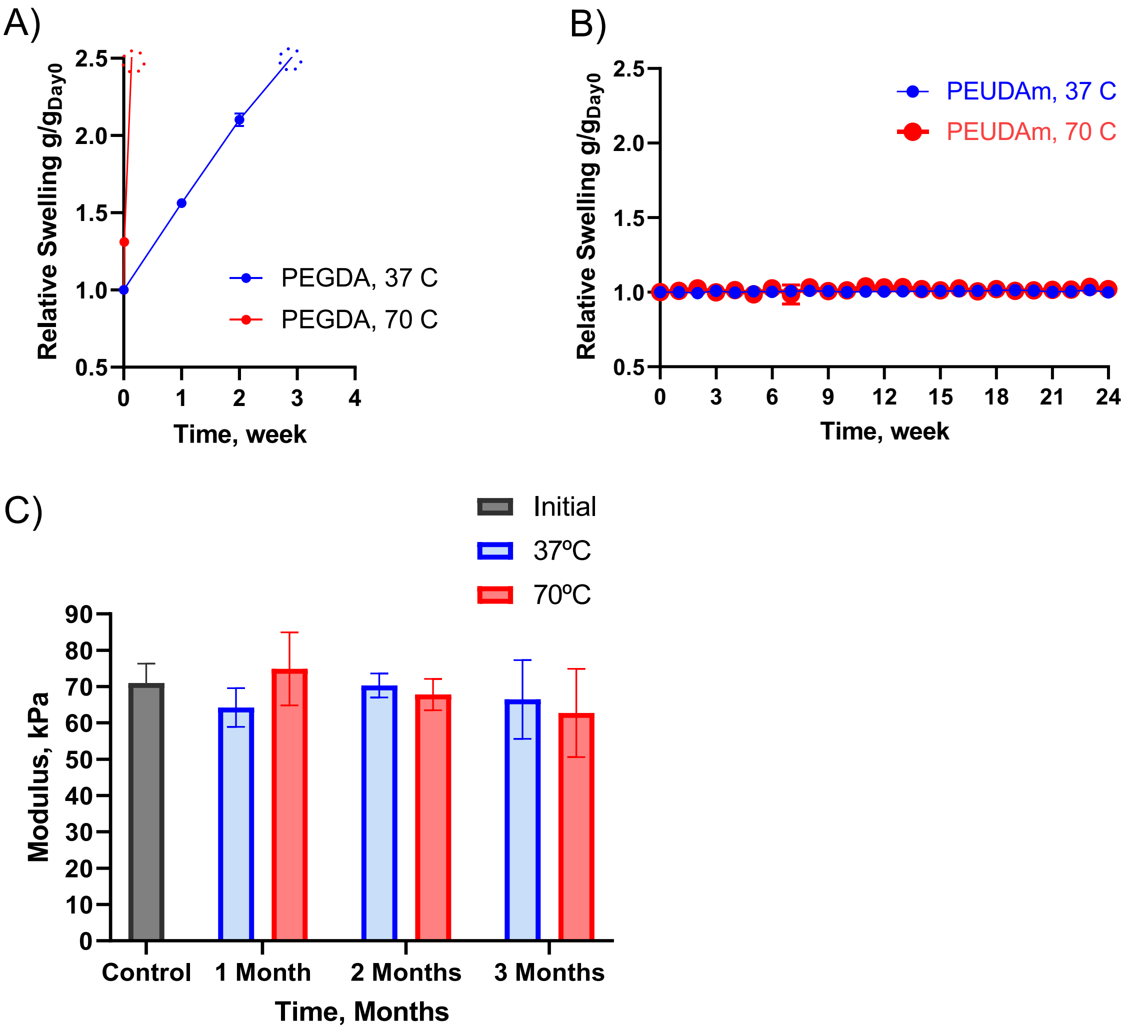


**Figure S7. PEUDAm stability under hydrolytic conditions.** PEGDA 20kDa, 20% + 1% NAGA was used as a positive control to assess improved biostability of PEUDAm 20kDa, 20% + 1% NAGA gels. Hydrogel swelling of (A) PEGDA and (B) PEUDAm in PBS was recorded as a function of time at 37ºC and under accelerated conditions (70ºC). Means and standard deviations are presented (n = 4). In most cases, the standard deviation is smaller than the bullets' size and was not displayed by the software**.** Storage modulus of PEUDAm hydrogels (C) under 1% strain at 0.5 Hz (compression) (n = 4).

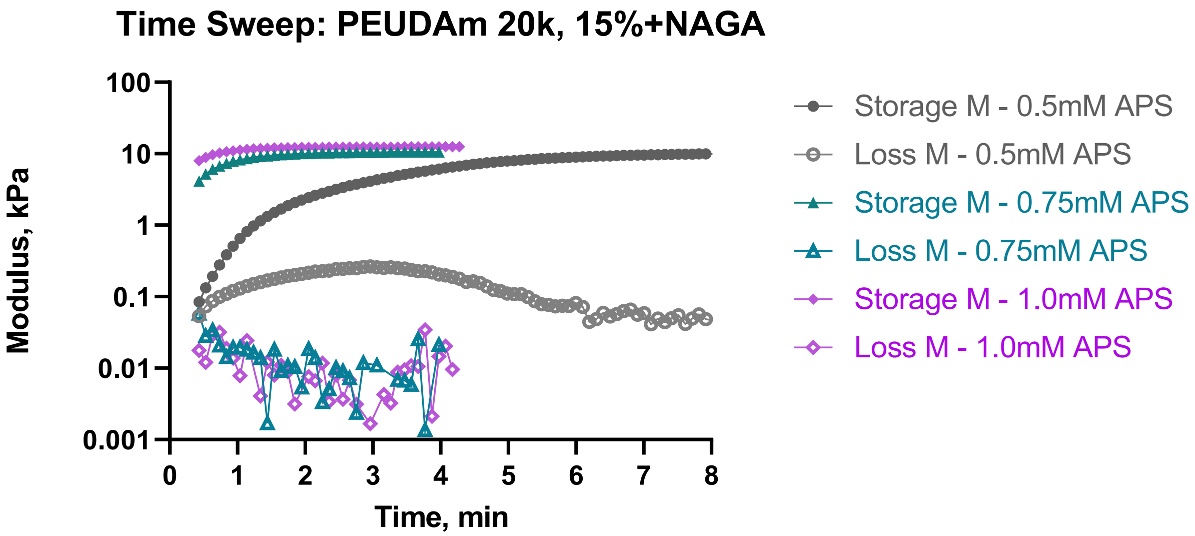


**Figure S8. Rheological data: Time sweep of PEUDAm hydrogels.** Time sweep of PEUDAm 20kDa, 15% cured at a concentration at 0.5, 0.75, and 1.0mM APS. The concentration of IG was twice the APS concentration: 1.0, 1.5, and 2.0, respectively.

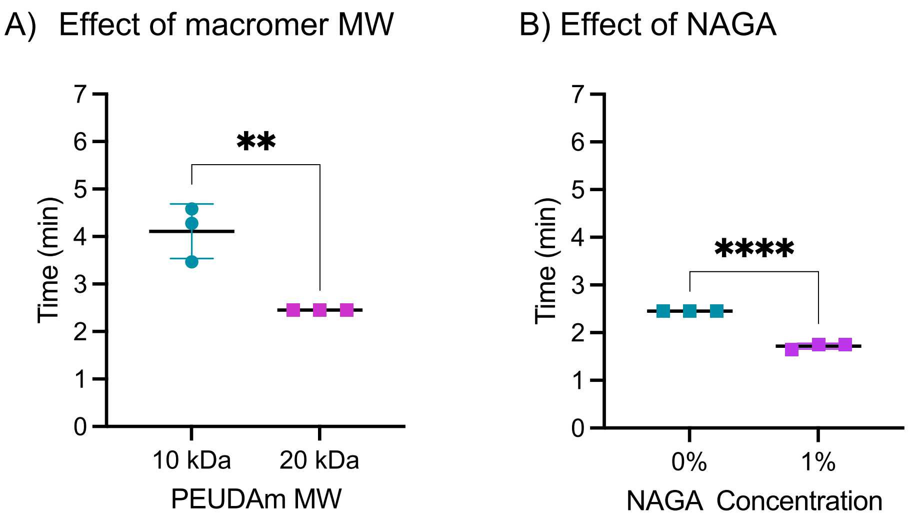


**Figure S9. Effect on time to complete network formation.** A) Effect of PEUDAm molecular weight (at 15 w/w%). B) Effect of adding 1% NAGA (to a PEUDAm 20k, 15% solution). Significant differences between groups are marked as follows: ** ($p < 0.01$), **** ($p < 0.0001$).

**Table S1.** Effect of incorporating NAGA on hydrogel gel fraction.

| Polymer formulation | Gel fraction, % |
|---|---|
| PEUDAm 20kDa 15% | 71.1 ± 1.4 |
| PEUDAm 20kDa 15% + 1%NAGA | 93.5 ± 1.1 |

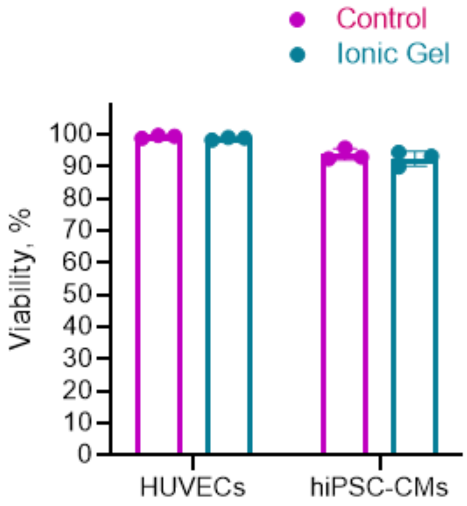


| | HUVECs | hiPSC-CMs |
|---|---|---|
| Control group viability, % | 99.3 ± 0.5 | 93.7 ± 1.8 |
| Ionic Gel group viability, % | 98.7 ± 0.4 | 92.5 ± 2.3 |
| Statistical differences (p-value) | No (0.8740) | No (0.5711) |

**Figure S10. Cytocompatibility by indirect contact with the hydrogels.** Percentage of HUVECs and hiPSC-cardiomyocytes viability 1 day after continuous exposure to hydrogel in comparison with TCPS. All data represents average ± standard deviation for n = 3 wells (x 3 location per well). There was no statistical differences when comparing to the TCPS control

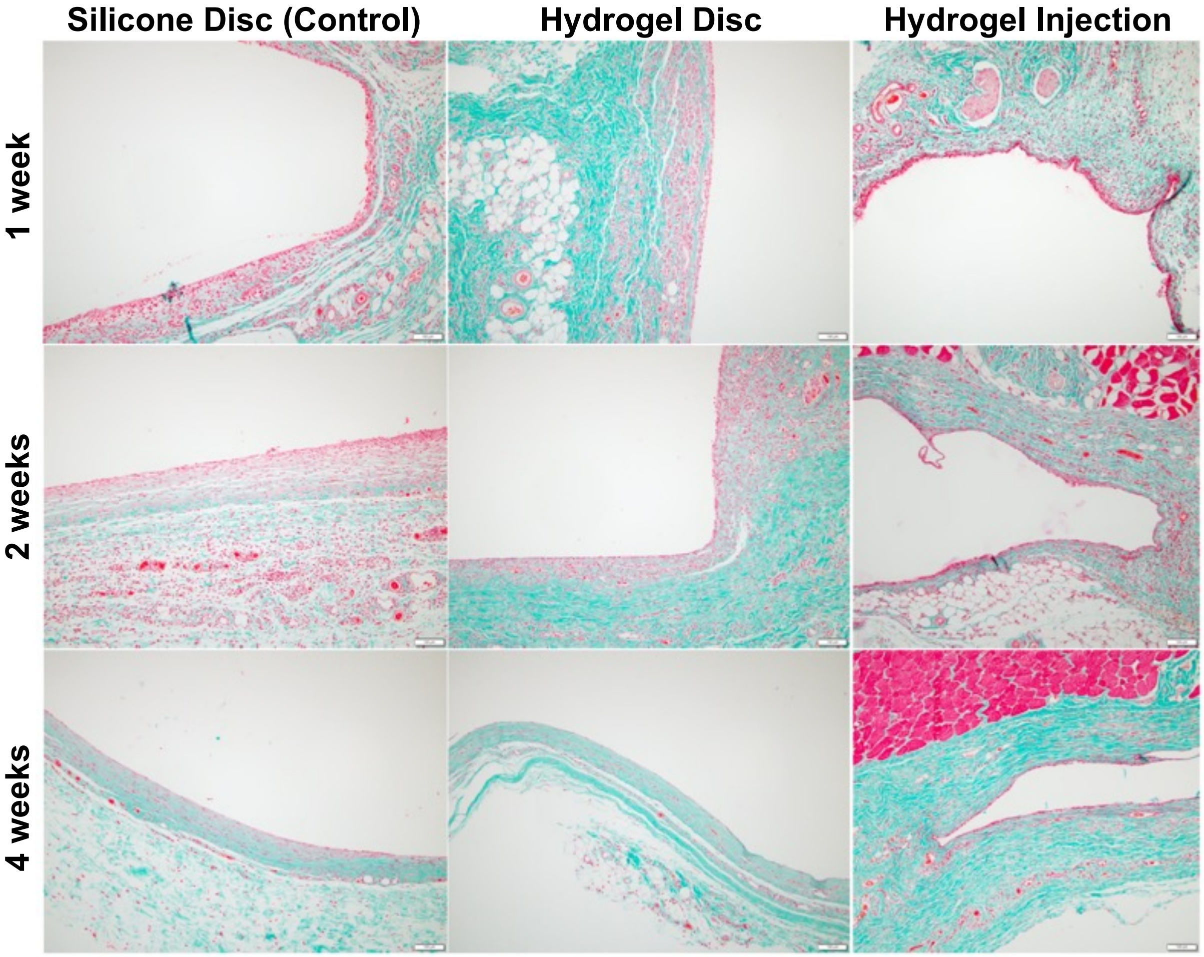


**Figure S11. Biocompatibility of PEUDAm hydrogel.** Representative higher magnification images showing the host response and evolution of a thin fibrous cap around the implanted hydrogel discs. (A, B) At 1 week post-implant, the interface between the hydrogel disc and surrounding tissue is composed mostly of macrophages, fibroblasts and capillaries (neovascularization, arrows) within a background of incipient extracellular matrix deposition. (C, D) By 4 weeks post-implant, a hypocellular fibrous capsule with increased collagen content has developed. Note the interface has remained of similar thickness (double headed arrows). Stains: H&E (left column) Masson's trichrome (right column). Scale bar = 20 μm.

**Table S2. Fibrous capsule thickness.** (Units in μm)

| Animal ID | Accession No. | Silicone Disc | | Hydrogel Disc | |
|---|---|---|---|---|---|
| | | By Group | By Animal | By Group | By Animal |
| *1 week* | | | | | |
| 7373 | 21-147 | | 53.7 ± 18.9 | | 82.3 ± 22.4 |
| 7374 | 21-148 | 50 ± 27.7 | 41.1 ± 7.1 | 64.6 ± 33.1 | 54.9 ± 18.3 |
| 7375 | 21-149 | | 53 ± 37.2 | | 54.6 ± 48.7 |
| *2 weeks* | | | | | |
| 7376 | 21-151 | | 131.2 ± 43.6 | | n/a |
| 7377 | 21-152 | 131.2 ± 43.6 | n/a | 83.2 ± 21.9 | 83.6 ± 21.6 |
| 7378 | 21-153 | | n/a | | 82.8 ± 23.1 |
| *4 weeks* | | | | | |
| 7379 | 21-162 | | 93.2 ± 36.6 | | 66.8 ± 26.5 |
| 7380 | 21-163 | 83.9 ± 28.5 | 73.1 ± 17.8 | 68.1 ± 21.8 | n/a |
| 7381 | 21-164 | | 80.5 ± 18 | | 66 ± 10.5 |

All values (mean ± SD)
n/a = measurements not taken due to histologic artifact

**Table S3. Inflammation severity scoring**

| | Silicone Disc (Control) | | | | | | | | |
|---|---|---|---|---|---|---|---|---|---|
| | 1 week | | | 2 weeks | | | 4 weeks | | |
| | R1 | R2 | R3 | R4 | R5 | R6 | R7 | R8 | R9 |
| Polymorphonuclear Leukocytes | - | -/+ | -/+ | + | -/+ | -/+ | - | -/+ | + |
| Macrophages | ++ | ++ | + | + | + | + | 1 | + | + |
| Lymphocytes | + | + | + | -/+ | -/+ | - | - | -/+ | -/+ |
| Fibroblasts | ++ | ++ | + | + | + | + | + | -/+ | + |
| Multinucleated giant cells | -/+ | -/+ | -/+ | -/+ | - | - | - | - | - |
| Vascularization | + | ++ | + | + | + | -/+ | - | -/+ | -/+ |

| | Hydrogel Disc | | | | | | | | |
|---|---|---|---|---|---|---|---|---|---|
| | 1 week | | | 2 weeks | | | 4 weeks | | |
| | R1 | R2 | R3 | R4 | R5 | R6 | R7 | R8 | R9 |
| Polymorphonuclear Leukocytes | -/+ | -/+ | - | -/+ | -/+ | - | - | * | - |
| Macrophages | ++ | ++ | ++ | + | + | + | -/+ | * | -/+ |
| Lymphocytes | + | + | + | - | -/+ | - | - | * | - |
| Fibroblasts | ++ | ++ | ++ | + | + | + | -/+ | * | -/+ |
| Multinucleated giant cells | -/+ | -/+ | -/+ | - | -/+ | - | -/+ | * | -/+ |
| Vascularization | ++ | ++ | ++ | + | + | + | -/+ | * | -/+ |

| | Hydrogel Injection | | | | | | | | |
|---|---|---|---|---|---|---|---|---|---|
| | 1 week | | | 2 weeks | | | 4 weeks | | |
| | R1 | R2 | R3 | R4 | R5 | R6 | R7 | R8 | R9 |
| Polymorphonuclear Leukocytes | * | * | - | - | - | - | - | * | - |
| Macrophages | * | * | + | ++ | + | + | + | * | + |
| Lymphocytes | * | * | -/+ | -/+ | -/+ | + | -/+ | * | -/+ |
| Fibroblasts | * | * | + | + | + | + | + | * | + |
| Multinucleated giant cells | * | * | - | - | - | - | - | * | - |
| Vascularization | * | * | + | -/+ | -/+ | -/+ | -/+ | * | - |

| Ranking | Absent | Occasional | Present to Markedly Present | | |
|---|---|---|---|---|---|
| | - | -/+ | + | ++ | +++ |

* Not able to recover specimens from the rats

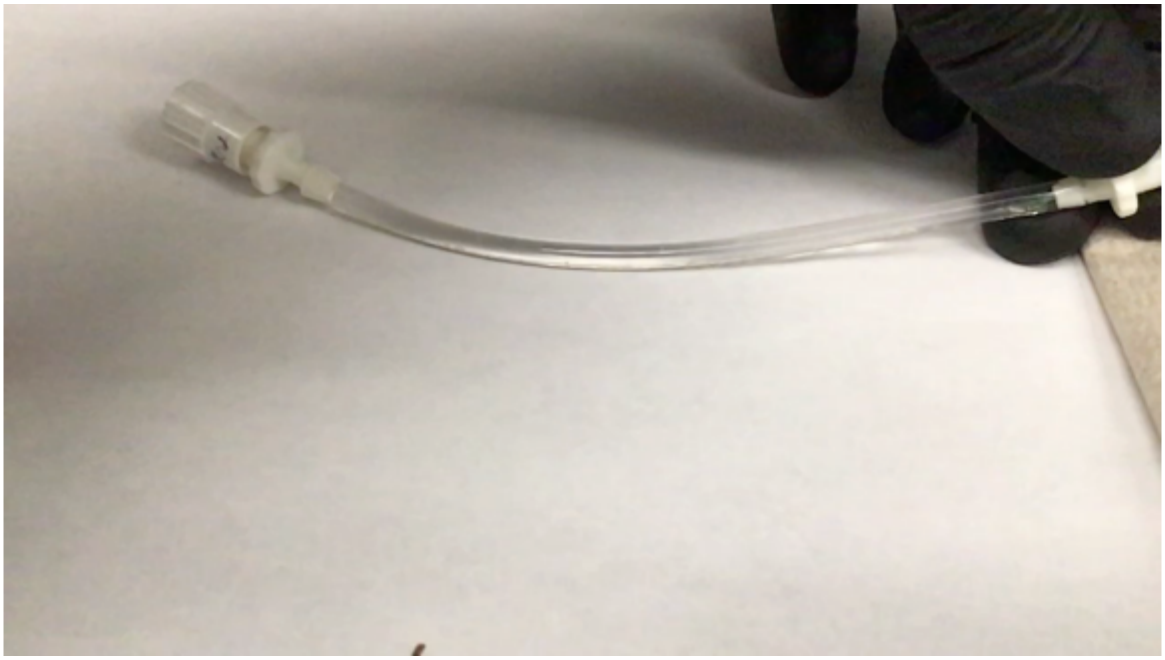
**Movie S1. Mixing of precursors solutions using a dual lumen catheter for dispensing.** Press play to see the video.

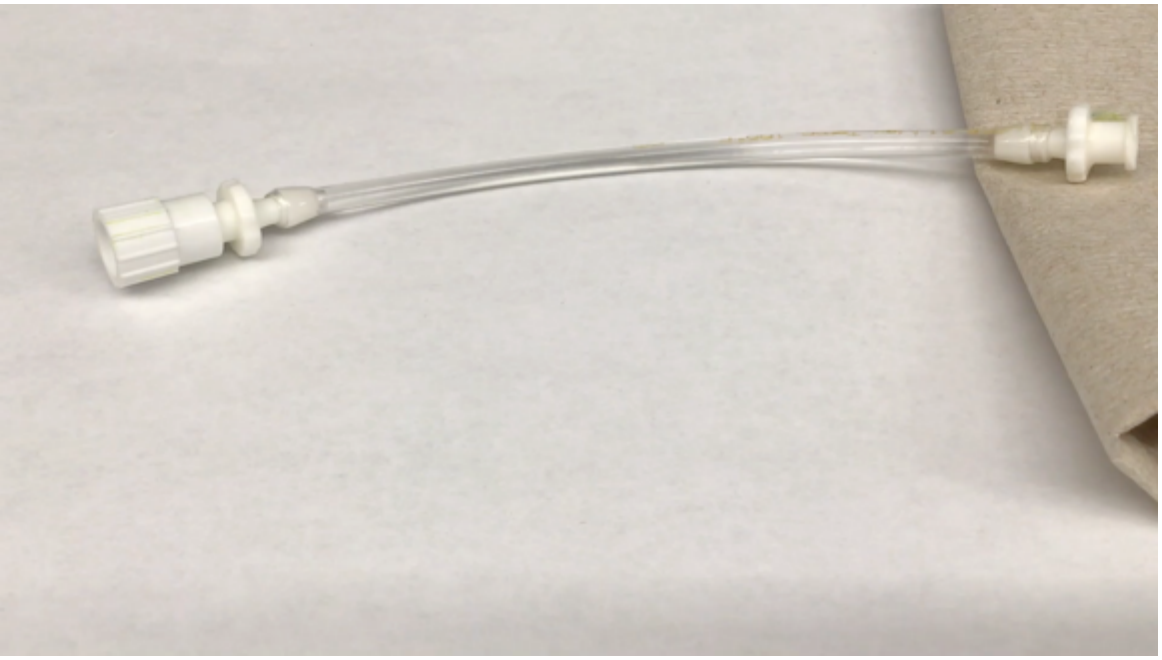

**Movie S2. Mixing of precursors solutions when dispensed using a double barrel syringe with a mixing head.** Press play to see the video.

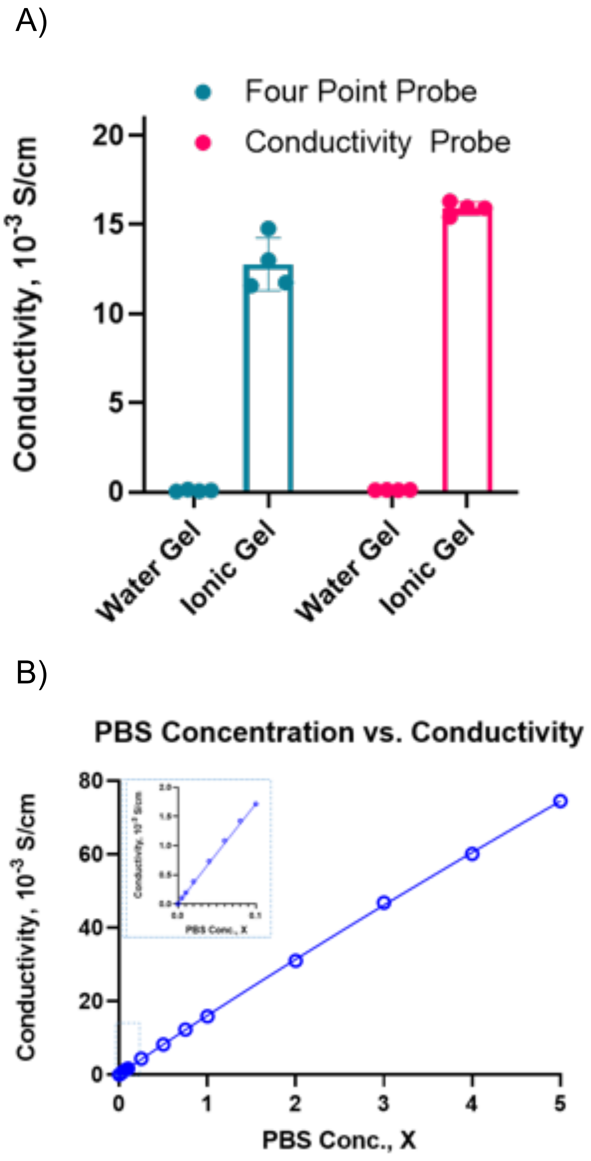


**Figure S12. Conductivity of hydrogel determined by 4-point probe and conductivity probe.** A) Comparison of hydrogels conductivity using the four-point probe and the hand-held conductivity probe of hydrogels in equilibrium in deionized water (water gel) or PBS (ionic gel). B) Standard curve of conductivity vs. phosphate buffered saline solution concentration.

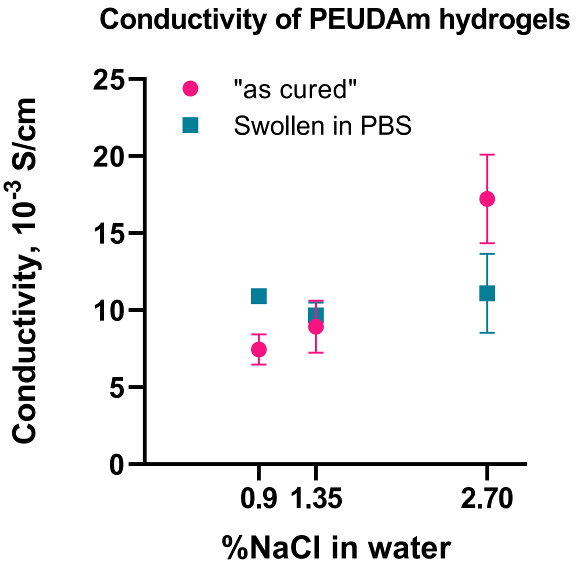


**Figure S13. Hydrogel conductivity at different NaCl concentrations.** Initial hydrogel conductivity at different NaCl concentrations and conductivity after reaching swelling equilibrium in PBS.

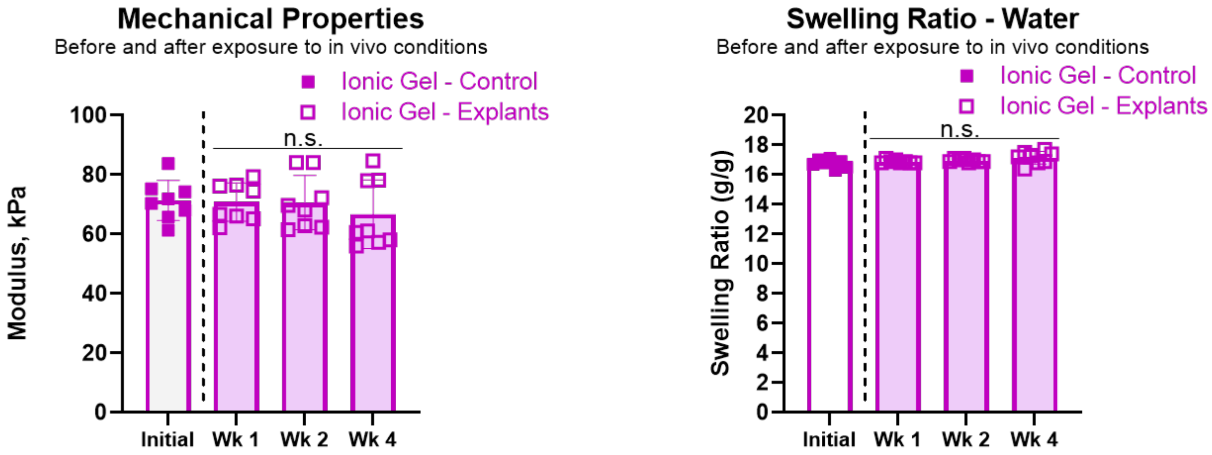


**Figure S14. Network properties of PEUDAm prior to and after subcutaneous implantation in rats for 1, 2 or 4 weeks (n=8).**

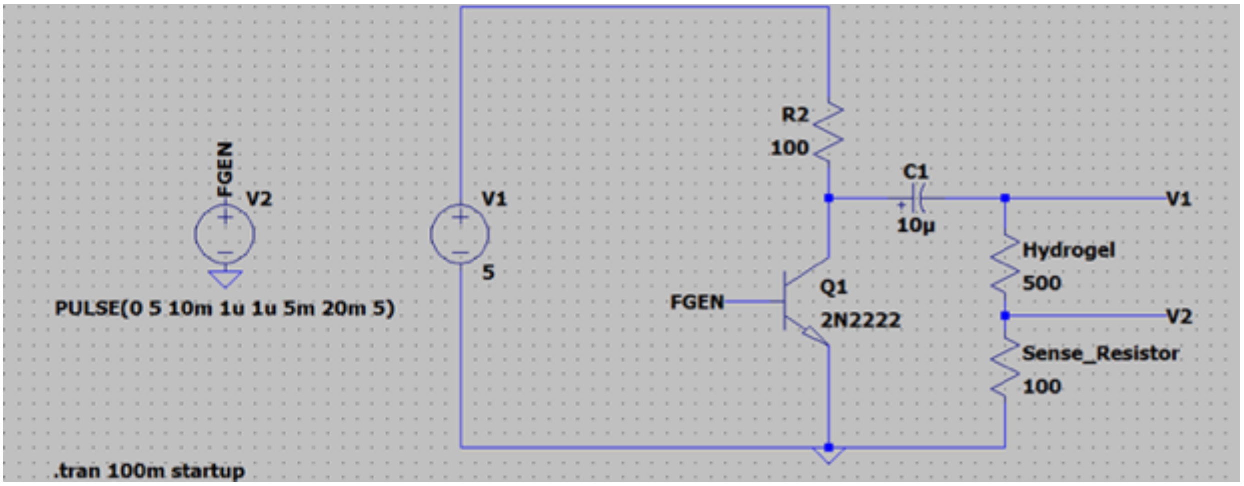


**Figure S15. Circuit used to test the impact of hydrogel length on current.** An oscilloscope measured the voltage across the sense resistor to determine the current.

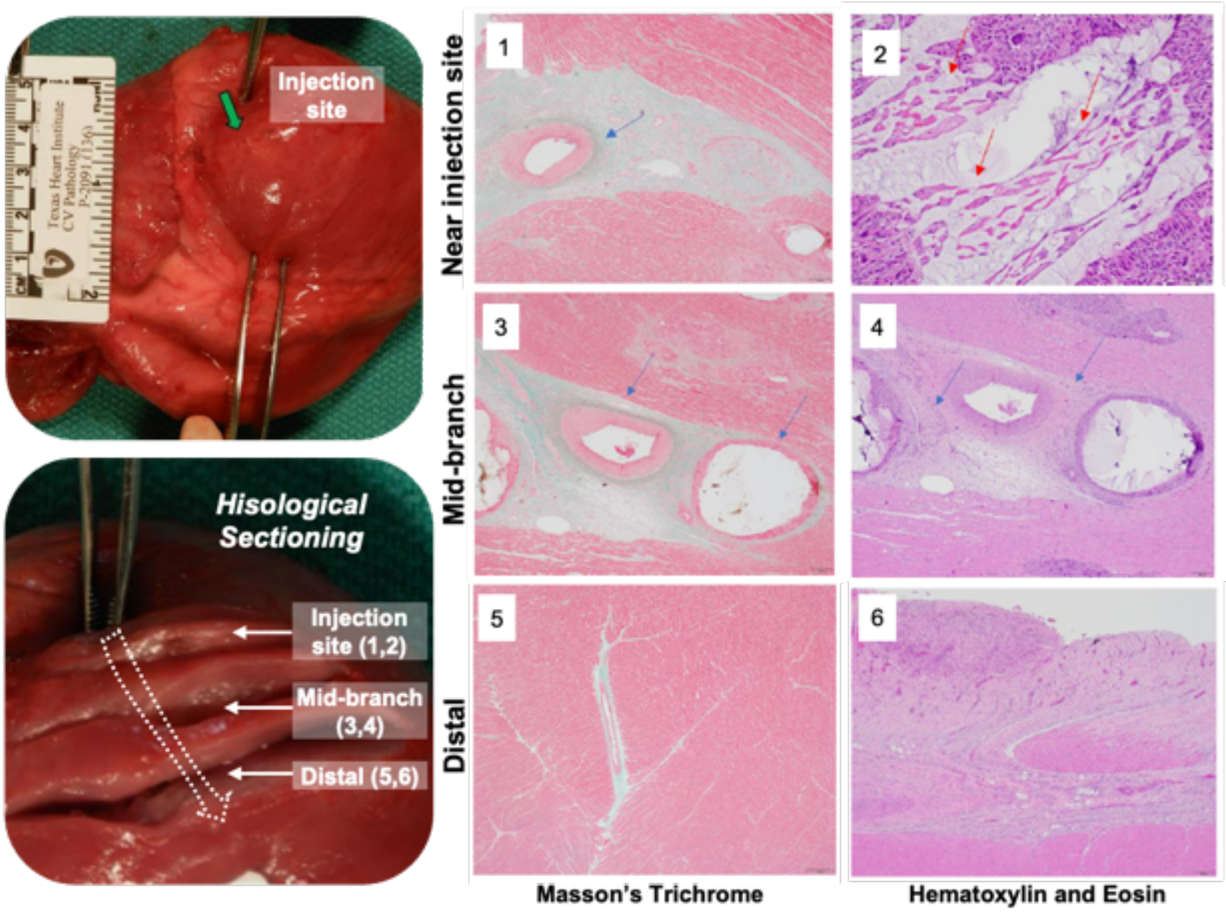


**Figure S16. *In vivo* assessment of injectable electrode in coronary vein (middle cardiac vein) of a porcine model.** Host response after 2 weeks implantation in MCV using slices of the location proximal to the injection site, middle of the vein, and distal to the injection site (green arrow). The white arrow indicates the approximate location of the hydrogel in the vein. The response includes damage induced at the hydrogel injection site, with 1) moderate perivascular and interstitial fibrosis with focal replacement fibrosis (blue arrows), and 2) fibrosing epicarditis with foreign body giant cell reaction (red arrows). Mid branch of the hydrogel injection show 3) mild perivascular and interstitial fibrosis, 4) mild focal replacement fibrosis, and fibrosing epicarditis. Distal branch indicated 5) preserved myocardium with 6) only fibrosing epicarditis with focal extension into myocardium.

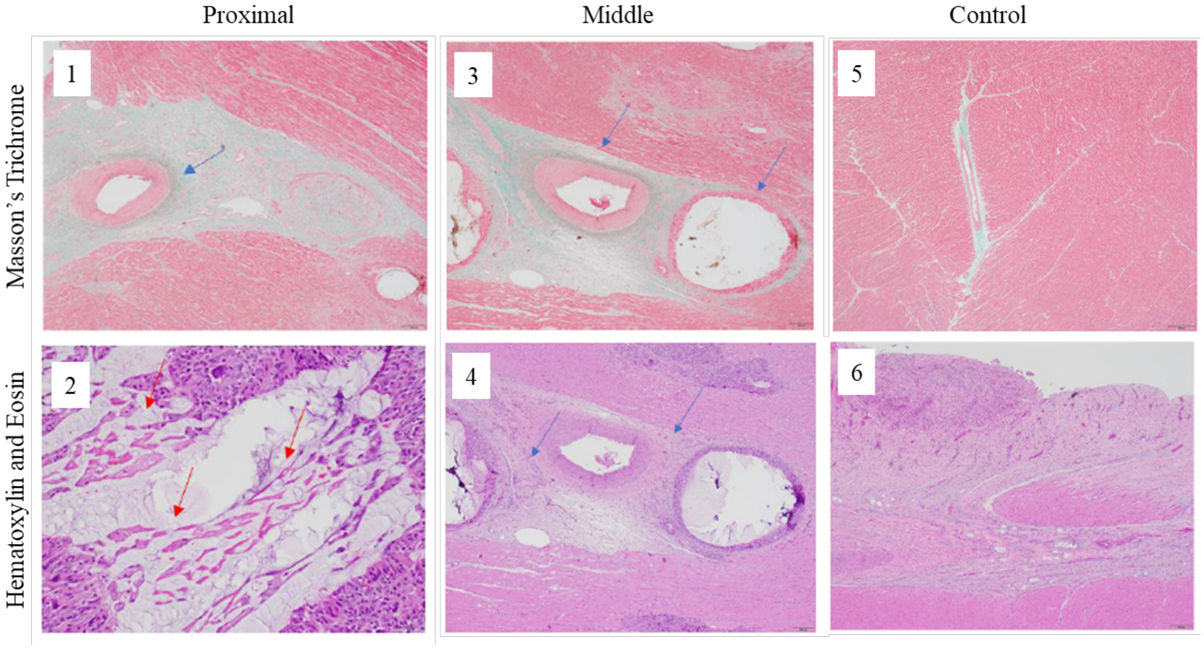


**Figure S17. *In vivo* assessment of injectable electrode in coronary vein (anterior interventricular vein) of a porcine model (Animal 1).** Host response after 4 weeks implantation in AIV using slices of the location proximal to the injection site, middle of the vein (indicated by V), and the control that is distal to the hydrogel. The response includes damage induced at the hydrogel injection site, with 1) moderate perivascular and interstitial fibrosis with focal replacement fibrosis (blue arrows), and 2) fibrosing epicarditis (red arrows) with foreign body giant cell reaction (blue arrows). Mid branch of the hydrogel injection show 3) mild perivascular and interstitial fibrosis, 4) mild focal replacement fibrosis, and fibrosing epicarditis. The control images at the distal branch indicated 5) preserved myocardium with 6) only fibrosing epicarditis with focal extension into myocardium.

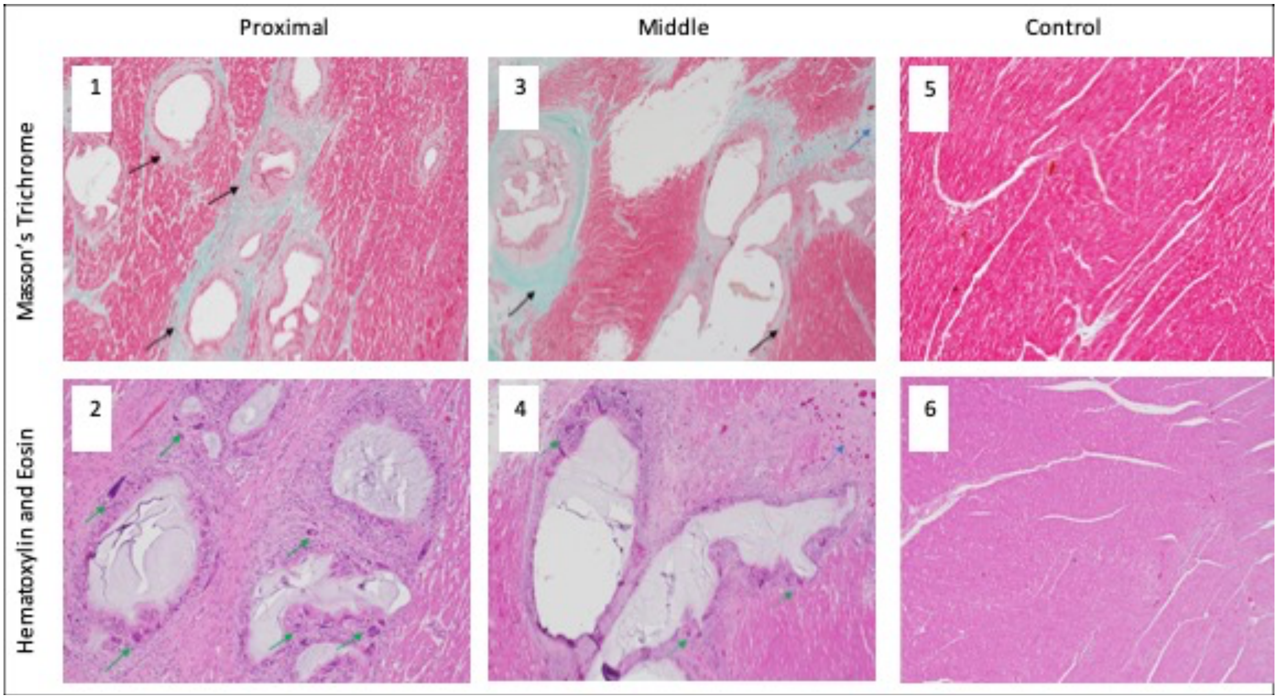


**Figure S18.** ***In vivo*** **assessment of injectable electrode in coronary vein (anterior interventricular vein) of a porcine model (Animal 2).** Host response after 4 weeks implantation in AIV using slices of the location proximal to the injection site, middle of the vein (indicated by V), and the control from an alternate section of the anterior wall. The response includes damage induced at the hydrogel injection site, with 1) moderate perivascular and interstitial fibrosis with focal replacement fibrosis (blue arrows), and 2) fibrosing epicarditis with foreign body giant cell reaction (green arrows). Mid branch of the hydrogel injection show 3) moderate perivascular and interstitial (black arrows) fibrosis, and replacement fibrosis (blue arrows), 4) focal replacement fibrosis, and fibrosing epicarditis. The control images at an alternate section of the anterior wall indicated 5) 6) preserved myocardium

**A.** Time matched ECG tracings. **B.** QRS and vector morphology matched ECG tracings.

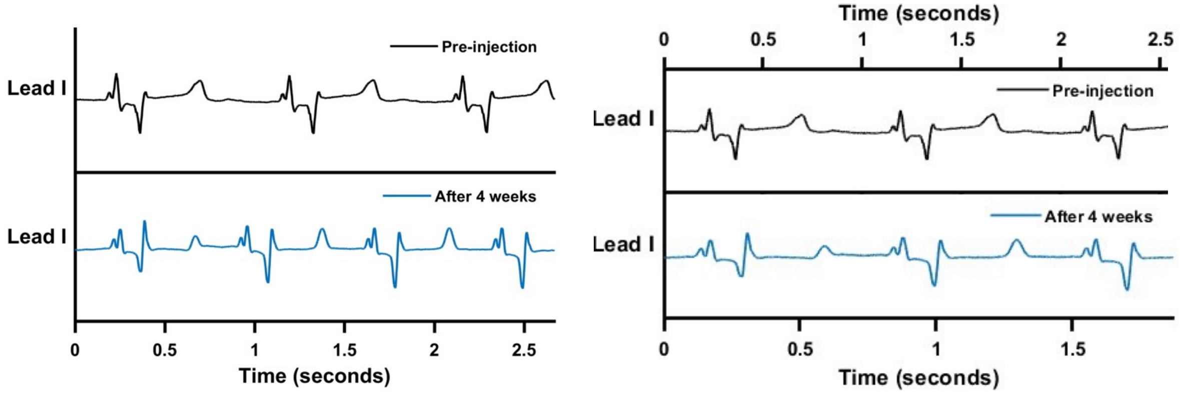


**Figure S19. *In vivo* assessment of injectable electrode in coronary vein (anterior intraventricular vein) of a porcine model.** Animal AIV 1. ECG tracings from lead 1 of the cardiac activity before injection of the hydrogel (black tracing) and 4 weeks after injection (blue tracing). The overall QRS morphology is preserved. Heart rate differs between the two recordings, but this is not indicative of a disease state.

**A.** Time matched ECG tracings with same heart rate.

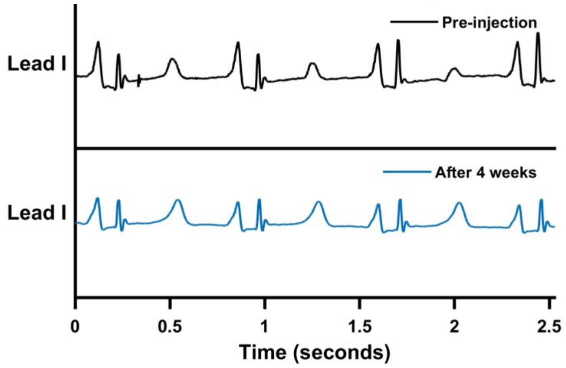


**Figure S20. *In vivo* assessment of injectable electrode in coronary vein (anterior intraventricular vein) of a porcine model.** Animal AIV 2. ECG tracings from lead 1 of the cardiac activity before injection of the hydrogel (black tracing) and 4 weeks after injection (blue tracing). The overall QRS morphology is preserved.

**A.** Time matched ECG tracings.

**B.** QRS and vector morphology matched ECG tracings.

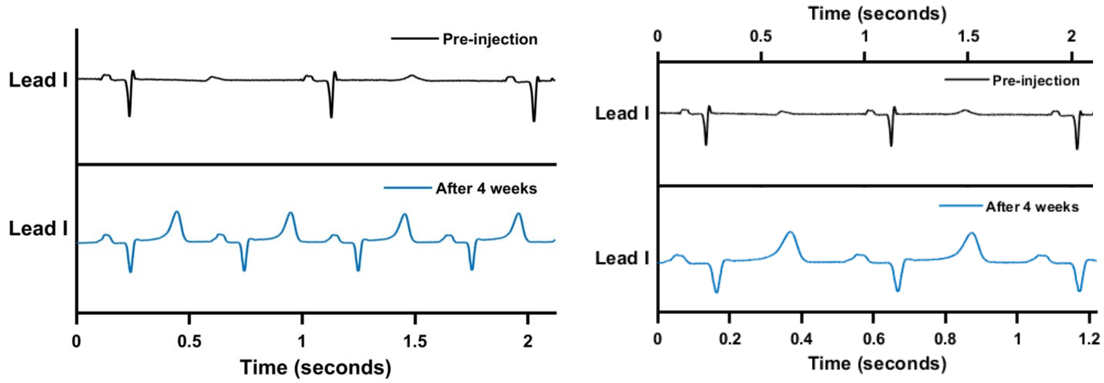


**Figure S21.** ***In vivo*** **assessment of injectable electrode in coronary vein (anterior intraventricular vein) of a porcine model.** Animal AIV 3. ECG tracings from lead 1 of the cardiac activity before injection of the hydrogel (black tracing) and 4 weeks after injection (blue tracing). The overall QRS morphology is similar, with nearly identical vectors. Heart rate differs between the two recordings, but this is not indicative of a disease state.

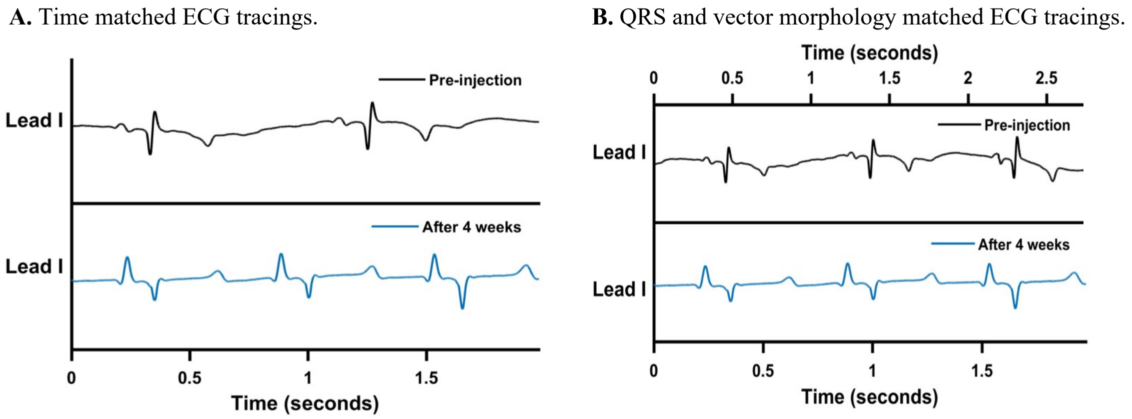


**Figure S22.** ***In vivo* assessment of injectable electrode in coronary vein (middle cardiac vein) of a porcine model.** Animal MCV 1. ECG tracings from lead 1 of the cardiac activity before injection of the hydrogel (black tracing) and 2 weeks after injection (blue tracing). The overall QRS morphology is similar, with nearly identical vectors. Heart rate differs between the two recordings, but this is not indicative of a disease state. The pre-injection ECG tracing contains low frequency baseline drift which is not reflective of the model's electrophysiology.

**Table S4. High sensitivity Troponin I measurements.** Measurements from four separate studies from two weeks after occlusion of the MCV and from four weeks after occlusion of the AIV in a porcine model. The normal range for troponin I is from 3 to 70 ng.

| Vein - Animal | Troponin levels (ng/L) | | | | | |
|---|---|---|---|---|---|---|
| | **Pre-Op** | **Post-Op** | **Week 1** | **Week 2** | **Week 3** | **Week 4** |
| **MCV - 1** | -- | 3313 | -- | 34 | -- | -- |
| **AIV - 1** | 17 | 421 | 99 | 97 | 99 | 49 |
| **AIV - 2** | 121 | N/A | 202 | 95 | 116 | 54 |
| **AIV - 3** | 17 | 140 | 44 | 28 | 62 | 45 |

**2 weeks after injection on MCV (EF = 65%)**

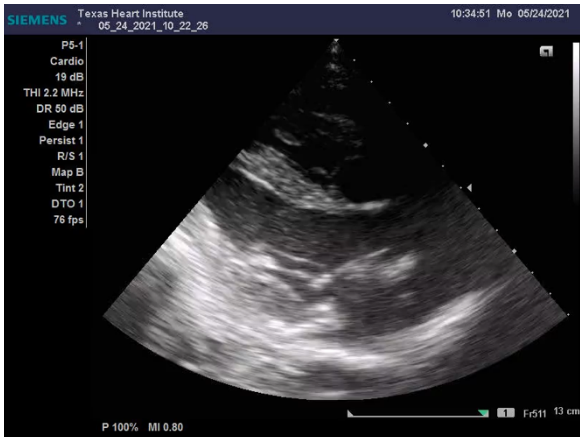


**Movie S3. Echocardiogram 2 weeks after injection on the MCV in a pig model.** Echocardiograms show no evidence of regional wall motion abnormalities after two weeks. Press play to see the videos.

**Baseline (Prior hydrogel injection) (EF = 70%).** **4 weeks after injection on AIV (EF = 65%)**

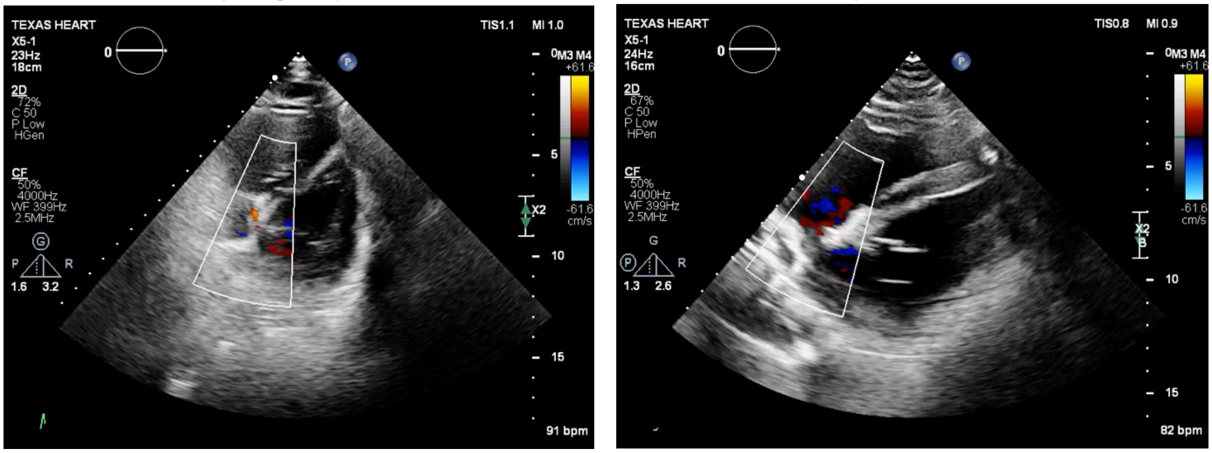


**Movie S4. Echocardiogram prior and 4 weeks after injection on the AIV in a pig model.** Echocardiograms show no evidence of regional wall motion abnormalities at four weeks compared to baseline imaging. Press play to see the videos.

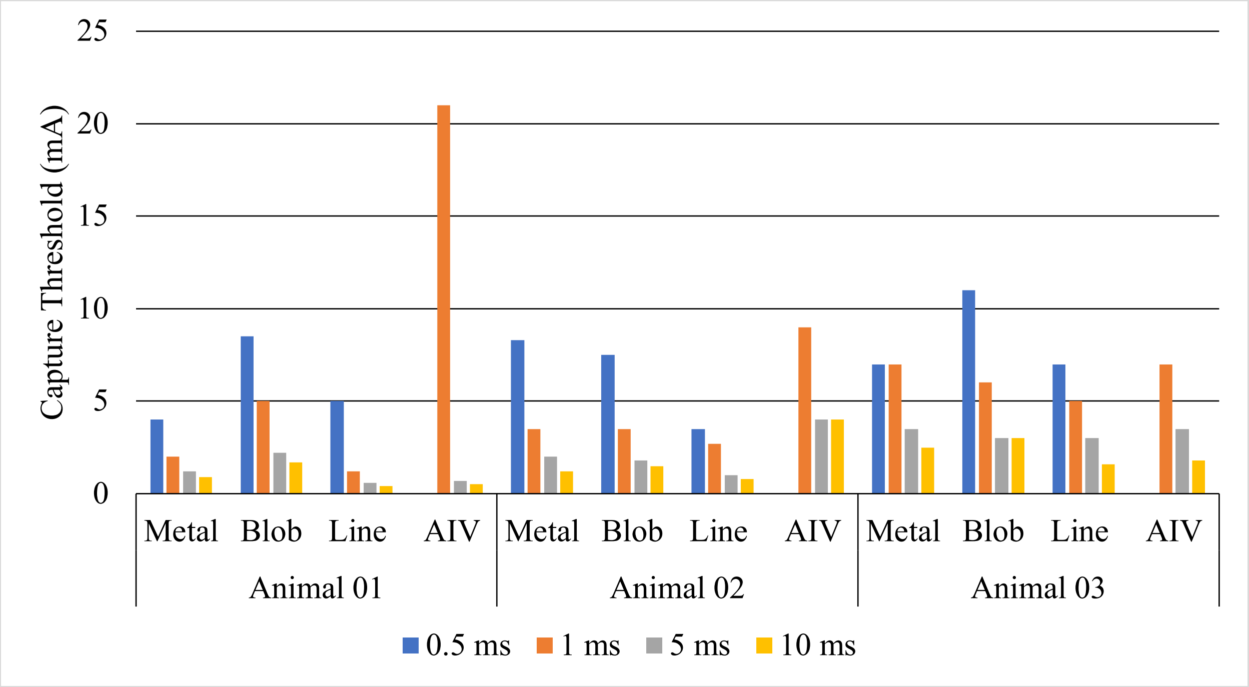


**Figure S23. Capture thresholds for each acute porcine study (n=3).**

**Table S5. Capture thresholds (in mA) for each material and acute animal tested (n=3).** NC = no capture.

| **PW (ms)** | **Animal 01** | | | | **Animal 02** | | | | **Animal 03** | | | |
|---|---|---|---|---|---|---|---|---|---|---|---|---|
| | **Metal** | **Blob** | **Line** | **AIV** | **Metal** | **Blob** | **Line** | **AIV** | **Metal** | **Blob** | **Line** | **AIV** |
| 0.5 | 4 | 8.5 | 5 | - | 8.3 | 7.5 | 3.5 | - | 7 | 11 | 7 | - |
| 1 | 2 | 5 | 1.2 | 21 | 3.5 | 3.5 | 2.7 | 9 | 7 | 6 | 5 | 7 |
| 5 | 1.2 | 2.2 | 0.6 | 0.7 | 2 | 1.8 | 1 | 4 | 3.5 | 3 | 3 | 3.5 |
| 10 | 0.9 | 1.7 | 0.4 | 0.5 | 1.2 | 1.5 | 0.8 | 4 | 2.5 | 3 | 1.6 | 1.8 |

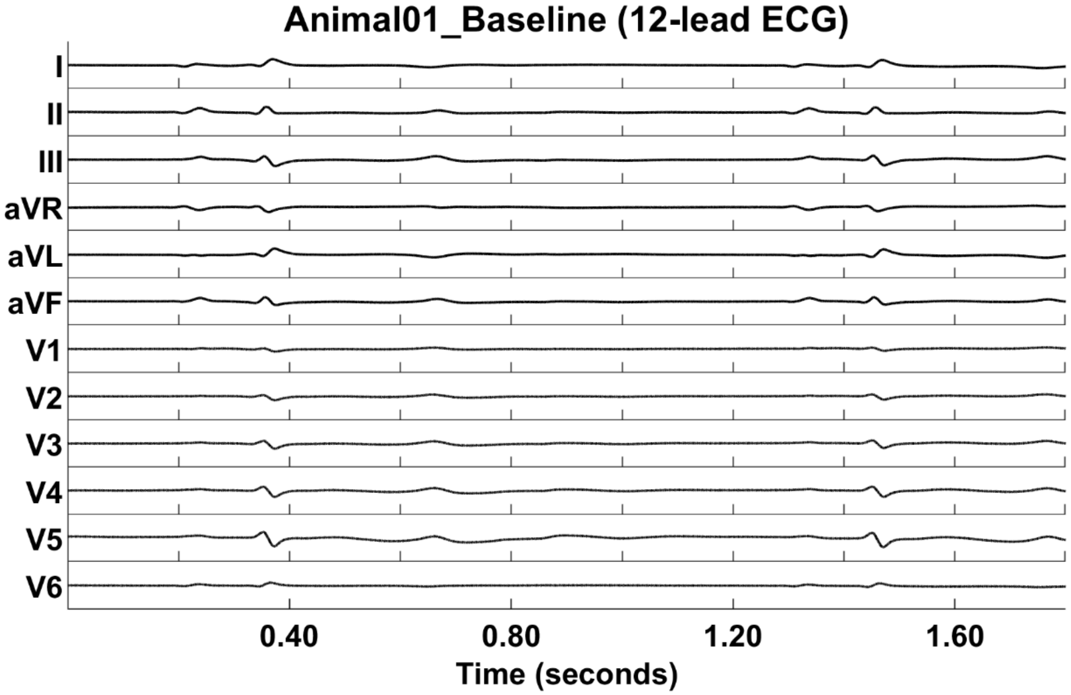

**Figure S24. 12-lead ECG of the heart at sinus rhythm.**

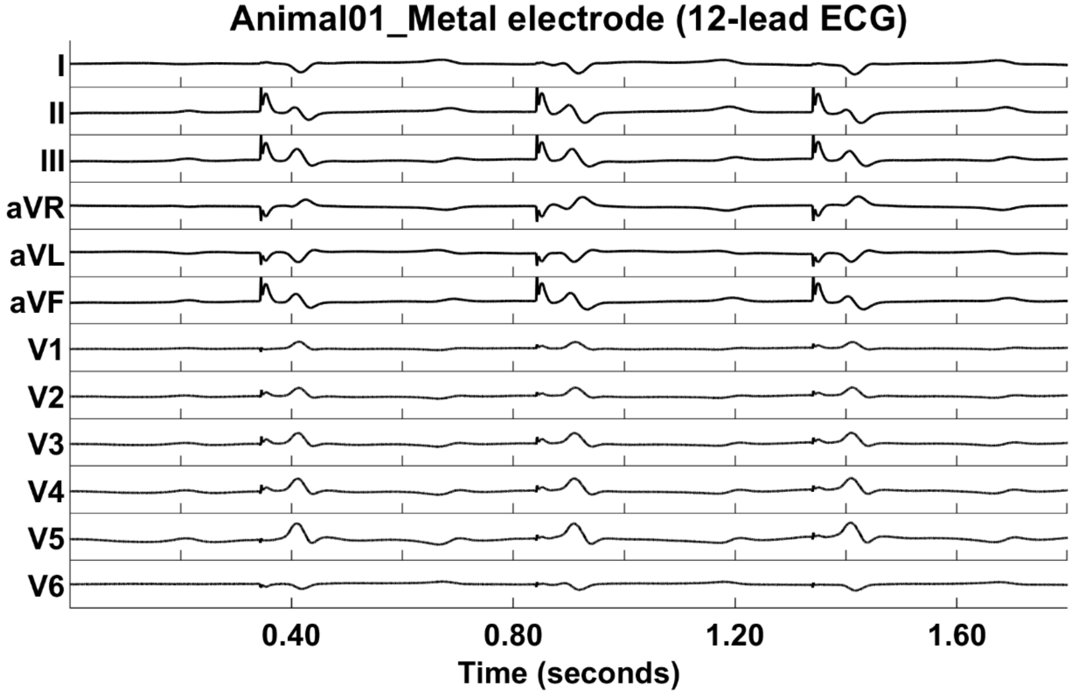


**Figure S25. 12-lead ECG of unipolar pacing with a metal electrode placed directly on the left ventricular myocardium**. Pacing artifacts can be seen as spikes in each of the channels. Pacing was performed at 120 bpm.

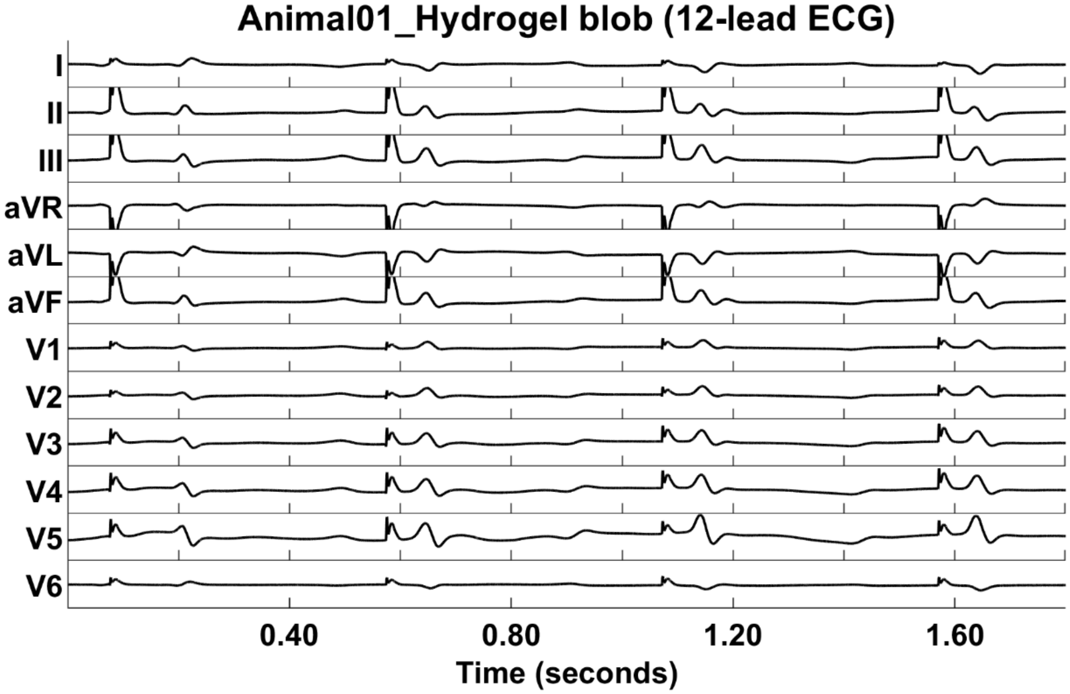


**Figure S26. 12-lead ECG of unipolar pacing with a small blob of hydrogel electrode placed directly on the left ventricular myocardium.** Pacing artifacts can be seen as spikes in each of the channels. Pacing was performed at 120 bpm.

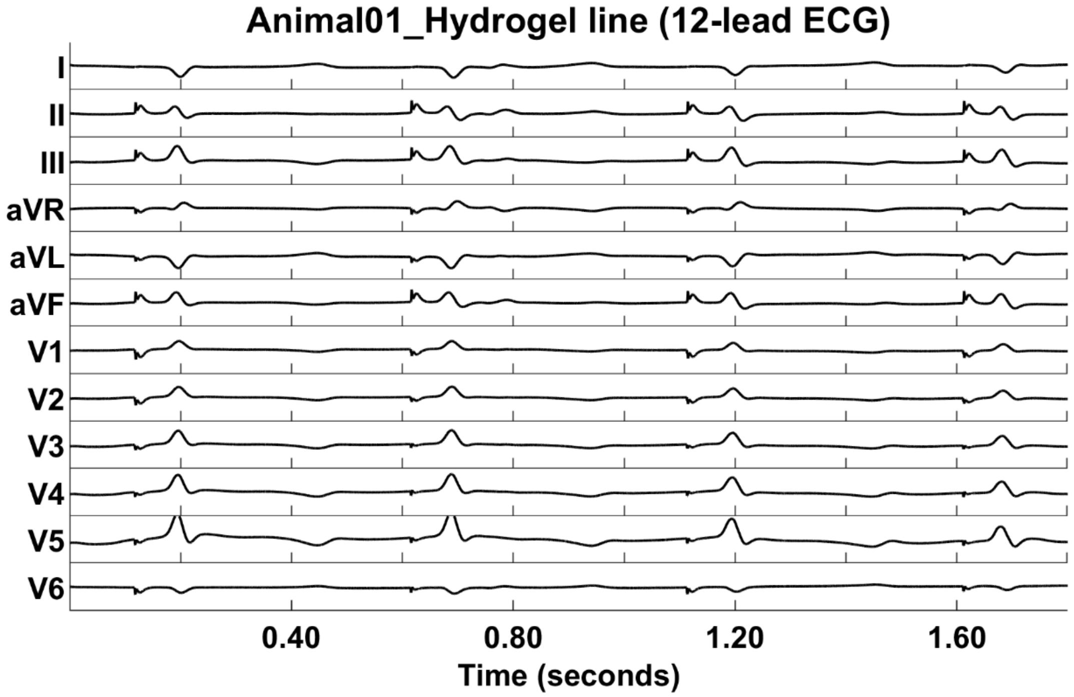


**Figure S27. 12-lead ECG of unipolar pacing with a linear hydrogel electrode (~4 cm) placed directly on the left ventricular myocardium close to the interventricular septum.** Pacing artifacts can be seen as spikes in each of the channels. Pacing was performed at 120 bpm.

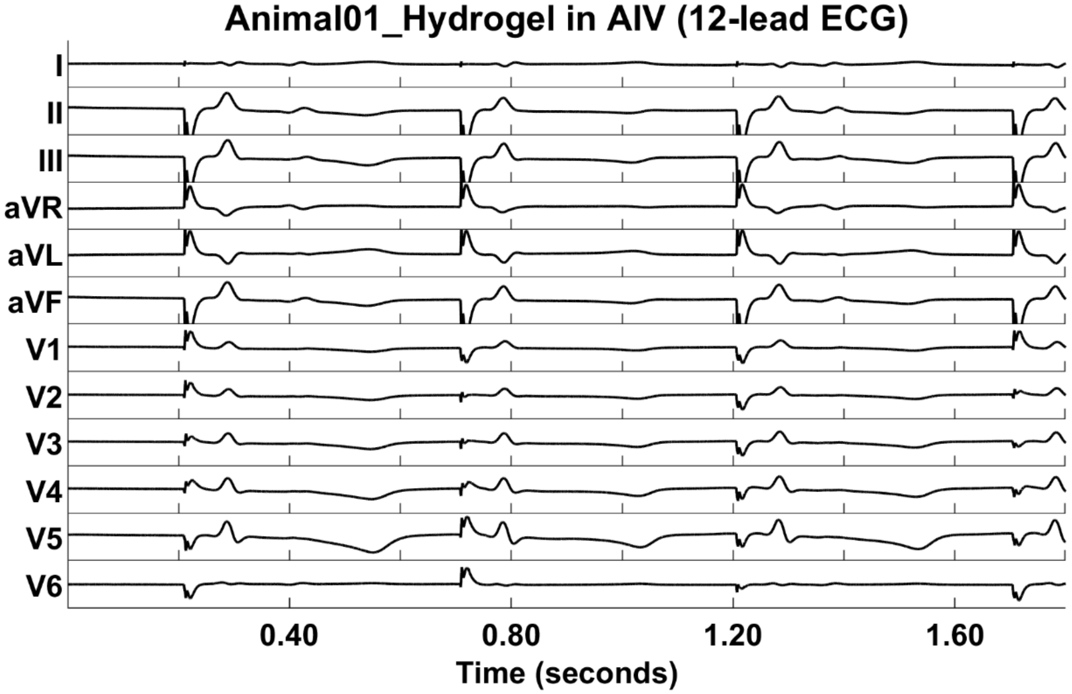


**Figure S28. 12-lead ECG of unipolar pacing from hydrogel cured in situ in the AIV. Pacing artifacts can be seen as spikes in each of the channels.** Pacing was performed at 120 bpm.

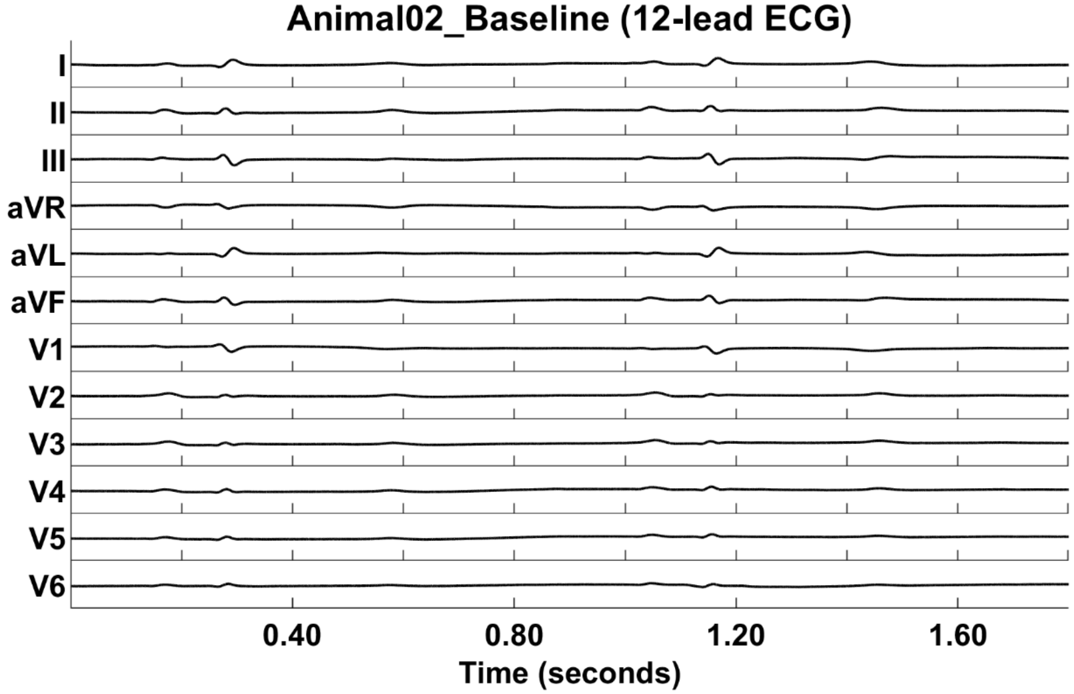


**Figure S29. Animal02: 12-lead ECG of the heart at sinus rhythm.**

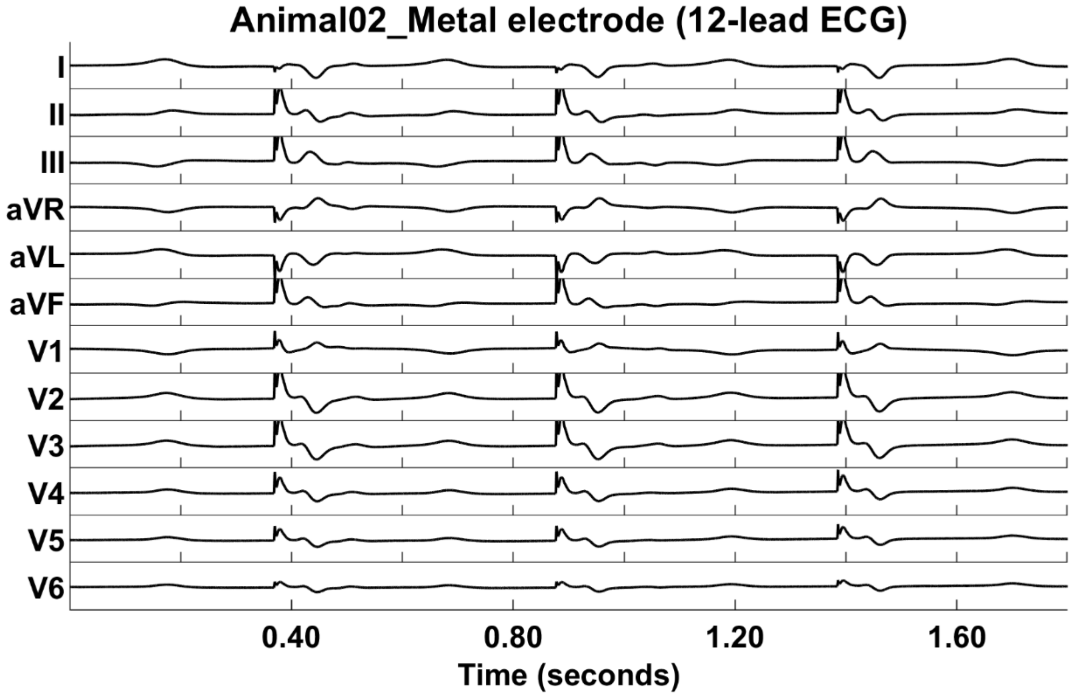


**Figure S30. Animal02: 12-lead ECG of unipolar pacing with a metal electrode placed directly on the left ventricular myocardium.** Pacing artifacts can be seen as spikes in each of the channels. Pacing was performed at 120 bpm.

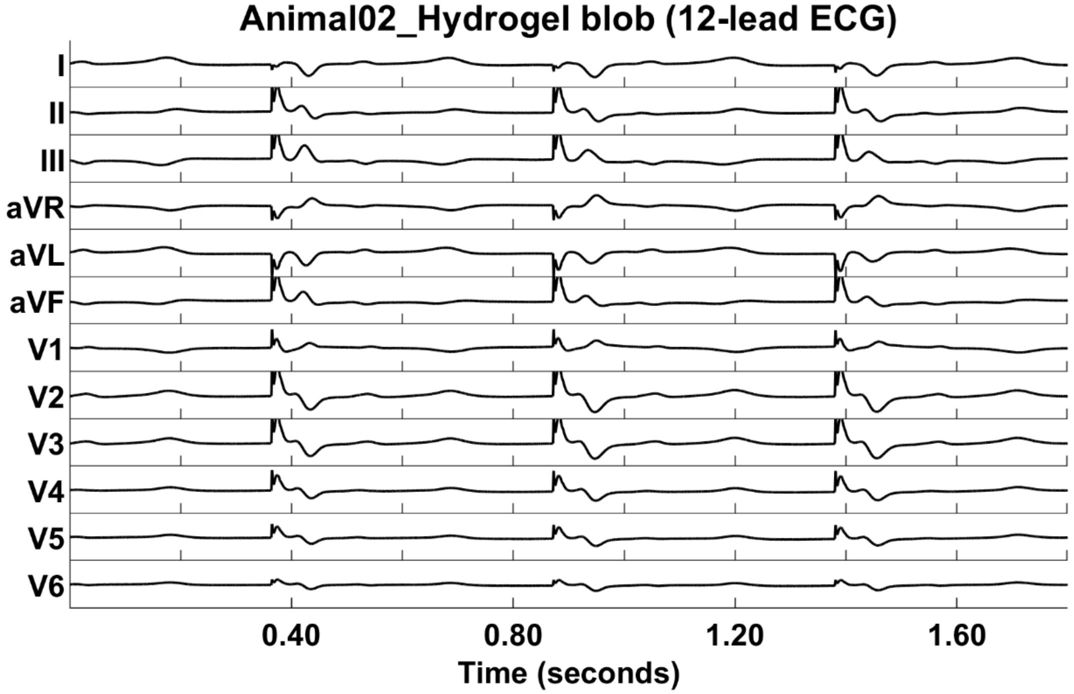


**Figure S31. Animal02: 12-lead ECG of unipolar pacing with a small blob of hydrogel electrode placed directly on the left ventricular myocardium.** Pacing artifacts can be seen as spikes in each of the channels. Pacing was performed at 120 bpm.

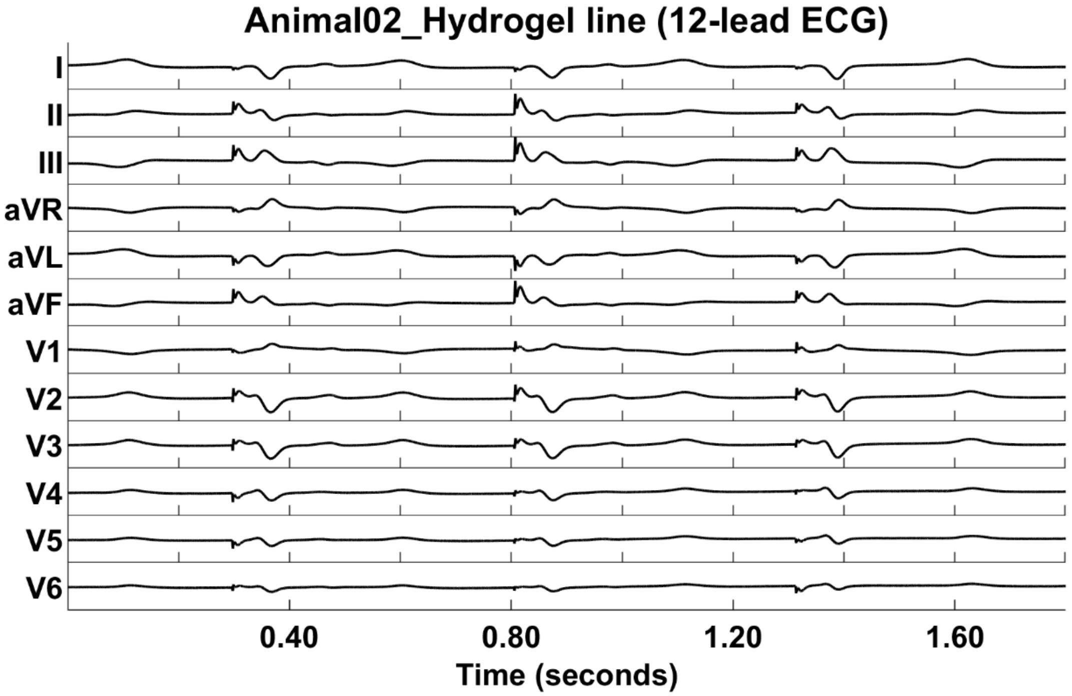


**Figure S32. Animal02: 12-lead ECG of unipolar pacing with a linear hydrogel electrode (~4 cm) placed directly on the left ventricular myocardium close to the interventricular septum.** Pacing artifacts can be seen as spikes in each of the channels. Pacing was performed at 120 bpm.

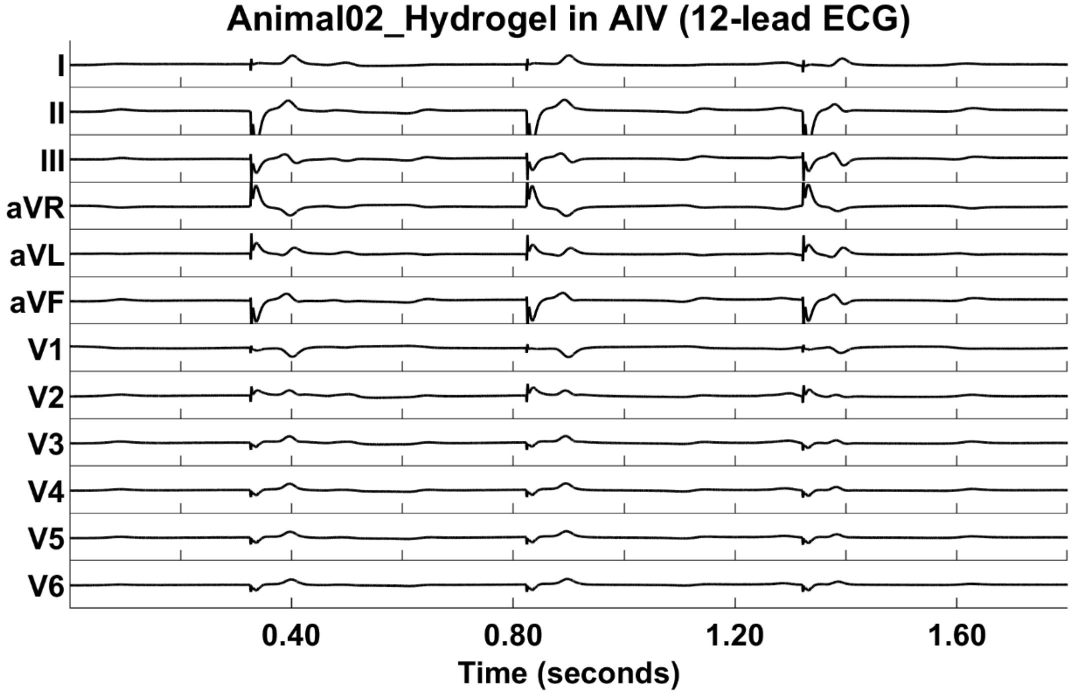


**Figure S33. Animal02: 12-lead ECG of unipolar pacing from hydrogel cured in situ in the AIV.** Pacing artifacts can be seen as spikes in each of the channels. Pacing was performed at 120 bpm.

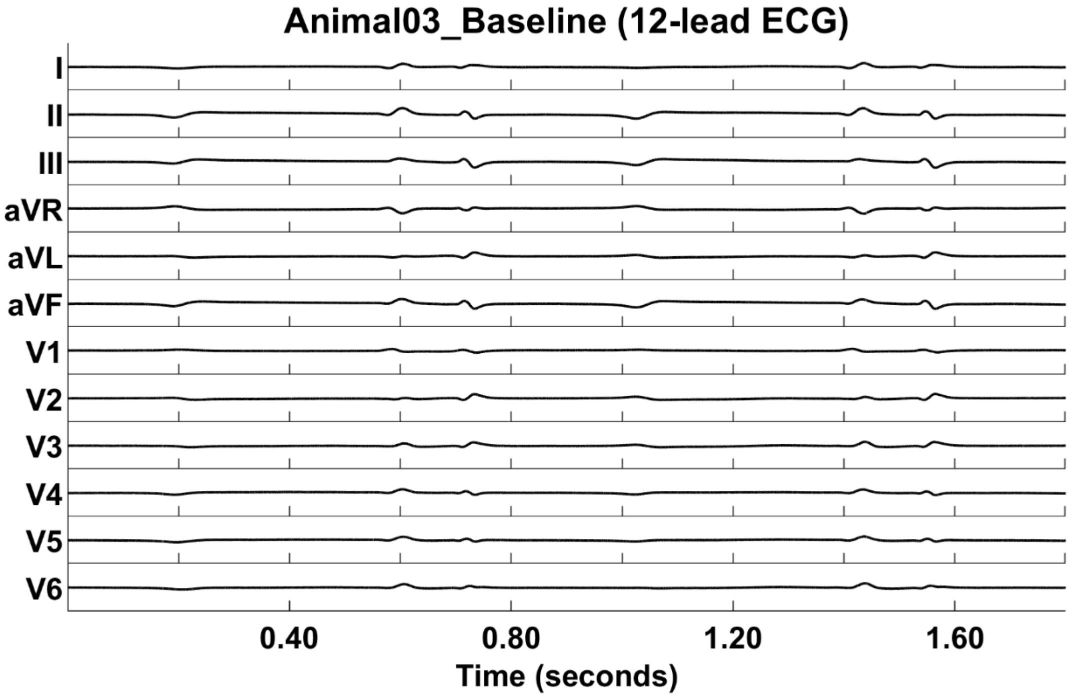


**Figure S34. Animal03: 12-lead ECG of the heart at sinus rhythm.**

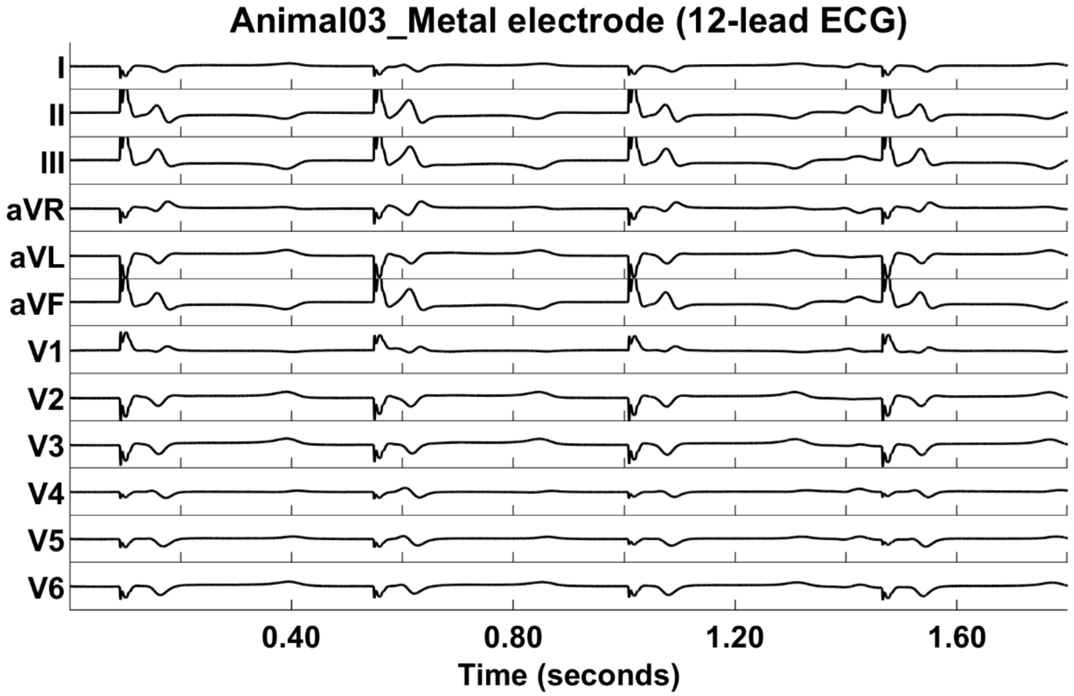


**Figure S 35. Animal03: 12-lead ECG of unipolar pacing with metal electrode placed directly on the left ventricular myocardium.** Pacing artifacts can be seen as spikes in each of the channels. Pacing was performed at 120 bpm.

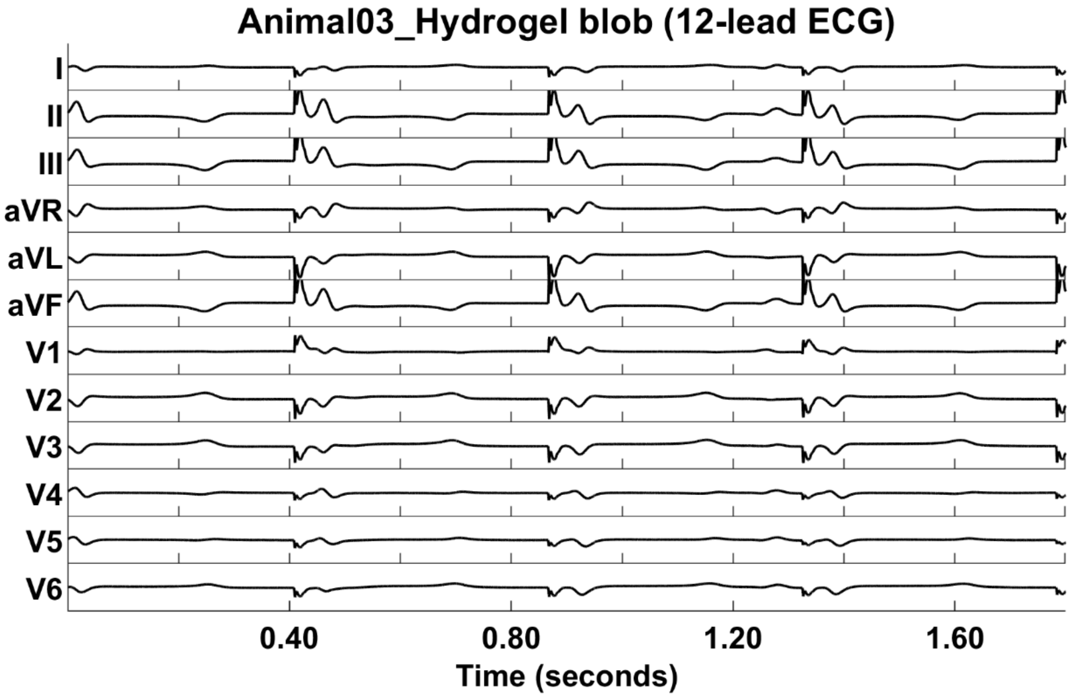


**Figure S36. Animal03: 12-lead ECG of unipolar pacing with a small blob of hydrogel electrode placed directly on the left ventricular myocardium.** Pacing artifacts can be seen as spikes in each of the channels. Pacing was performed at 120 bpm.

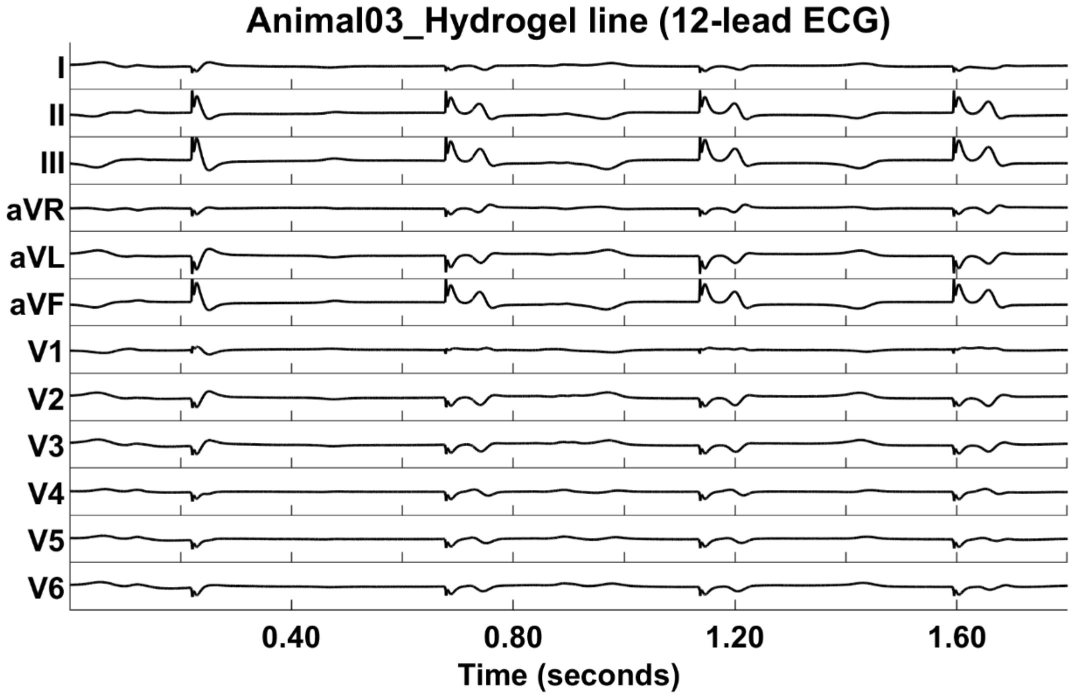


**Figure S37. Animal03: 12-lead ECG of unipolar pacing with a linear hydrogel electrode (~4 cm) placed directly on the left ventricular myocardium close to the interventricular septum.** Pacing artifacts can be seen as spikes in each of the channels. Pacing was performed at 120 bpm.

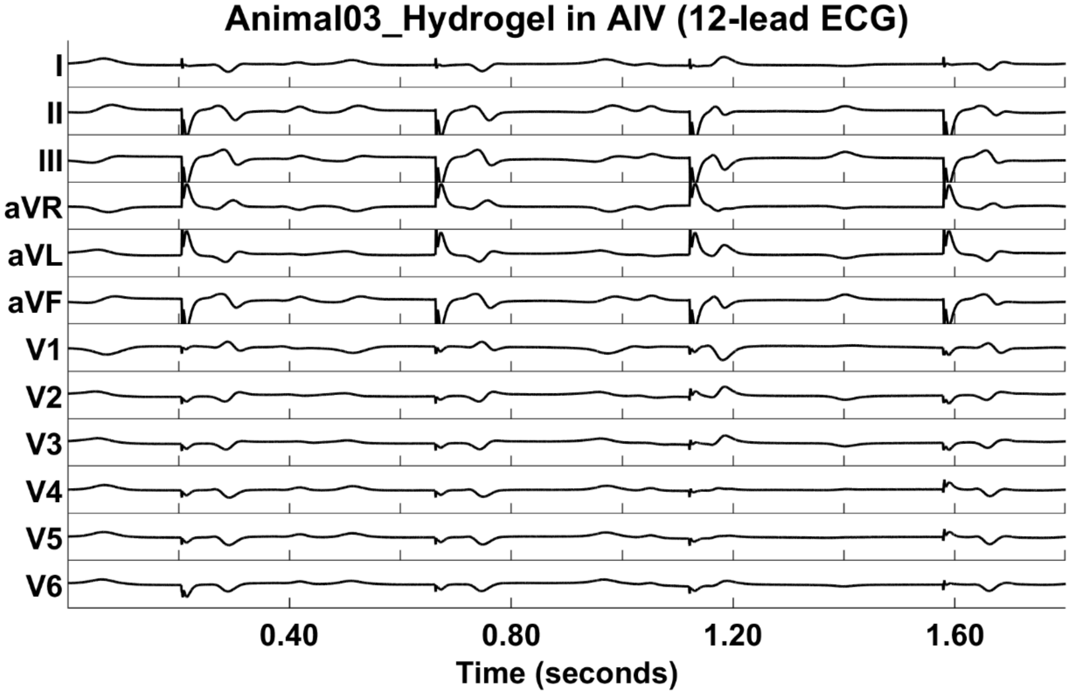


**Figure S38. Animal03: 12-lead ECG of unipolar pacing from hydrogel cured in situ in the AIV. Pacing artifacts can be seen as spikes in each of the channels.** Pacing was performed at 120 bpm.

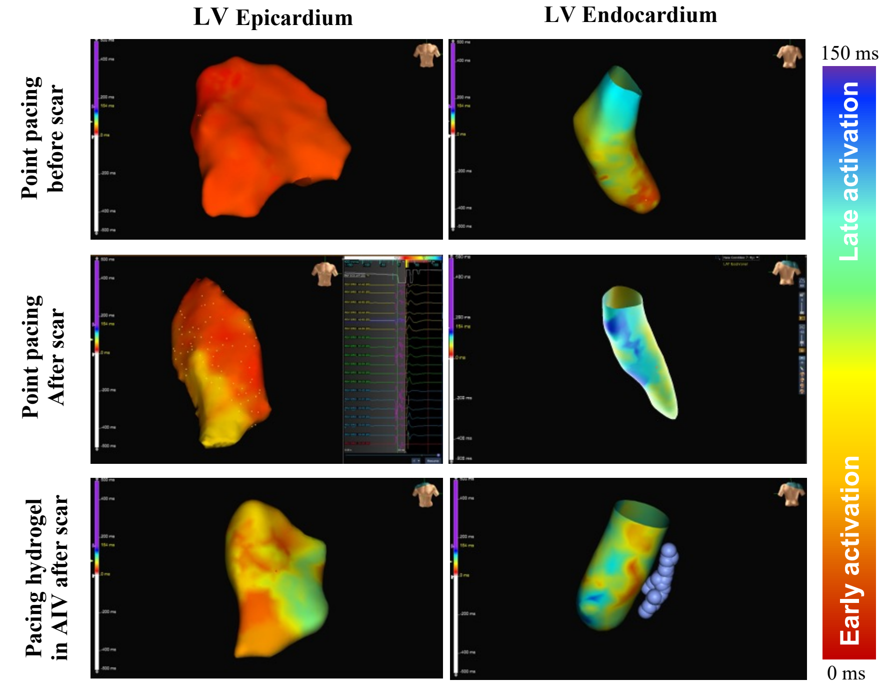


**Figure S39. Epicardial and endocardial electroanatomical mapping for animal AIV 1.** The AIV hydrogel pacing is indicated by the blue circles.

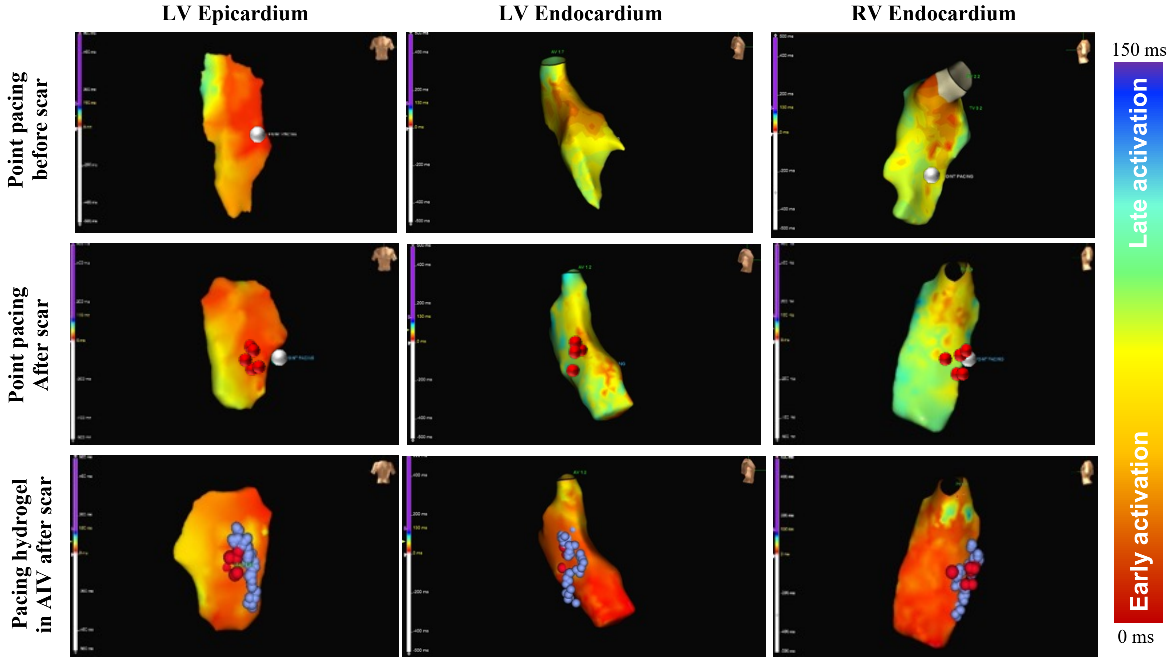


**Figure S40. Epicardial and endocardial electroanatomical mapping for animal AIV 2.** Point pacing location is indicated by a small white circle, and the AIV hydrogel pacing is indicated by the blue circles. Points of ablation are marked by the red circles.

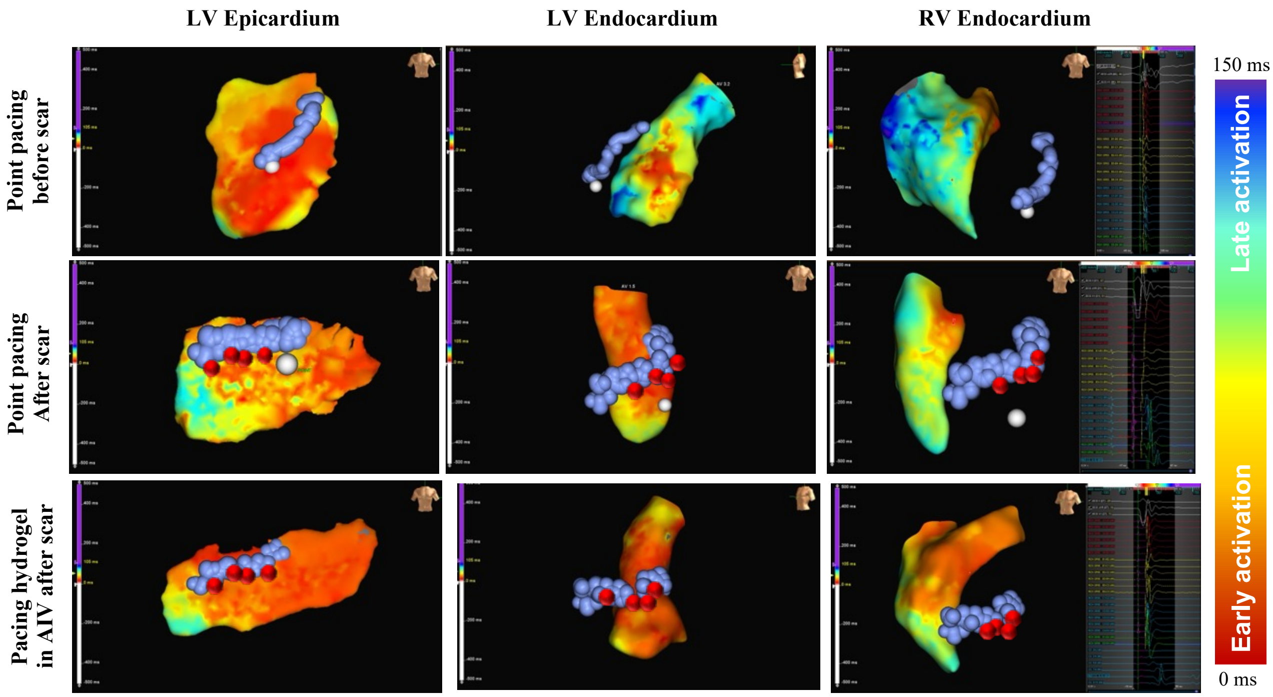


**Figure S41. Epicardial and endocardial electroanatomical mapping for animal AIV 3.** Point pacing location is indicated by a small white circle, and the AIV is indicated by the blue circles. Points of ablation are marked by the red circles

**Point pacing** | **Hydrogel (AIV) pacing**

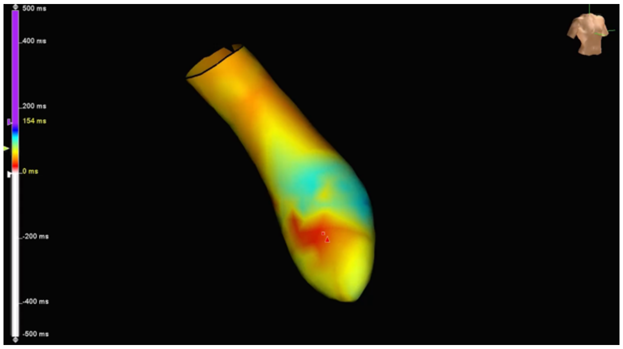


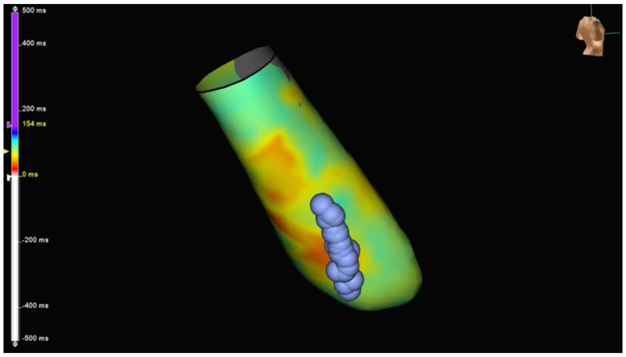


**Movie S5. Myocardium activation comparison after pacing with a single point (metal electrode) or with the hydrogel in the vein for Animal AIV 1**. The AIV is indicated by the blue circles. Press play to see the animations.

**Point pacing**

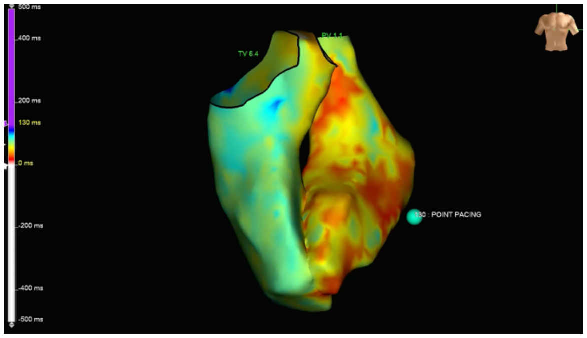


**Hydrogel (AIV) pacing**

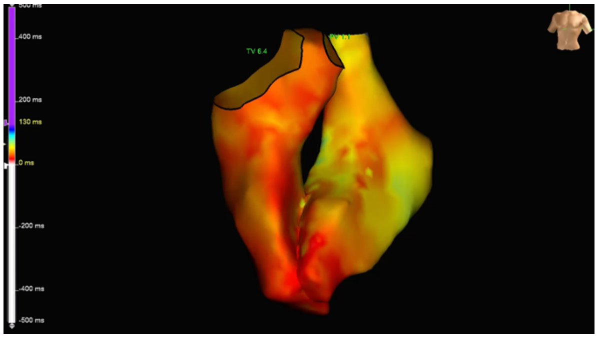


**Movie S6. Myocardium activation comparison after pacing with a single point (metal electrode) or with the hydrogel in the vein for Animal AIV 2**. A small green circle indicates point pacing location on the first image. Press play to see the animations.
